# Satellite DNA fragility accompanies complex genome rearrangements and ecDNA oncogene amplification in canine osteosarcomas

**DOI:** 10.64898/2025.12.31.697157

**Authors:** Feyza Yilmaz, Wonyoung Kong, Sabriya A. Syed, Francis H. O’Neill, Jody T. Lombardi, Patrick Kwok Shing Ng, Ching C. Lau, Charles Lee

**Affiliations:** - The Jackson Laboratory for Genomic Medicine, CT, 06032, USA; - University of Connecticut School of Medicine, Farmington, CT, 06030, USA; - Connecticut Children’s Medical Center, Hartford, CT, 06106, USA

**Keywords:** Canine Osteosarcoma, Structural Variation, Satellite Repeats, Extrachromosomal DNA, Long-Read Sequencing

## Abstract

Canine osteosarcoma (OS) is an aggressive bone cancer with remarkable similarity to human OS. Despite its value as a comparative oncology model, the drivers of genomic instability remain poorly defined, hindering cross-species insights into disease progression. We applied long-read PacBio HiFi whole genome sequencing to matched tumor and blood samples from four purebred dogs to generate high-resolution maps of somatic structural variants (SVs). Tumor genomes showed extensive rearrangement complexity, with satellite repeat regions, particularly SAT1_CF, enriched near somatic SV breakpoints and occurring within genomic contexts marked by focal hypomethylation. We also identified multiple extrachromosomal DNA (ecDNA) elements carrying oncogenes indicating ecDNA-mediated oncogene amplification. Together, these data provide an integrated genomic and epigenomic view of canine OS, implicating satellite repeats as recurrent substrates for rearrangement and ecDNA as a prominent mode of genome remodeling. This work advances our understanding of OS genome dynamics and establishes a comparative framework for investigating repeat-driven instability and oncogene amplification across species.

## Introduction

Osteosarcoma (OS) is a rare but devastating bone cancer in humans, occurring primarily in children and adolescents with an incidence of only ∼4-5 cases per million (1). Despite advances in treatment, outcomes for patients with metastatic or relapsed disease remain poor with five-year survival rates below 30% (2, 3), highlighting an urgent need for deeper biological insight and improved therapeutic strategies. Among canines, OS is a common and highly aggressive cancer found particularly in large- and giant-breed dogs and occurring at an incidence 27-fold higher than in humans (4). Importantly, spontaneous canine OS closely recapitulates key clinical and molecular features of human OS, including anatomic distribution, metastatic behavior, and histopathology, making dogs a powerful model for comparative oncology and translational discovery (5–7). Cross-species genomic and transcriptomic studies increasingly demonstrate conserved tumor biology and clinical outcomes, underscoring the value of canine OS for biomarker discovery and therapeutic development relevant to human disease (8).

At the genomic level, both canine and human OS share critical molecular hallmarks. Both exhibit relatively low somatic point mutation burdens compared to most adult cancers (1.2 mutations/Mb), yet higher mutation rates than other pediatric malignancies (such as neuroblastoma and medulloblastoma; ∼0.4-0.5 mutations/Mb) (9, 10). Moreover, OS genomes are dominated by extensive structural variation (SV) and copy-number alterations (9, 11–14). High-grade human OS tumors frequently display chromothripsis and other complex, clustered rearrangements that profoundly remodel chromosomal architecture. Comparative analyses of canine tumors and cell lines reveal similarly chaotic karyotypes, suggesting that large scale structural genome instability is a conserved and fundamental feature of OS across species (12, 15, 16).

Growing evidence from cancer genomics and epigenetics implicates repetitive DNA, especially pericentromeric satellite repeats, as key substrates for genome instability (17, 18). These tandemly repeated elements are concentrated in heterochromatic regions flanking centromeres, where loss of DNA methylation has been linked to heterochromatin decondensation, replication stress, and elevated rates of chromosomal instability (18–20). Together, these observations position satellite repeats as potential hotspots for SV formation, yet their role in OS genome remodeling remains poorly defined.

Another recurrent feature of aggressive solid tumors, including OS, is focal oncogene amplification on extrachromosomal DNA (ecDNA; historically referred to as “double minutes”) (12, 21–23). Pan-cancer studies demonstrate that ecDNA is common, preferentially harbors oncogenes and regulatory elements, drives intratumoral heterogeneity, and is associated with poor clinical outcomes (21). In human bone tumors, recurrent amplicons (*e.g.*, 12q13–15/*MDM2–CDK4*) are frequently localized to extrachromosomal or ring structures (24, 25), implicating ecDNA as an important mechanism of oncogene amplification in this disease (21, 22, 26). While extrachromosomal DNA (ecDNA) has emerged as a major driver of oncogene amplification and treatment resistance across many human cancers, its prevalence and functional significance in human OS remain incompletely characterized (12, 26). Similarly, the role of ecDNA in canine OS is unknown.

Here, we apply PacBio HiFi whole-genome sequencing to primary tumors and matched blood samples (including normal stroma) from four purebred dogs with OS to investigate the mechanisms of structural genome instability. By jointly analyzing SVs, repetitive DNAs, native CpG methylation and ecDNA, we observed oncogene-bearing ecDNAs and also found satellite-associated, hypomethylated DNAs to be enriched near somatic SV breakpoints as prominent features of the tumors studied. Given the extensive molecular and clinical parallels between canine and human OS, these findings from a naturally occurring large animal model provide insights into repeat-driven instability and ecDNA-mediated oncogene amplification in canine OS, with direct implications for understanding and ultimately targeting these processes in human OS.

## Results

### Sample Characteristics and Sequencing Overview

We generated PacBio HiFi long-read whole genome sequencing data from matched tumor and peripheral blood samples from four dogs with osteosarcoma (OS): an 18-year-old male Rottweiler (Ro1759), a 12-year-old female Great Pyrenees (GP1899), an 11-year-old male Golden Retriever (GR2023), and a 6.5-year-old female Great Dane (GD2044) (*SI Appendix,* **Table S1**). Primary tumors arose in the humerus or radius, consistent with typical appendicular OS sites. HiFi sequencing provided 51× - 96× tumor genome coverage (*SI Appendix,* **Table S1**), enabling high confidence SV and methylation analyses. Estimated tumor purity ranged from 0.53 to 0.81, and genome-wide ploidy from 1.93 to 4.69 (**Figs. S1-S4**). For three dogs (Ro1759, GP1899, and GR2023), additional normal stromal tissue was available and sequenced to provide independent non-neoplastic controls.

### Single Nucleotide Variants and Mutational Spectrum

Long-read whole genome sequencing revealed substantial intertumor variability in somatic SNV burden across canine OS, with total counts ranging from 2,506 to 6,855 variants and mutation rates of 1.06 to 2.91 SNVs/Mb (median 1.95 SNVs/Mb; **Fig 1A**; *SI Appendix,* **Table S2-S3**). These burdens are consistent with prior canine OS studies (27) and comparable to those reported for human pediatric OS (9), supporting a shared low-to-moderate mutational burden across species.

**Fig. 1.**
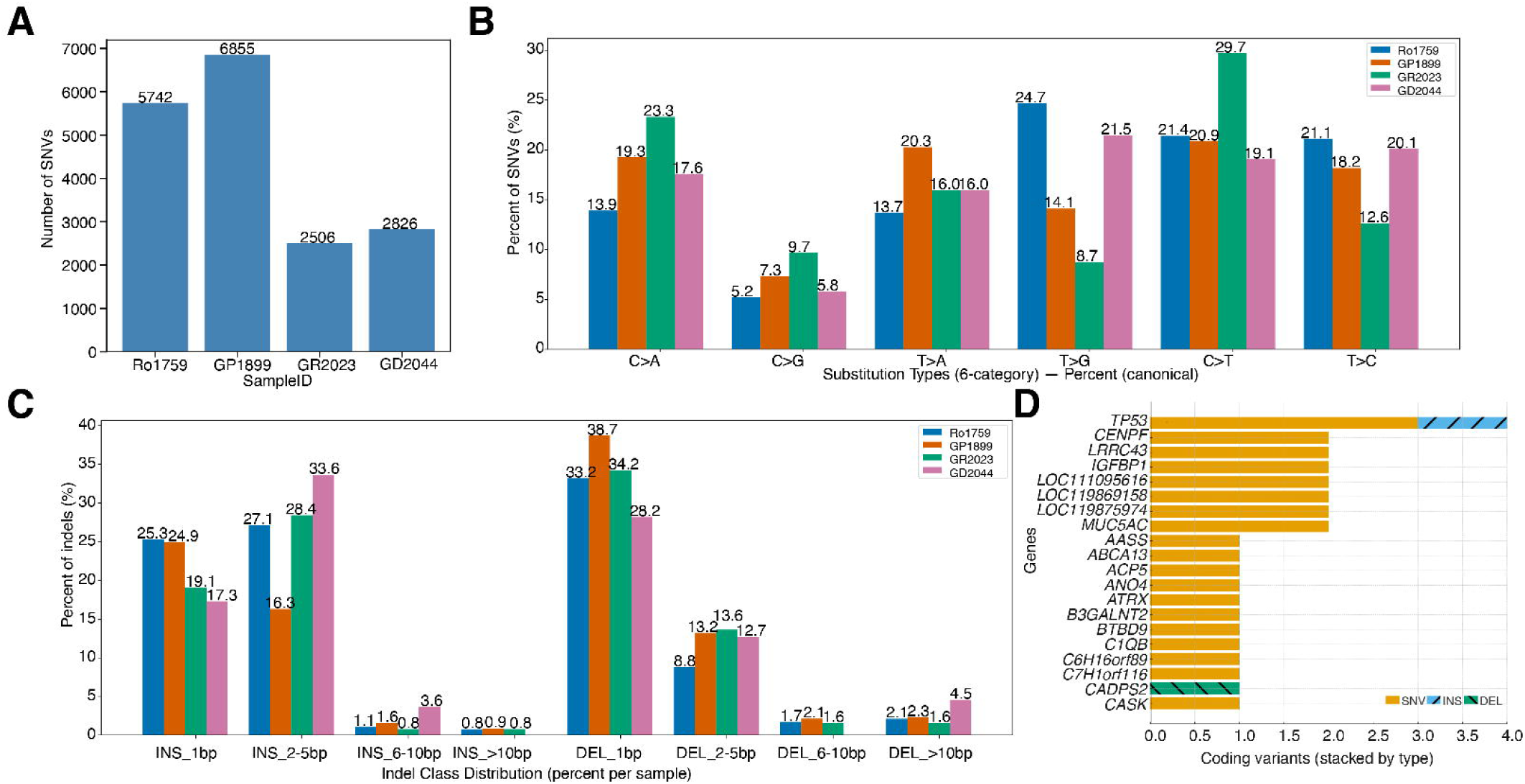
Somatic SNV and indel landscapes in four canine osteosarcomas. (A) Total number of somatic single nucleotide variants (SNVs) per tumor. (B) Counts of somatic SNVs stratified by base substitution class (C>A, C>G, C>T, T>A, T>C, T>G) for each tumor. Transversions (C>A, C>G, T>A, T>G) are more frequent than transitions (C>T, T>C) in all four tumors (binomial test, p < 10⁻⁷ for all samples). (C) Somatic insertion and deletion (indel) counts grouped by event type (insertion vs deletion) and length bin (1 bp, 2-5 bp, 6-10 bp, >10 bp). Single-base deletions (DEL_1bp) are the most common class overall (ranging from 31 to 399), followed by single-base insertions (INS_1bp). Larger indels (>10bp) are rare. Sample GP1899 exhibits substantially higher indel counts than the other tumors. (D) Recurrently mutated genes showing coding variants stacked by variant type. Bar segments are subdivided by variant type (SNV in orange, insertions in blue hatched, deletions in teal hatched).

SNVs were overwhelmingly noncoding, with only 0.89% mapping to coding sequences (CDS) and the remainder localized primarily to introns and untranslated regions (UTRs) (**Fig. S5**). Accordingly, CDS-restricted mutation rates were substantially higher (20.0-54.8 SNVs/Mb; *SI Appendix,* **Table S3**) reflecting strong selective constraint on protein-coding regions. The marked depletion of coding mutations despite their higher occurrence rate indicates that most protein-altering changes are incompatible with tumor cell fitness (28, 29), as cancer cells depend on preserving the function of essential genes required for basic cellular processes and tumor survival (30). Functional annotation identified only 0.42-0.77% of SNVs as having moderate or high predicted impact, including alterations in established cancer-associated genes such as *TP53*, *RB1* (*SI Appendix,* **Table S4**). Consistent with the genetic heterogeneity characteristic of OS, only three SNVs were shared across tumors (*SI Appendix,* **Table S5**).

The mutational spectrum was dominated by transversions, yielding a low transition/transversion ratio (Ti/Tv = 0.685; **Fig. 1B**) and distinguishing these tumors from transition-enriched, aging associated mutational patterns reported previously in canine OS (27, 31). Instead, G>T and C>A transversions were enriched, consistent with oxidative DNA damage-associated mutagenic processes (32, 33). This transversion-dominated mutational profile parallels patterns observed in human OS, where >80% of tumor exhibit *BRCA*-deficiency-associated signature characterized by specific single-base substitutions and oxidative damage markers (34, 35). Sample specific analyses further revealed additional heterogeneity with distinct enrichment of C>T, C>A, or T>G substitutions across individual tumors, indicating variability in dominant mutational processes within this cohort (*SI Appendix,* **Table S6**).

Chromosomal distribution of SNVs was broadly uniform with localized enrichments on specific chromosomes in individual tumors (e.g., CFAX and CFA22 in GP1899; **Fig S6**) suggesting regional mutation accumulation rather than genome-wide mutational bias. This pattern is consistent with the heterogeneous chromosomal instability patterns reported in human OS, where distinct chromosomes and genomic regions are variably affected across tumors (9, 36).

Intersecting somatic variants with canFam4 CDS annotations identified 142 coding variants across the cohort, with modest per-tumor counts (18–59 variants; *SI Appendix,* **Table S7**). These mutation burdens are comparable to human OS, which similarly displays low coding mutation frequencies (median 36 mutations, range 7-194 per tumor (37)) and generally low point-mutation burden relative to many adult cancers (9).

### Insertion and Deletion (indel) Characteristics

Across the four tumors, we identified 2,058 somatic indels (<50 bp), corresponding to indel burdens of 0.047–0.438 indels/Mb (*SI Appendix,* **Tables S8–S9**). Indels were dominated by short events. Nearly 60% were single–base pair insertions or deletions with frequency declining sharply as indel size increased and fewer than 5% exceeding 10 bp (**Fig. 1C**). This size distribution is consistent with replication-associated mutational processes rather than large-scale DNA repair defects.

Indels were overwhelmingly noncoding, with only 1–28 CDS-overlapping events per tumor and just three coding indels observed across the entire cohort (**Fig. S7**). This extreme depletion from coding regions (∼99% noncoding) is consistent with strong purifying selection against frameshift and protein-disruptive mutations. Accordingly, functional annotation showed that nearly all indels were predicted to have minimal impact, with rare, non-recurrent coding events and exceptionally few frameshift mutations (five genes total, each observed once; *SI Appendix,* **Table S10**). These patterns indicate that indels contribute little to the functional mutational burden of canine OS compared with SNVs.

Chromosome-level analyses revealed non-uniform indel distribution, with recurrent enrichment on CFA13, CFA22, CFA31, and CFAX (**Fig. S8**). These enrichments persisted after correction for chromosome size and coding content. Their overlap with regions previously implicated in copy-number alterations in canine OS, argues against technical bias.

Analysis of indel formation mechanisms supported replication slippage as the dominant source for the large number of short insertions and deletions observed. There appeared to be limited contribution from other mechanisms such as mismatch repair deficiency or repeat expansion processes (*SI Appendix,* **Table S11**). These findings indicate that while small indels are common in canine OS genomes, they are largely biologically neutral and unlikely to be major drivers of genomic instability in this disease.

### Genes Affected by Somatic SNVs and Indels

*TP53* was the only gene recurrently mutated in all four tumors (**Fig. 1D**), consistent with prior reports documenting near-universal *TP53* disruption in canine OS and human human OS where *TP53* is altered in approximately 90% of cases through both point mutations and SVs (11, 38), supporting its role as a core driver of disease pathogenesis across species. Additional recurrently mutated genes overlapped with those implicated in human OS and included *CENPF, LRRC43, IGFBP1, MUC5AC, AASS,* and *ABCA13* (**Fig. S9,** *SI Appendix,* **Tables S12, S13**), as well as tumor suppressors previously linked to OS biology (*LSAMP, DACH1*) (11, 34). Pathway-level analysis revealed convergence of affected genes on cancer-relevant networks involving WNT/β-catenin, PI3K/AKT/mTOR, TGF-β signaling, cell adhesion and epithelial–mesenchymal transition, metabolism, and cell-cycle regulation (*SI Appendix,* **Table S14**), indicating that despite limited coding mutation burden, recurrent disruption of core oncogenic pathways is preserved in canine OS and mirrors the pathway-level convergence observed in human OS. These pathways—including PI3K/AKT/mTOR, WNT/β-catenin, and TGF-β signaling—are consistently implicated in human OS pathogenesis (9, 39, 40), reinforcing the translational relevance of canine OS as a comparative model for human disease.

### Extensive Structural Variation and Complex Rearrangements

Genome-wide analysis of structural variants (SVs ≥50 bp) revealed pervasive and highly heterogeneous structural genome instability across canine OS. Total SV counts ranged from 184 to 738 events per tumor, corresponding to an approximately threefold difference in overall SV burden, while SV density within coding regions varied by more than 200-fold across samples (1.9–421.8 SVs/Mb CDS; **Fig. 2A**; **Tables S15–S16**). Breakend (BND) events predominated in all tumors, followed by deletions (DELs), duplications (DUPs), and insertions (INSs), consistent with extensive genome rearrangement rather than simple copy number alteration (*SI Appendix*, **Figs. S10–S13**). SV size distribution differed by class: DUPs were the largest events (median ≈41 kb), spanning 150 bp to 42 Mb; DELs had a median size of 9 kb and ranged from 50 bp to 18 Mb; INSs were substantially smaller, with a median of 240 bp and lengths ranging from 50 bp to 6.3 kb (**Fig. S14**).

**Fig. 2.**
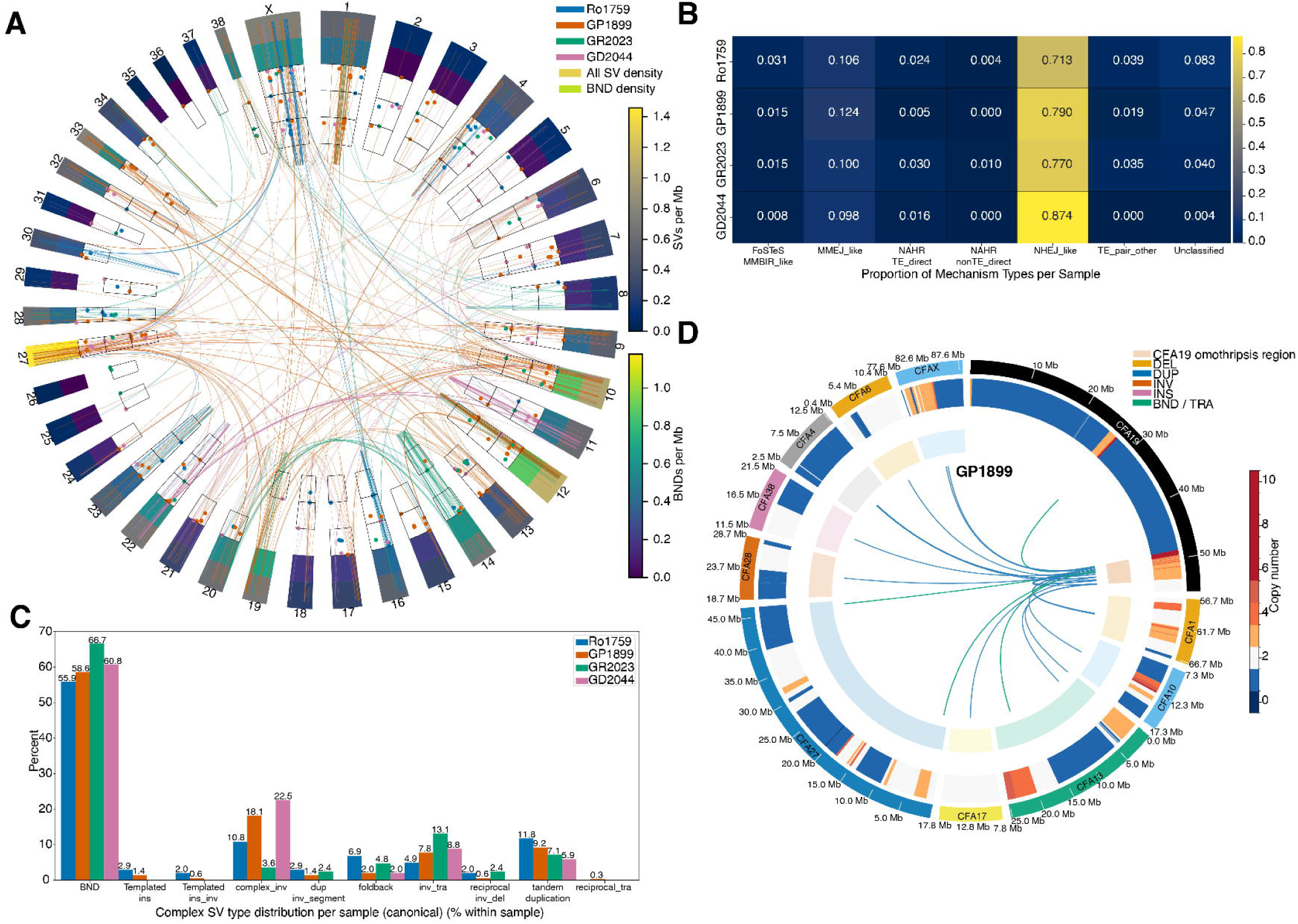
Structural variant architecture and inferred formation mechanisms in four canine osteosarcomas. (A) Circos plot summarizing somatic structural variants (SVs) across all autosomes and the X chromosome for tumors Ro1759, GP1899, GR2023, and GD2044. Concentric rings (outer to inner) show: chromosome ideograms, total SV density, breakend (BND) density, and per class densities for deletions (DEL), insertions (INS), duplications (DUP), inversions (INV), and copy number variant SVs (CNV). Colored arcs represent BND links connecting rearranged loci. Each tumor is plotted in a distinct color (legend, top right). (B) Distribution of complex SV classes in each tumor. Bars show counts of BND-derived architectures including templated insertions (Templated_ins), templated insertions with inversions (Templated_ins_inv), complex inversions (complex_inv), duplicated inverted segments (dup_inv_segment), foldback inversions, inversion-translocation events (inv_tra), reciprocal inversions with associated deletions (reciprocal_inv_del), tandem duplications, and reciprocal translocations (reciprocal_tra). (C) Proportional contribution of inferred SV formation mechanisms by tumor. Each cell shows the fraction of SVs assigned to a given mechanism class: Fork Stalling and Template Switching/Microhomology-Mediated Break-Induced Replication-like (FoSTeS_MMBIR_like), Microhomology-Mediated End Joining-like (MMEJ_like), Non-Allelic Homologous Recombination with transposable elements direct (NAHR_TE_direct), NAHR non-TE direct (NAHR_nonTE_direct), Non-Homologous End Joining-like (NHEJ_like), TE pair other, and Unclassified. (D) Focal view of a chromothripsis region on CFA19 in tumor GP1899. The Circos plot displays CFA19 (outer ring) and its rearrangement partners (CFA1, CFA4, CFA6, CFA10, CFA13, CFA17, CFA27, CFA28, CFA38, and CFAX). Genomic coordinates are marked every 5 Mb. From outer to inner, tracks depict copy number profile along CFA19 (heatmap; copy number 0-10), SV classes on CFA19 [deletions (DEL, orange), duplications (DUP, blue), inversions (INV, green), insertions (INS, red), and breakends/translocations (BND/TRA, purple)]. Arcs are connecting clustered breakpoints and are colored by cluster assignment where present. Partner chromosomes are color-coded with ideograms shown around the periphery. This pattern of dense, localized rearrangements and oscillating copy number is consistent with chromothripsis centered on CFA19.

SVs were distributed non-uniformly across the genome, with multiple chromosomes showing significant enrichment or depletion after normalization for chromosome length (**Fig. S15**).

Across the cohort, CFA12, CFA1, CFA10, CFA27, and CFAX, harbored the highest cumulative SV burdens. A pronounced hotspot on CFA12 in tumor GP1899 encompassed focal duplications and intrachromosomal rearrangements extending into the dog leukocyte antigen (DLA) region, suggesting localized structural instability involving immune-related loci.

Annotation of SV junctions based on microhomology length, inserted sequence, and repeat-mediated homology revealed a consistent predominance of error-prone repair mechanisms across tumors (**Fig. 2B**, *SI Appendix,* **Table S17**). Non-homologous end joining (NHEJ-like) events accounted for 70.1–81.7% of SVs, with microhomology-mediated end joining (MMEJ-like) contributing an additional 10.2–11.4%. Replication-associated mechanisms such as FoSTeS/MMBIR (0.8–2.8%) and homology-driven recombination (NAHR; 2.8–7.0%) were comparatively rare. Together, these patterns indicate that double-strand break repair with little or no homology is the dominant mode of SV formation in canine OS, with microhomology-mediated processes playing a consistent secondary role.

Long-read sequencing resolved extensive rearrangement complexity, revealing hundreds of multi-breakpoint SVs per tumor (161–438 BNDs), including templated insertions, foldback inversions, inverted translocations, and clustered reciprocal rearrangements (**Fig. 2C**). Many SVs were organized into discrete breakpoint clusters spanning one or more chromosomes, consistent with chromothripsis- and chromoplexy-like genome remodeling processes (**Table S18, S19**). Notably, tumor GP1899 exhibited a dense cluster of interconnected rearrangements on CFA19 displaying alternating segment orientation and size, a pattern characteristic of chromothripsis (**Fig. 2D**). Optical genome mapping independently validated the majority of HiFi-detected breakpoints (**Fig. S16**), enabling reconstruction of this localized catastrophic event from base-pair to megabase scales and confirming its substantial impact on tumor genome architecture. Despite preferential localization of SV breakpoints to intronic and other noncoding regions (**Fig. S17**), gene-level annotation identified recurrent, high-impact SVs affecting cancer-relevant genes (*SI Appendix,* **Table S20**). In tumor Ro1759, interchromosomal rearrangements resulted in transcript ablation of *VCL*, consistent with complete functional knockout. Tumor GR2023 harbored multiple independent BND events generating recurrent *GRIP1–LRCH2* gene fusions with predicted truncation of both partners, implicating disruption of cytoskeletal organization and signaling pathways. In tumor GP1899, an inversion–translocation event produced high-impact fusions involving *CALHM4* and truncation of the DNA repair polymerase *POLB*. In contrast, tumor GD2044 contained a single large tandem duplication spanning 8.3 Mb on CFA1, affecting hundreds of genes including *BCL2* and *TNFRSF11A*. This is consistent with a dosage-driven copy-number gain rather than focal transcript disruption.

Across tumors, while most SV-associated annotations were predicted to have modifier or moderate effects, a recurrent subset of high-impact rearrangements directly disrupted genes involved in cytoskeletal regulation, DNA repair, apoptosis, and signaling. Collectively, these findings demonstrate that canine OS genomes are shaped by extreme, error-prone structural remodeling, in which clustered SVs arising from non-homologous repair processes contribute both to large-scale genome reorganization and to focal disruption of cancer relevant genes. The extent and heterogeneity of SVs observed here are highly concordant with prior canine and human OS studies describing a structurally dominated, “chaotic” genomic landscape, in which chromosomal lesions represent the predominant mutation class. Previous analyses of canine OS tumors and cell lines have reported recurrent disruption of genes such as *DMD, DLG2,* and *SETD2*, as well as extensive rearrangement across both primary and metastatic disease (27, 31, 41). Our data reinforce these observations and extend them by resolving complex, multi-breakpoint SV architectures in primary tumors using long-read sequencing. Notably, the pronounced enrichment of SVs on specific chromosomes (*e.g.*, CFA12) and the presence of distinct SV size classes suggest contributions from both localized DNA damage repair errors and larger-scale catastrophic events. The predominance of intronic and other noncoding SVs, consistent with reports in human OS, further underscores the importance of large-scale regulatory and architectural genome disruption (rather than recurrent point mutations) as a central feature of OS pathogenesis.

### Differential enrichment of repetitive elements at somatic variant breakpoints

To determine whether specific repetitive genomic elements mark regions of heightened mutational susceptibility in canine OS, we performed permutation-based enrichment analyses of repetitive sequences at SNV, indel, and SV breakpoints across the four tumors. This variant-stratified approach revealed highly reproducible, repeat-class associations that differed markedly by mutation type, indicating distinct underlying mutational mechanisms.

At SNV breakpoints, pericentromeric satellite repeats (most notably SAT1_CF) showed the strongest and most consistent enrichment, observed in three of four tumors (19.3–46.3-fold; **Fig. 3A, B**; *SI Appendix,* **Table S21**). Simple tandem repeats, particularly tetranucleotide motifs, were also recurrently enriched at SNV breakpoints in most tumors, whereas SINE elements and ancient MIR families were consistently depleted (**Table S21**). In contrast, the canine-specific retrotransposon subfamily, L1_Canis1, exhibited universal SNV breakpoint enrichment across all four samples (**Fig. 3C**), with enrichment strongest in evolutionarily young LINE-1 subfamilies, consistent with preferential mutability at recently active elements.

**Fig. 3.**
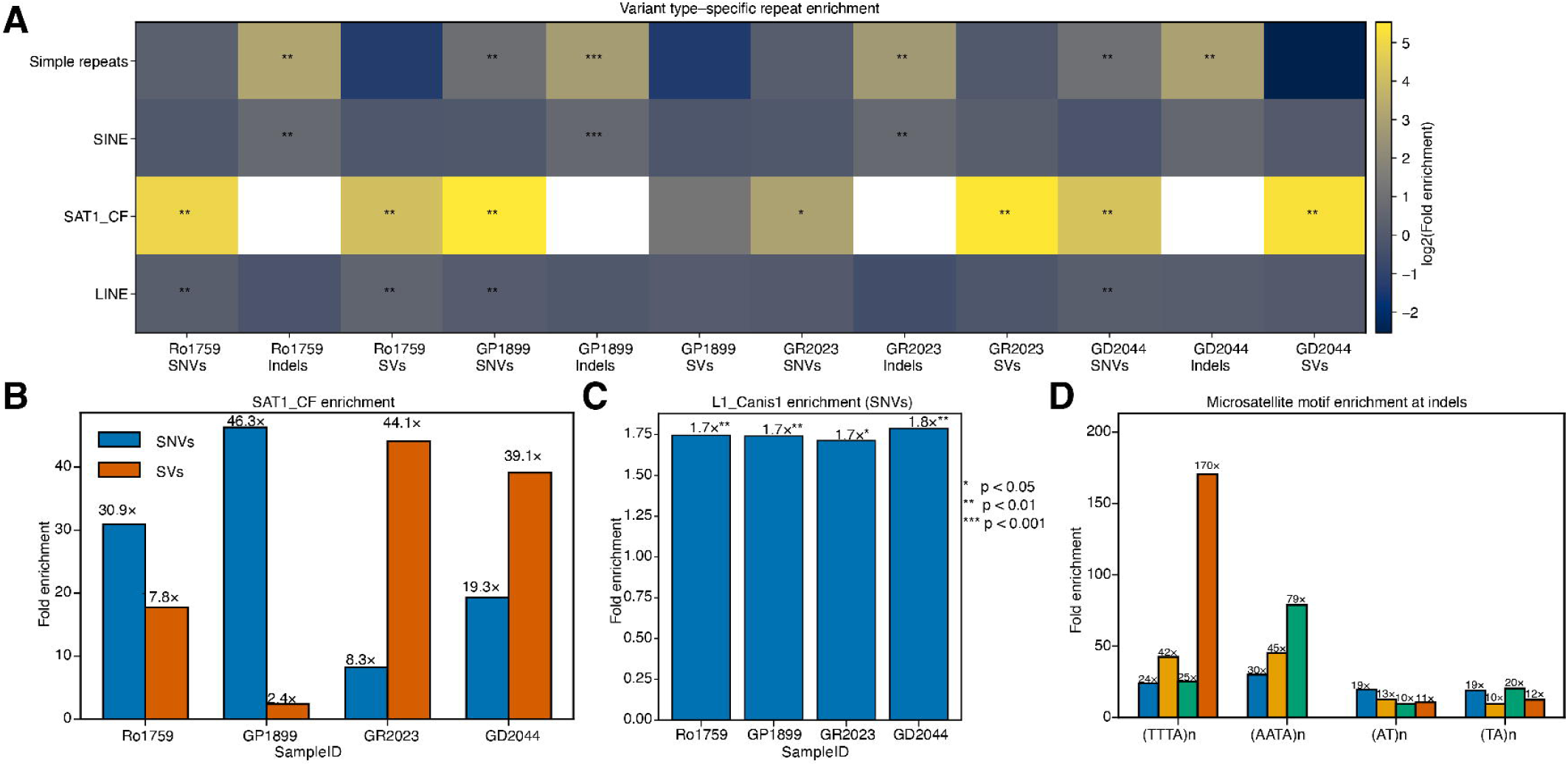
Enrichment of different variant types in repetitive genomic elements across dog samples. (A) Heatmap showing log2 fold enrichment of SNVs (single nucleotide variants), indels (insertions/deletions), and SVs (structural variants) across different repeat classes (simple repeats, SINE, SAT1_CF, and LINE) in four samples (Ro1759, GP1899, GR2023, GD2044). Asterisks indicate statistical significance (*p < 0.05, **p < 0.01, ***p < 0.001). (B) Fold enrichment of SNVs (blue) and SVs (orange) in SAT1_CF satellite repeats across the four samples. SNVs show consistently higher enrichment (19.3-46.3×) compared to SVs (2.4-44.1×), with GP1899 showing the highest SNV enrichment. (C) Fold enrichment of SNVs in L1_Canis1 (LINE-1) elements across samples, showing modest but consistent enrichment (1.7-1.8×) with statistical significance (**p < 0.01, *p < 0.05) in all samples. (D) Enrichment of different microsatellite motifs at indel sites. (TTTA)n and (AATA)n repeats show the highest enrichment (up to 170× and 79×, respectively), while (AT)n and (TA)n motifs show lower enrichment levels (10-20×) across samples.

Indel breakpoints displayed a distinct and highly convergent pattern. Simple repeats were the dominant sequence context for indel formation, showing strong and uniform enrichment across all tumors (6.5–9.1-fold; **Fig. 3D**; *SI Appendix,* **Table S22**), particularly at thymine-rich microsatellite motifs (adjusted p-value = 0.00112). This pattern supports replication slippage as the primary mechanism underlying indel formation. SINE elements, including tRNA-derived families and carnivore-specific SINEC subfamilies, showed moderate but consistent enrichment, whereas LTR retrotransposons and evolutionarily ancient LINE families (e.g., L2, MIR) were recurrently depleted, establishing a hierarchy of indel susceptibility governed by repeat type, sequence composition, and evolutionary age.

SV breakpoints exhibited a contrasting repeat landscape: SAT1_CF centromeric satellites showed the most consistent enrichment, detected in three of four tumors (17.8–44.1-fold; **Fig. 3B**; *SI Appendix,* **Table S23**), with broader satellite classes also enriched in multiple samples. In striking contrast to indels, simple repeats were universally depleted at SV breakpoints across all tumors (0.17–0.99-fold), indicating that repeat contexts prone to polymerase slippage are not major substrates for large-scale rearrangement. Retrotransposon-associated SV enrichment was more heterogeneous. Some tumors showed enrichment of LINE-1 or LTR subfamilies, including dog-specific L1 elements, whereas others did not, suggesting tumor-specific contributions from euchromatic retrotransposon-associated processes.

Across variant classes, these analyses reveal a reproducible and mechanistically coherent pattern: (1) Simple repeats act as primary hotspots for indels but are coldspots for SVs. (2) Centromeric SAT1_CF satellites are recurrently enriched at both SNV and SV breakpoints. (3) SINE and LINE elements exhibit variant-type–specific and age-dependent associations. Together, these findings demonstrate that somatic mutagenesis in canine OS is strongly shaped by local repetitive sequence context, with distinct mutational processes operating preferentially within specific genomic environments. The convergence of SNV and SV breakpoints at centromeric satellite DNA, in particular, nominates these regions as recurrent sites of structural fragility and aberrant repair, motivating further analysis of their epigenetic state and contribution to genome instability.

Given the recurrent enrichment of SV breakpoints at centromeric satellite repeats, we next asked whether these rearrangements are associated with localized epigenetic alterations, such as DNA hypomethylation, that are characteristic of structurally unstable genomic regions.

### Local hypomethylation at structural variant breakpoints

Analysis of CpG methylation inferred from HiFi sequencing kinetics revealed focal hypomethylation at somatic SV breakpoints in tumors relative to matched normal samples (*SI Appendix,* **Table S24**). SV breakpoints were frequently located within, or immediately adjacent to, hypomethylated differentially methylated regions (DMRs), indicating a strong spatial association between SV and localized methylation loss.

Overlaying SV breakpoints and SV-spanning intervals with tumor-intrinsic DMRs and gene annotations identified 107–506 genes intersected by SVs per tumor (*SI Appendix,* **Table S25**). Among SVs overlapping DMRs (SV-DMRs), 69 SVs intersected hypomethylated DMRs, whereas only 5 SVs overlapped hypermethylated regions (*SI Appendix,* **Table S25**), revealing a pronounced bias toward hypomethylation at sites of SV. Of these SV–DMR overlaps, 30 occurred within gene bodies or promoter regions, highlighting candidate loci at which structural rearrangement and epigenetic alteration co-occur.

Within individual tumors, several genes were disrupted by multiple independent SVs, consistent with locus-level clustering of structural instability. These included *PCSK5, SIPA1L2,* and *RUNX2* in tumor GP1899, as well as *GRID1, GRIP1,* and *CCDC88C* in other tumors (*SI Appendix,* **Table S25**). Genes affected by hypomethylated SVs included examples with functions relevant to OS biology, including osteogenic differentiation (*RUNX2*), cytoskeletal organization (*GRIP1, GRID1, SIPA1L2*), signaling and transcriptional regulation (*AUTS2, CCDC88C*), and chromatin-associated processes (*KANSL3*) (*SI Appendix,* **Figures S18-S23**). Collectively, these findings demonstrate that SVs in canine OS are consistently associated with focal hypomethylation and tend to cluster at specific genomic loci within tumors.

### Extrachromosomal DNAs concentrate cancer-associated genes and canonical OS pathways

Extrachromosomal DNA (ecDNA) was detected in 3 of 4 tumors (75%) and was absent from matched normal samples, indicating tumor-specific formation (**Fig. 4A, B**). Across these tumors, we identified 16 distinct circular amplicons (Ro1759: n=4; GP1899: n=11; GD2044: n=1) ranging from 103 kb to 3.07 Mb in size (median 274 kb) and amplified to copy numbers of 3.3–8.4 (median 4.5), consistent with focal oncogenic amplification. Six ecDNAs met stringent Tier 1 criteria, while ten were classified as Tier 2 (probable ecDNA) (*SI Appendix*, **Figures S23-S31**). Notably, nearly one-third (5/16) contained interchromosomal junctions involving segments from multiple chromosomes, consistent with complex, chromothripsis-like ecDNA formation mechanisms (*SI Appendix*, **Figures S23-S31**).

**Fig. 4.**
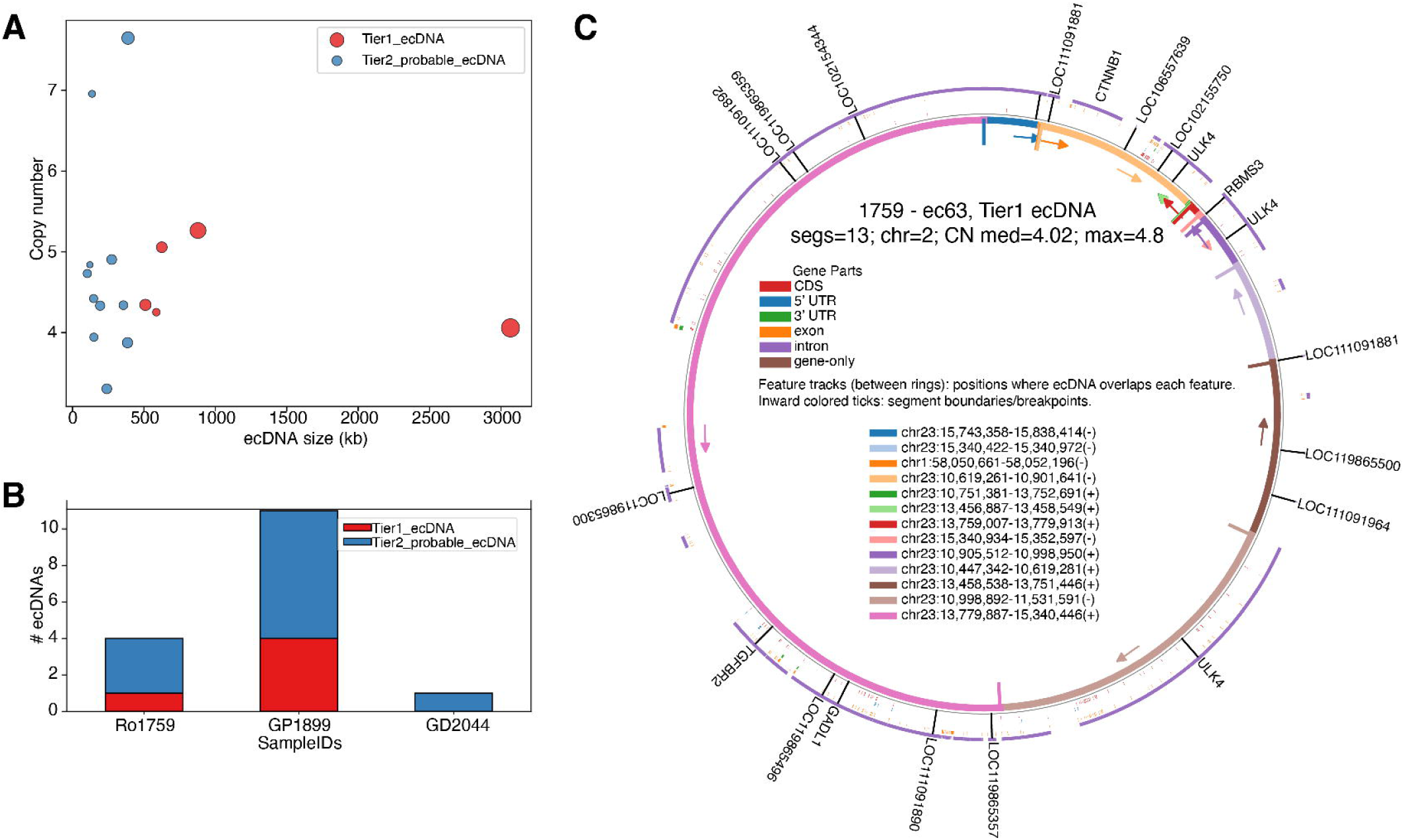
Extrachromosomal DNA (ecDNA) burden, oncogene content, and architecture in osteosarcoma. (A) Relationship between ecDNA size (kb) and copy number. Each point represents a circular amplicon detected across the three ecDNA positive tumors (Ro1759, GP1899, GD2044). The x-axis shows ecDNA size and the y-axis shows estimated copy number. Tier 1 ecDNA (high-confidence circular amplicons) are shown in red and Tier 2 probable ecDNA are shown in blue. The dot size is proportional to the size of ecDNAs. (B) Per-tumor ecDNA burden. Stacked bars indicate the number of ecDNAs detected in each tumor (Ro1759, GP1899, and GD2044), stratified by Tier 1 (red) and Tier 2 probable (blue) elements. (C) Architecture of a representative Tier 1 ecDNA (ec63) from tumor Ro1759. Circular plots depict a 3.07 Mb ecDNA comprising 13 genomic segments (legend, bottom) with a median copy number of ∼4 (maximum 4.8). The outer track shows the reference chromosomal coordinates for the contributing segments. Inner annotation tracks mark gene features overlapped by the ecDNA (CDS, 5’ UTR, 3’ UTR, exons, introns, gene-only regions). Gene labels highlight notable loci carried on this ecDNA, including CTNNB1, TGFBR2, and neighboring genes. Inward colored vertical lines denote segment boundaries and junctions, illustrating the rearranged structure of the circular amplicon.

ecDNAs recurrently harbored genes implicated in OS-relevant signaling pathways. In tumor Ro1759, the largest ecDNA (ec63; 3.07 Mb; Tier 1) comprised 13 rearranged segments and amplified *CTNNB1* (β-catenin; copy number validated by ddPCR; **Fig. S32**) and *TGFBR2*, linking ecDNA formation to Wnt/β-catenin and TGF-β signaling (**Fig. 4C, Fig. S33**). A second ecDNA, ec4, in this tumor amplified *FGF7*, consistent with involvement of the FGF/FGFR axis in OS proliferation (**Fig. S33**, *SI Appendix,* **Table S26**). In tumor GP1899, a high-copy ecDNA encompassed *MDM2* (copy number 6.55) together with *FRS2* and *YEATS4*, recapitulating the canonical 12q13–15 amplicon architecture characteristic of human OS (**Fig. S33**, *SI Appendix,* **Table S26**). Additional ecDNAs in this tumor included an amplicon containing *LSAMP*, with circular junctions independently validated by PCR (**Fig. S34**).

Across tumors, ecDNA-associated genes converged on a limited set of oncogenic pathways (including p53/*MDM2*, Wnt/β-catenin, TGF-β, and FGFR signaling) despite heterogeneity in ecDNA number, structure, and chromosomal origin. These findings indicate that ecDNA-mediated amplification is a recurrent feature of canine OS and provides a flexible, high-copy substrate for activation of core oncogenic pathways. These results reinforce ecDNA as a central mechanism of genome remodeling and pathway dysregulation in this disease.

## Discussion

Using long-read whole-genome sequencing, we show that canine osteosarcoma (OS) is characterized by extreme structural genome remodeling coupled with focal epigenetic disruption, closely mirroring key features of human OS. Although based on a modest cohort, the depth and resolution of these data refine existing models of OS pathogenesis and reinforce the value of spontaneous canine OS as a translational model for studying structural and epigenetic drivers of cancer evolution.

### Structural genome remodeling as a defining feature of OS

Across tumors, structural variation dominated the mutational landscape, with extensive multi-break rearrangements, clustered breakpoint architectures, and complex configurations that are difficult to resolve using short-read approaches. These findings reinforce prior work in human and canine OS identifying SVs (not point mutations) as the principal mode of genomic alteration. The chromothripsis-like event resolved on CFA19 provides direct evidence that chromosome-scale catastrophic rearrangements occur in spontaneous canine OS, consistent with observations in human disease. Whether such events arise early during tumor initiation or continue to accumulate during progression likely varies by tumor; our observation of substantial intertumor heterogeneity in SV burden and architecture suggests that OS evolution follows multiple trajectories rather than a single canonical path.

### Epigenetic context and structural fragility

A key insight from this study is the consistent association between SV breakpoints and focal hypomethylation. Long-read derived methylation profiles revealed sharp, localized loss of CpG methylation surrounding rearrangement junctions, with recovery toward baseline at increasing distance. This spatial pattern supports a close coupling between epigenetic relaxation and structural instability, potentially reflecting increased susceptibility of open chromatin to DNA breakage, replication stress, or error-prone repair. Importantly, all SVs overlapping differentially methylated regions intersected hypomethylated (but not hypermethylated) domains, indicating a strong directional relationship between methylation loss and rearrangement.

Repeat-enrichment analyses further implicate repetitive DNA as a substrate for this instability. While retrotransposons account for many breakpoint overlaps, centromeric satellite DNA (particularly the SAT1_CF family) showed the strongest and most consistent enrichment at both SNV and SV breakpoints. These sequences localize to heterochromatic, late-replicating regions previously linked to replication stress and double-strand break formation. Interpretation of this enrichment must account for canine chromosome architecture, as most canine chromosomes are acrocentric, placing pericentromeric satellites near chromosome ends. Without complete centromere annotations in the current reference genome, it remains unclear whether satellite enrichment reflects intrinsic sequence fragility, positional effects, or both. Nevertheless, these findings demonstrate that satellite-rich domains disproportionately participate in OS genome remodeling and highlight the power of long-read sequencing to interrogate highly repetitive regions that were previously inaccessible.

### Extrachromosomal DNA as a route to oncogenic amplification

We detect extrachromosomal DNA (ecDNA) in most tumors, with circular amplicons recurrently carrying genes central to OS biology, including *MDM2, CTNNB1, TGFBR2,* and *FGF7*. These ecDNAs recapitulate canonical oncogenic architectures described in human OS, such as the 12q13–15 amplicon, and frequently involve complex, multichromosomal rearrangements. Given their high copy number and non-Mendelian inheritance, ecDNAs provide a flexible mechanism for rapid oncogene dosage modulation and intratumoral heterogeneity. The convergence of ecDNA content on a limited set of pathways (p53/MDM2, Wnt/β-catenin, TGF-β, and FGFR signaling) suggests that ecDNA-mediated amplification represents a recurrent route to pathway activation in OS. Determining how these structures influence transcriptional programs, therapeutic sensitivity, and evolutionary dynamics will require integrated functional studies.

#### Conclusion and future directions

This study establishes that canine OS is shaped primarily by extensive structural genome remodeling rather than by accumulation of small coding mutations. Using long-read whole-genome sequencing, we provide the first integrated view of structural variation, repetitive sequence context, native DNA methylation, and extrachromosomal DNA in naturally occurring canine OS, revealing a cancer genome dominated by clustered rearrangements, focal hypomethylation, and ecDNA-mediated oncogene amplification.

Our findings identify satellite-rich genomic regions as recurrent sites of structural fragility, demonstrate that structural variants are consistently associated with localized methylation loss, and show that ecDNA frequently concentrates genes from canonical OS pathways. Together, these features point to replication stress, error-prone repair, and epigenetic destabilization as central forces shaping OS genome evolution. The convergence of ecDNA content on a limited set of signaling pathways further highlights ecDNA as a flexible substrate for oncogenic amplification and tumor adaptation.

Despite a limited cohort size, this work provides a foundational framework for understanding how structural and epigenetic instability interact to drive OS biology. Dogs with spontaneous OS offer a uniquely powerful translational system (combining biological relevance, clinical accessibility, and accelerated disease timelines) for testing hypotheses generated here. Future studies leveraging larger comparative cohorts, multiomic integration, and functional validation will be essential to define the causal relationships among satellite fragility, epigenetic disruption, and ecDNA formation, and to translate these insights into genome-informed therapeutic strategies for OS in both dogs and humans.

## Materials and Methods

### Sample collection

Blood, tumor, and stromal tissues were obtained from four client-owned dogs with OS, an 18 year old male castrated Rottweiler (sample Ro1759), 12 year old female Great Pyrenees (sample GP1899), 11 year old male castrated Golden Retriever (sample GR2023), and 6.5 year old female Great Dane (sample GD2044), that were presented to the Flint Animal Cancer Center at Colorado State University (CSU), Fort Collins, Colorado (*SI Appendix,* **Table S1**). All procedures were approved by the CSU Institutional Animal Care and Use Committee, and written informed consent was obtained from each owner. Dogs were selected for inclusion if they had treatment-naïve, primary appendicular OS documented by radiographic findings and confirmed by subsequent histopathology. Blood samples were collected via venipuncture prior to any surgical or medical intervention. Limb amputation was then performed, and tumor and stromal samples were harvested within 30 minutes of surgery. Tumor tissue was obtained using a stainless-steel Michele trephine biopsy needle, obtaining 3-5 cores from regions with clear mass effect and/or cortical bone lysis. Normal stromal tissue (bone and associated soft tissue) was sampled from anywhere around the tumor where there is grossly normal appearing tissue, to serve as a matched control. All tissues were either immediately flash-frozen in liquid nitrogen or processed fresh for downstream molecular analyses. Amputated limbs were subsequently submitted to the CSU Veterinary Diagnostic Laboratory for comprehensive histopathological confirmation and final tumor classification.

### PacBio HiFi long-read whole genome sequencing

Genomic DNA (gDNA) was extracted from blood and tissue samples using NEB Monarch cell/blood (NEB #T3050) and NEB Monarch tissue (NEB #T3060) kits, respectively. Briefly, 0.5-1ml of blood was mixed with 3 volumes of RBC Lysis buffer. Leukocytes were pelleted by centrifuge at 1000g at 4C for 10 min. For tissue samples, about 20-80mg of tissue was ground to a fine powder in liquid nitrogen. The pelleted cells or tissue powder were then lysed in the presence of proteinase K for 45 min. RNA was digested with RNase A for 10 min. Protein Separation Solution was added to remove proteins, and sample was centrifuged at 16000g for 10 min to pellet cell debris and proteins. The supernatant was treated with isopropanol to precipitate gDNA in the presence of glass beads. The sample was incubated at room temperature (RT) on a rotating mixer to allow gDNA binding to the glass beads. The supernatant was then removed, and the gDNA-bound glass beads were washed twice with Wash Buffer. The gDNA was eluted from the glass beads using Elution Buffer by incubation at 56C.

The library was prepared using SMRTbell prep kit 3.0 (PacBio). About 5ug of HMW DNA was first sheared using Megaruptor 3.0 (Diagenode). The sheared DNA was then cleaned up using 1X SMRTbell cleanup beads. The DNA was end-repaired, A-tailed and ligated with the barcoded SMRTbell adapters, followed by a cleanup to remove ligation enzyme and free adapters. This is followed by a nuclease treatment to remove DNA fragments without adaptors. Purification was performed using SMRTbell cleanup beads to remove excessive nuclease. Libraries were further subjected to gel size selection on Pippin HT (Sage Science) to remove small fragments <10kb. The size-selected and purified libraries were loaded on the Revio at 300-350pM. Sequencing was performed for 30 hours.

### Bionano Genomics optical mapping

Ultra-high-molecular-weight DNA was extracted from blood-tumor matched samples from one dogs (GP1899) using the Bionano Prep SP Fresh Cells DNA Isolation protocol, revision C (Document #30257), and the Bionano SP Blood & Cell DNA Isolation Kit (catalog #80030). In brief, 1.5 million cells were pelleted, resuspended in a lysis buffer containing detergents, proteinase K, and RNase A. DNA was then bound to a silica disk, washed, eluted, and homogenized by 1 hour of end-over-end rotation at 15 rpm followed by overnight equilibrium at room temperature. Isolated DNA was fluorescently labeled at CTTAAG motifs using the DLE-1 enzyme and counter-stained with the Bionano PrepTM DNA Labeling Kit – Direct Label and Stain (catalog #8005; Protocol revision F; Document #30206). Briefly, 750 ng of gDNA were incubated with DL-Green dye and DLE-1 enzyme in DLE-1 buffer for 2 hours at 37°C, followed by enzyme heat inactivation for 20 minutes at 70°C. The labeled DNA was treated with proteinase K at 50°C for 1 hour, and excess DL-Green dye was removed by membrane adsorption. The DNA was stored at 4°C overnight to facilitate DNA homogenization and then quantified using a Qubit dsDNA HS Assay Kit (Molecular Probes/Life Technologies). Prior to imaging, labeled DNA was stained with an intercalating dye and incubated at room temperature for at least 2 hours before loading onto a Bionano Saphyr chip. Molecules were linearized and imaged on Saphyr 2nd generation instruments (Part #60325) using Instrument Control Software (ICS) version 4.9.19316.1. The DNA Saphyr system software was used to measure backbone length and to detect the positions of fluorescent labels along each molecule. Resulting molecules were de novo assembled using Bionano Solve (v3.1) and aligned to canFam4 (42) dog genome reference assembly.

### Alignment of HiFi reads to reference genome

Raw PacBio HiFi reads were aligned to the CanFam4 (42) reference genome using pbmm2 (v1.17.0), a wrapper for minimap2 optimized for PacBio long reads. Alignments were executed via SLURM batch scripts, producing sorted and indexed BAM outputs for subsequent variant calling.

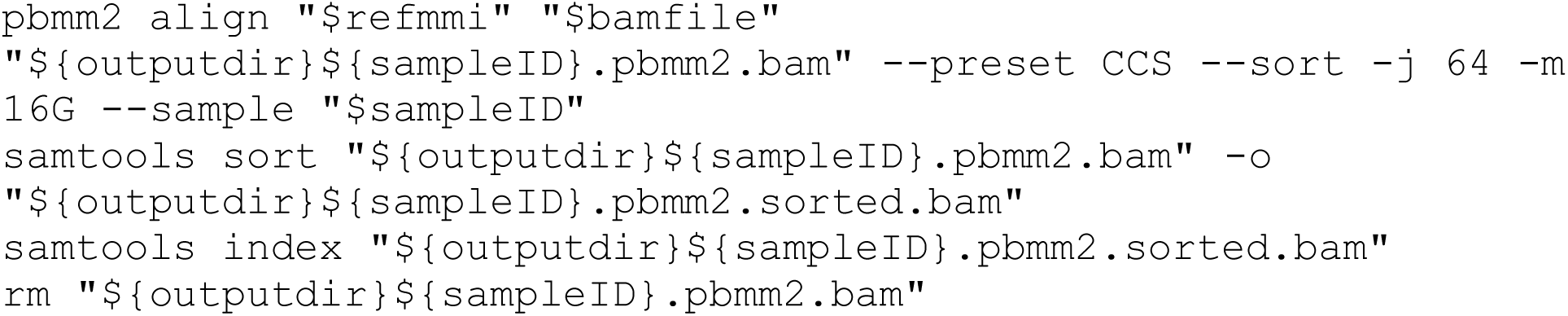

### Haplotagging

#### Small variant calling

Variant detection was performed using DeepVariant (v1.6.1) (43), run in CPU mode. For each sample, sorted BAM files were processed to produce GVCF (.vcf.gz) outputs containing high-confidence SNVs and small indels.

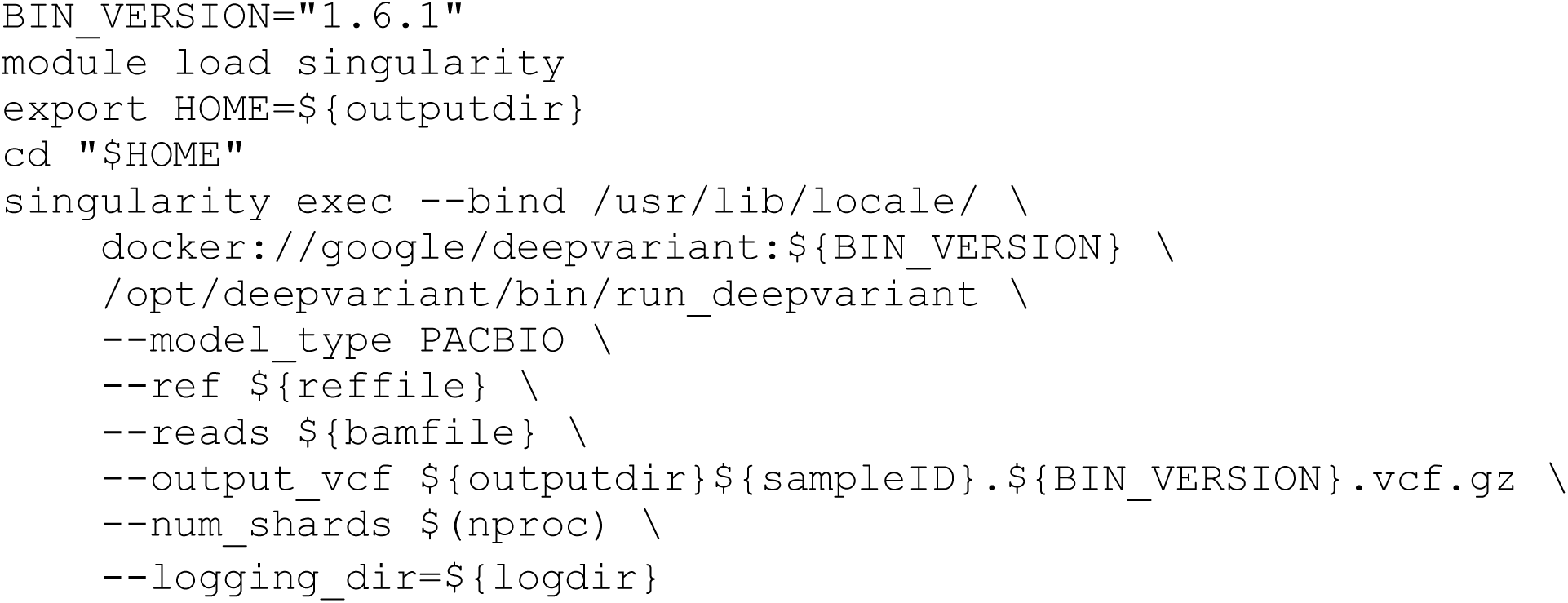

#### Variant phasing

Phasing of heterozygous variants was conducted using HiPhase (v1.5.0-8af3f1e) (44), which employs long-range read information to phase variants across haplotypes. Inputs included the aligned BAM file, the DeepVariant VCF (.vcf.gz), and the reference FASTA.

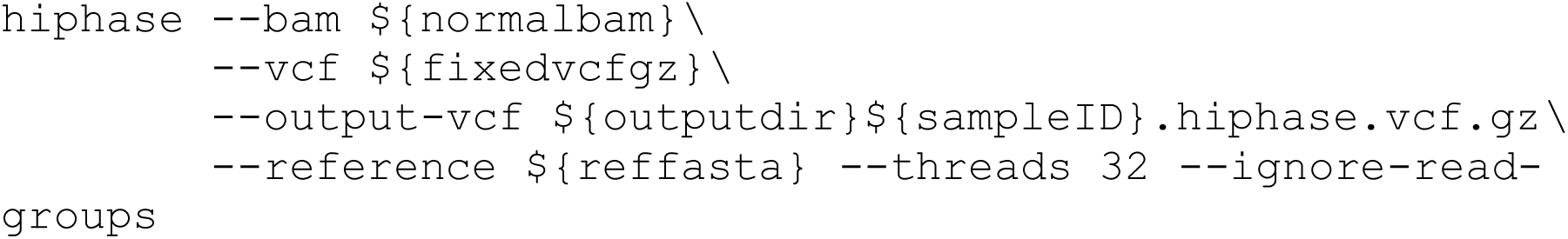

#### Haplotagging of reads

Reads were assigned to phased haplotypes using Whatshap (v2.5) (45), producing haplotagged BAM files (*.haplotagged.sorted.bam) for each sample. These files served as the foundation for downstream somatic variant detection and methylation analyses.

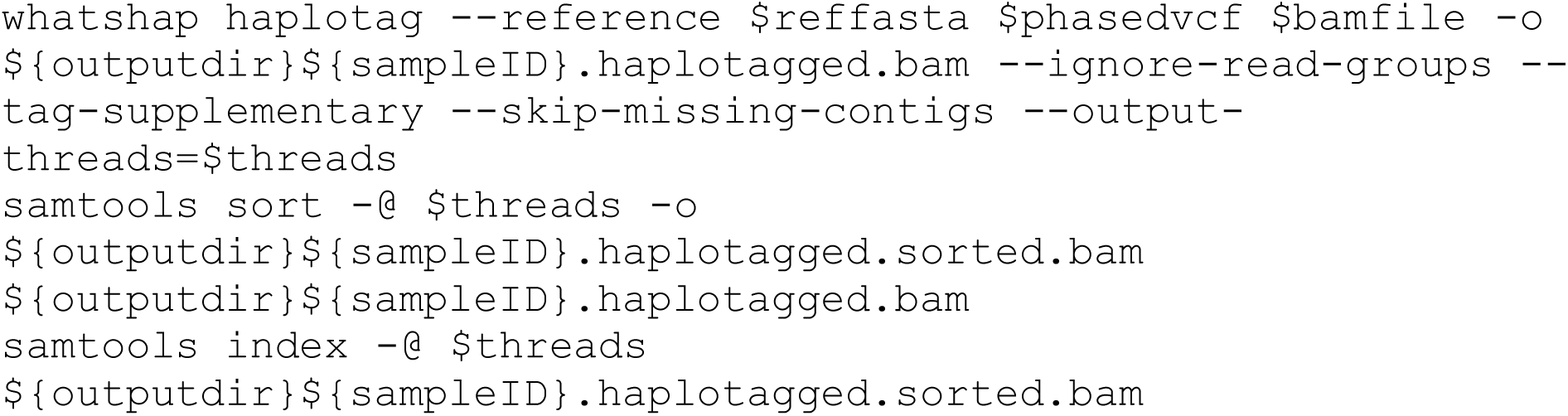

### Somatic variant calling

Somatic variant analysis was performed using multiple complementary pipelines to capture SNVs, indels, and structural variants (SVs). Tumor (*_T) and matched normal (*_B) samples were jointly analyzed.

#### Somatic SNVs and Indels

Two independent variant callers were implemented: ClairS (v0.4.1) (46), and DeepSomatic (v1.9.0) (47). Each tool was executed via SLURM.

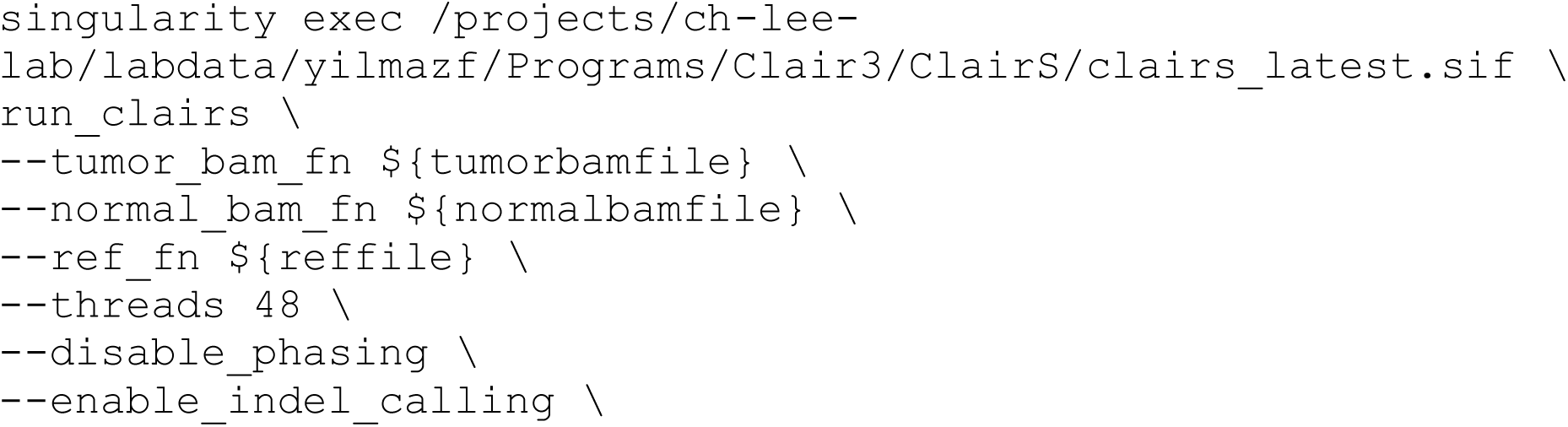
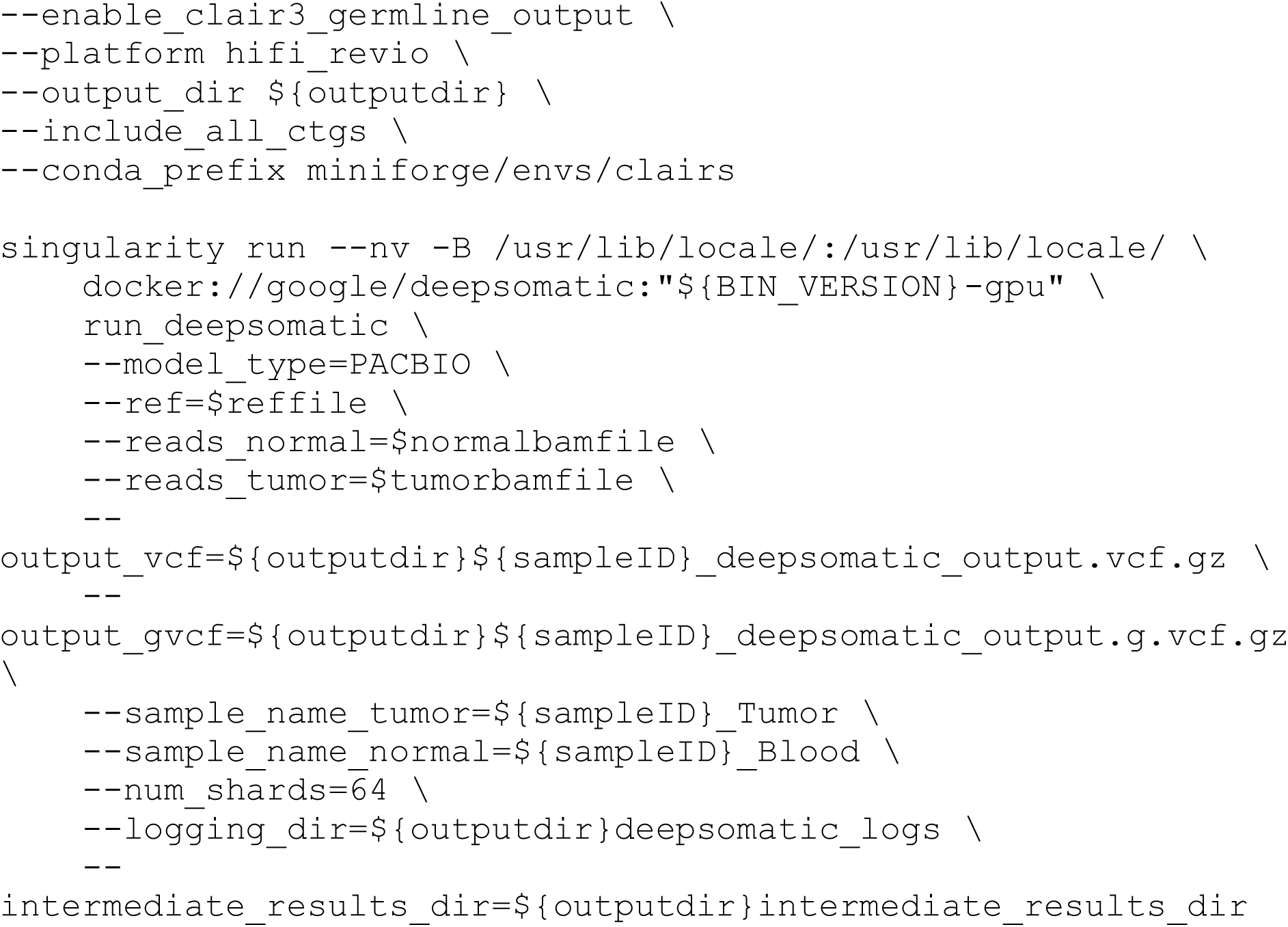

#### Somatic SVs

Two independent variant callers were implemented: savana (v1.3.5) (48), and severus (v1.6) (49). Each tool was executed via SLURM.

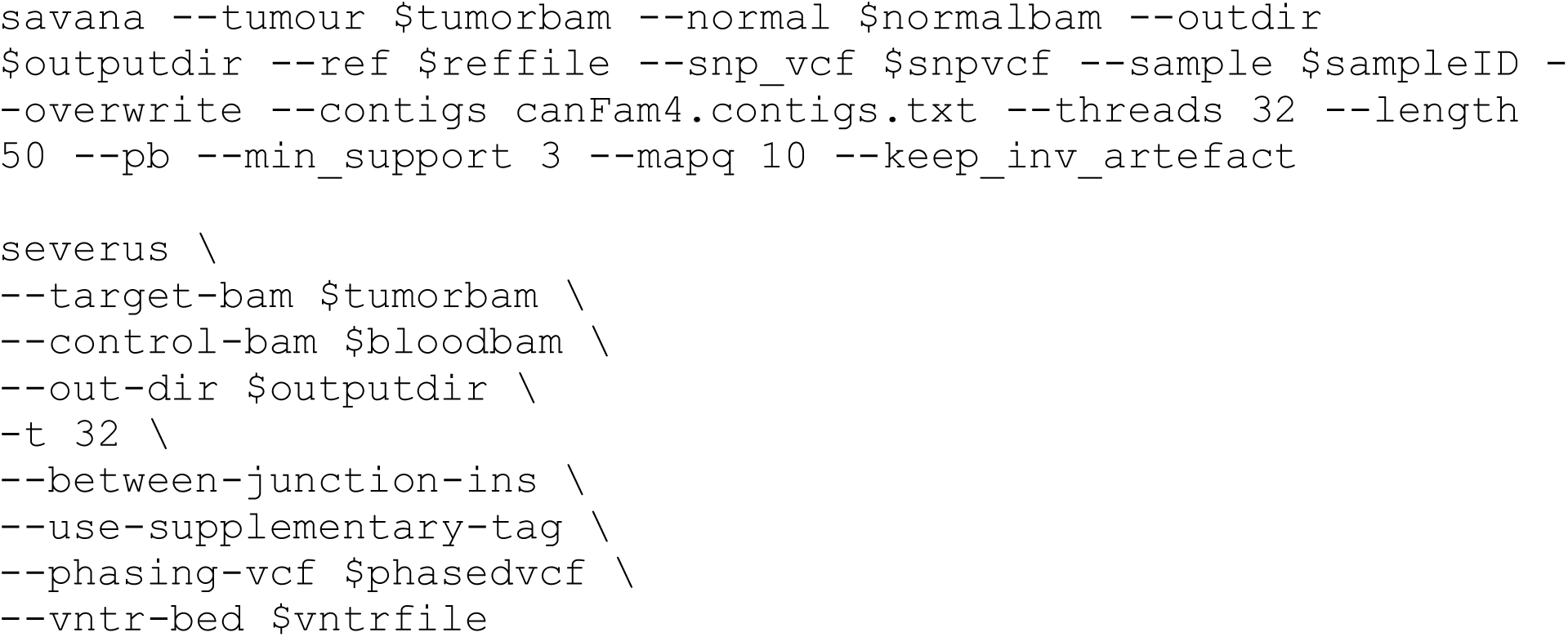

### Variant merging and consensus ensemble variant file generation

#### Ensemble integration

Final ensemble SNV and indel calls were constructed using a multistep approach (commands available under https://github.com/yilmazf/DogGenomesAnalyses). This ensemble served as the definitive variant set for downstream analyses.

Final ensemble SV calls were constructed using Minda (49), a meta-caller integrating multiple VCFs to generate high-confidence consensus variants. This ensemble served as the definitive variant set for downstream analyses.

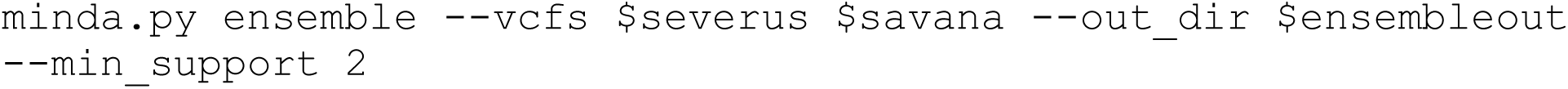

### Tumor purity and ploidy

The Wakhan (v0.2.0) tool was applied to integrate somatic SVs with haplotagged alignments and phased variants, providing insight into allele-specific SV patterns.

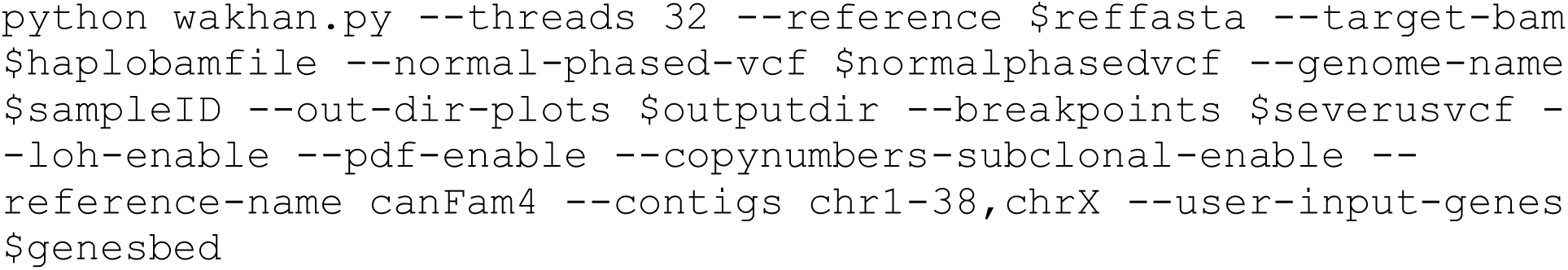

### Detection of chromothripsis, chromoplexy, and kataegis events

To systematically detect and classify three types of genomic catastrophe events: chromothripsis-like events, chromoplexy-like events, and kataegis-like events from tumor sequencing data.

#### Input data requirements

The pipeline requires structural variant (SV) calls in VCF format, copy number (CN) segment data (TSV, CSV, or VCF format), and gene annotation data (GTF format or pre-computed gene-overlap files). For kataegis detection, single nucleotide variant (SNV) and indel data in VCF format are optionally provided.

Chromosome nomenclature was standardized across all input files using a normalization procedure that either adds or strips the “chr” prefix, ensuring consistent chromosome identifiers throughout the analysis.

#### Chromothripsis detection

Chromothripsis detection utilized SV breakpoint clustering combined with CN oscillation patterns. The pipeline implements multiple detection models: a legacy three-boolean criterion model, (12) “minimal” and “full” (localized template assembly-like) chromothripsis classification schemes, and a unified confidence scoring system.

CN segments were optionally rounded to integer values before oscillation analysis. Two oscillation metrics were available: a “two-state” metric that evaluates transitions between distinct CN states, and an “adjacent” metric that examines consecutive segment pairs. The minimum run length criterion for oscillation was set (default: 7 segments), and consecutive segments with equal CN values could be compressed before oscillation counting. A tolerance parameter (cn_eps) was applied to treat CN values within a specified threshold as equivalent after rounding. For each detected cluster, the pipeline calculated breakpoint density (breakpoints per megabase), identified gene overlaps using the provided gene annotations, evaluated whether the event exhibited localized template assembly (LTA) characteristics, and assigned confidence levels (no-evidence, low-confidence, mid-confidence, high-confidence) based on the strength of chromothripsis features.

#### Chromoplexy detection

Chromoplexy events were detected using a graph-based approach that identifies chains of interconnected SVs spanning multiple chromosomes. The pipeline constructed an SV edge table from the input VCF, built a graph representation of SV connections, and identified connected components representing potential chromoplexy chains. CN segment data were integrated to assess the copy number context of detected chains. Each chromoplexy component was assigned a confidence score based on chain length, chromosomal complexity, and consistency with chromoplexy hallmarks.

### Structural variant formation event types

SVs were analyzed using a custom workflow that integrates breakpoint-level transposable-element (TE) annotations, non-TE homology matching, microhomology quantification, and templated-insertion detection to infer the most probable mutational mechanism for each event. The pipeline accepts SVs encoded in a VCF file containing standard SVTYPE annotations (DEL, DUP, INV, INS, BND). For INS and BND events where END coordinates may be missing, the breakpoint position is used as both POS and END, ensuring such events are retained. When available, the script preferentially uses the DETAILED_TYPE INFO field; otherwise SVTYPE values are normalized (e.g., DUP:TANDEM → DUP).

Breakpoints used for TE and homology annotation were provided as a BED7 file containing chromosome, start, end, SV identifier, coarse SV class, source label, and side designation (“start” or “end”).

#### Transposable element annotation of SV breakpoints

TE overlap was determined by intersecting breakpoint intervals with a RepeatMasker-formatted BED file using bedtools intersect. The RepeatMasker “name” field was parsed to extract TE family, class, and strand orientation. For breakpoints overlapping multiple TEs, the TE with the maximum overlap length was retained for scoring.

For each SV, TE annotations from the “start” and “end” breakpoints were paired by SV identifier. TE families, strand orientations, and overlap lengths were propagated into downstream scoring functions used to identify TE-mediated non-allelic homologous recombination (NAHR).

#### Annotation of Non-TE homologous Pairs

When provided, non-TE homology was supplied as a BEDPE file describing matched pairs of homologous genomic loci (e.g., segmental duplications). The BEDPE file was converted to a single-interval BED format with pair identifiers and strand relationships encoded in the feature name field. Each breakpoint was intersected with these intervals (left-outer join), and the best-overlapping homology annotation for each breakpoint side was retained.

Paired breakpoint annotations were compared to determine whether both breakpoints mapped to the same homology pair, the shared strand orientation (direct vs. inverted), and the maximum percent identity across the two sides.

#### Microhomology and templated-insertion analysis

Microhomology at the SV junction was computed using local reference-sequence queries to an indexed FASTA file (via pysam). For each SV, up to 50 bp upstream of POS and 50 bp downstream of END were compared to identify the longest exact microhomologous tract.

Templated-insertion length was extracted from VCF INFO fields (SVINSLEN or INSLEN), with robust parsing to accommodate integer or floating-point encodings. Missing insertion lengths were treated as zero.

#### Mechanism scoring framework

For each SV, three independent lines of evidence were evaluated:

- TE-mediated NAHR (TE family match + orientation rules)
- Non-TE NAHR (shared homology pair + strand and identity)
- Junction-based mechanisms (microhomology and insertion size)
- Each category contributes a score reflecting its support for specific mechanisms.

#### TE-mediated NAHR scoring

TE-NAHR was evaluated only when both breakpoints overlapped TEs of the same family. Expected strand relationships were required:

- DEL/DUP: concordant TE strands
- INV: opposite TE strands

Scores were increased by greater TE overlap. If the TE family matched but orientations were inconsistent with NAHR, the event was labeled TE_pair_other.

#### Non-TE NAHR scoring

Homology-mediated NAHR was considered only when both breakpoints mapped to the same homology pair. Percent identity modulated the score. Direct (+) orientation supported DEL/DUP, whereas inverted (–) orientation supported INV.

#### Junction-based mechanism scoring

Microhomology length and templated-insertion size were used to score canonical repair mechanisms:

- FoSTeS/MMBIR-like: ≥2 bp microhomology and ≥20 bp insertion
- MMEJ-like: 2–25 bp microhomology and ≤10 bp insertion
- NHEJ-like: ≤1 bp microhomology and ≤10 bp insertion

Events not meeting any criteria were classified as Unclassified.

#### Mechanism selection

All evidence categories were scored independently. The final mechanism call was chosen using a highest-score-wins strategy. Ties were resolved by alphabetical order of tied labels. The full score map, along with TE-based, non-TE-based, and junction-based reasoning strings, was recorded for auditability.

### Repetitive elements enrichment analysis

All analyses were performed on a high-performance computing (HPC) cluster managed by SLURM. Each job was executed on a single compute node using 64 CPU cores and 256 GB RAM. The environment was managed using a dedicated Conda environment, ensuring reproducibility of all Python- and bedtools-based analyses. All intermediate processing steps were executed within sample-specific output directories, using a temporary directory (TMPDIR) isolated per sample.

#### Structural variant pre-processing

SV calls for each sample were supplied as VCF files. Before analysis, VCFs were cleaned to remove non-standard or unplaced contigs and extraneous metadata lines. We excluded entries mapping to Super-Scaffold_3465, KP081776.1, and CM016470.1, following the reference genome filtering standards of the project.

Copy-number (CN) profiles generated by Savana were converted to BEDGRAPH format and similarly filtered to retain only autosomal and assembled chromosomes. These CN data were later used to correct for genomic biases in the permutation-based enrichment tests.

#### Breakpoint extraction and definition of uncertainty windows

SV breakends were extracted from each VCF using a custom Python parser (Step2.1.vcf_to_breakbends.py). The parser reports 1-bp breakend intervals for all SV classes (DEL, DUP, INV, INS, BND), incorporating POS and END when present and respecting PASS filters.

Filtered breakends were expanded into genomic “uncertainty windows” using bedtools slop, applying a ±200 bp window for SVs. This window length provides conservative inclusion of breakpoint uncertainty in the absence of CIPOS/CIEND fields and is chosen to be consistent with previous SV breakpoint enrichment frameworks.

Resulting windows were restricted to accessible genomic regions using an externally defined accessibility mask, ensuring all downstream randomization respected regions of mappable and callable reference sequence.

#### Detection of observed TE–breakpoint overlap

To quantify observed breakpoint–TE intersections, we overlapped accessible SV breakpoint windows with a RepeatMasker annotation BED file using bedtools intersect. For each overlapping event, RepeatMasker metadata fields were parsed to extract TE family, class (LINE, SINE, LTR, DNA), and repName. A summary of observed TE families intersecting SV breakpoints was generated for quality control and for guiding TE-level enrichment tests.

#### Permutation-based TE enrichment analysis

We quantified enrichment of TE sequence at SV breakpoints using a permutation framework implemented in Step2.2.permute_enrichment_v8.py. This method evaluates:

- Enrichment of breakpoints in any TE,
- Enrichment by TE class (e.g., LINE, SINE, LTR, DNA),
- Enrichment by TE family (e.g., L1, SINEC_Cf),
- Enrichment by specific TE names/repNames.

#### Randomization framework

Randomized breakpoints were generated using bedtools shuffle with the following constraints:

- Chromosome-preserving shuffling (-chrom) to maintain chromosome-level breakpoint density.
- Restriction to accessible genome only (-incl), matching the filtering applied to real breakpoints.
- Preservation of interval lengths (±200 bp windows).
- Exclusion of reference gaps and other uncallable regions.
- A total of 10,000 permutations per sample were performed (seed = 13) using 64 parallel threads.

#### Bias correction

To mitigate genomic covariates influencing TE density, the permutation model incorporates stratified shuffling across GC-content bins (n=3), mappability bins (n=3), and copy-number bins from Savana CN tracks (n=4). The codebase supports optional centromere-proximity bins, but these were not applied here because high-confidence centromere coordinates are not available for canFam4.

This matched-covariate randomization ensures that enrichment statistics are not confounded by heterogeneous genomic landscapes, including the variable mappability characteristic of repetitive sequences.

#### Significance testing

For each TE category with ≥1 observed overlap (default behavior), the following were computed:

- Observed overlap count
- Mean and 95% CI of overlaps across permutations
- Z-scores
- Empirical permutation p-values
- Benjamini–Hochberg FDR-adjusted q-values

An enrichment was considered significant when q < 0.05. Null distributions for “any TE” enrichment tests were optionally saved for visualization.

### Methylation analysis

We used haplotagged bam files with pb-cpg-tools (v3.0.0) for methylation analysis. pb-CpG-tools provides the tool aligned_bam_to_cpg_scores, which can generate site methylation probabilities from mapped HiFi reads, including probabilities for each haplotype when reads are haplotagged.

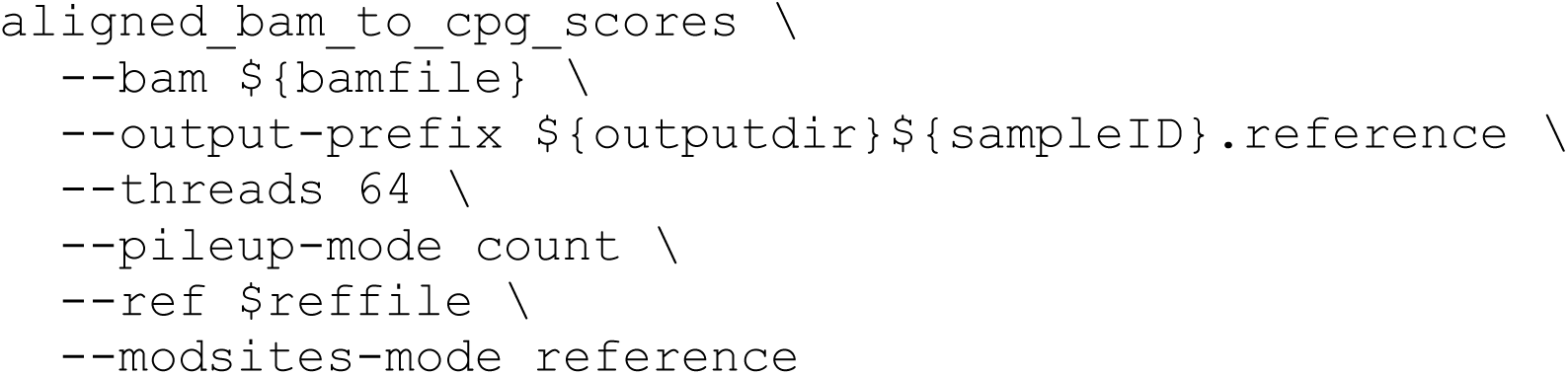

Next, we processed the results using the following step-by-step approach.

#### Input data and preprocessing

CpG methylation data were processed using pb-CpG-tools v3 counts. For each sample (blood, tumor, and optional normal tissue), we read the output files containing per-CpG methylated and unmethylated read counts. Only genomic coordinates and count fields were retained. BED inputs were treated as 0-based, half-open intervals, while GTF/GFF inputs were converted from 1-based closed to 0-based half-open coordinates.

To ensure base-level comparability across samples, CpG sites were merged using exact coordinate matching. Sites failing to meet minimum coverage thresholds (default ≥10 reads per sample) were excluded from downstream testing.

### Differentially methylated cytosine (DMC) calling

#### Blood-only mode

In settings where only blood was available, or when explicitly requested, we performed single-sample DMC calling using an exact binomial test against a user-specified null methylation probability (p-naught equals 0.5 by default). For each CpG, we calculated the methylation proportion beta equals mod divided by the quantity mod plus unmod, and evaluated deviation from the null. Resulting p-values were FDR-corrected using the Benjamini–Hochberg method. CpGs with absolute value of delta beta greater than or equal to 0.20 and q less than or equal to 0.05 were labeled as significantly hypo- or hyper-methylated.

##### Two-sample contrasts

For pairwise comparisons—blood vs tumor (T–B), blood vs normal (N–B), or tumor vs normal (T–N)—CpG sites present in both samples were tested using Fisher’s exact test on 2 by 2 tables of methylated vs unmethylated counts. Effect sizes were defined as delta beta equals beta subscript 2 minus beta subscript 1, and significance was assessed using FDR-corrected p-values. Significant CpGs (absolute value of delta beta greater than or equal to 0.20, q less than or equal to 0.05) were classified as hyper- or hypo-methylated relative to the first sample. All statistical tests were parallelized when requested, and used conservative handling of edge cases (e.g., zero-coverage rows → p = 1.0).

#### Differentially Methylated Region (DMR) Construction

Significant CpGs from each contrast were clustered into DMRs using a greedy genomic clustering algorithm. CpGs appearing within less than or equal to 250 bp of each other (default) were merged into the same DMR when all contributing CpGs showed methylation changes in the same direction (hyper or hypo).

Clusters were retained if they contained greater than or equal to 3 CpGs and the mean absolute value of delta beta exceeded a user-defined threshold (typically greater than or equal to 0.20). Each DMR was annotated with:

- genomic span
- CpG count
- mean delta beta
- minimum q-value
- hyper/hypo direction
- DMRs were exported in BED and table formats.

#### Triad-Based Tumor-Intrinsic Methylation Analysis

When blood (B), tumor (T), and matched normal tissue (N) were all available, we performed a triad analysis integrating the T–B, T–N, and N–B contrasts.

#### Triad merging and metric computation

For each CpG appearing in any contrast, values from all contrasts were merged. We computed three tumor-specificity metrics:

- comp_ratio equals absolute value of delta beta N-B divided by absolute value of delta beta T-B - Quantifies the fraction of tumor–blood difference explainable by normal tissue.
- tumor_frac equals absolute value of delta beta T-N divided by absolute value of delta beta T-B - Estimates the proportion of tumor–blood effect originating from tumor tissue.
- Tumor Specificity Index (TSI) equals the quantity absolute value of delta beta T-N minus absolute value of delta beta N-B, divided by the quantity absolute value of delta beta T-N plus absolute value of delta beta N-B - Ranges from negative 1 to 1, with positive values favoring tumor-specific alterations.

#### Classification of CpG sites

Each CpG was assigned to one of four categories based on significance patterns, effect sizes, and the above metrics:

- Tumor-intrinsic hyper/hypo (high-confidence T–N signal, low normal contribution, strong tumor fraction or TSI)
- Blood-driven changes (normal–blood differences dominate over tumor–normal)
- Mixed effects (comparable contributions of tumor and normal or discordant directions)
- Insufficient evidence (failure to meet thresholds)

High-confidence tumor-intrinsic CpGs required absolute value of delta beta T-N greater than or equal to 0.30 and q T-N less than or equal to 0.01, unless otherwise specified.

#### Tumor-intrinsic DMRs

If requested, tumor-intrinsic CpGs were clustered (TN-anchored) into final tumor-intrinsic DMRs using the same clustering strategy as above.

#### Genomic element and structural variant annotation

We supplied genomic annotations (BED/GTF) (canFam4) and structural variants (BED, BEDPE, or VCF). All DMRs were annotated with their minimum distance to each feature class, using a sweep-line interval matching algorithm optimized for large genome-wide sets.

Distances were classified as:

- Overlap (distance = 0)
- Proximal (0 < distance ≤ 1 kb; user-configurable)
- Distant/none (>1 kb or no nearby interval)

For BEDPE structural variants, both breakpoints were treated as independent 1-bp intervals. An annotated DMR table was produced containing all distances across all feature types and SVs.

### ecDNA detection

We used CReSIL (v1.2.0) (50) to detect ecDNAs on tumor tissue and, when available, matched blood and/or normal stromal controls.

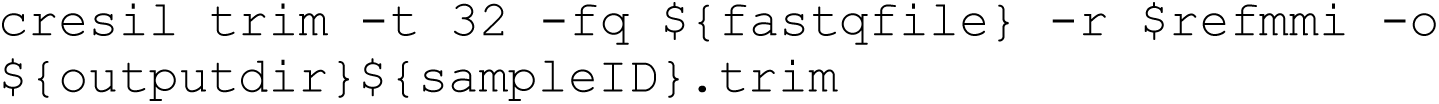

Then, we processed cresil ecDNA predictions using the following step-by-step process. CReSIL output tables require the fields id, merge_region, and merge_len, with additional columns (e.g., numreads, coverage, totalbase, ctc) incorporated when present. Tumor data were required for all analyses, whereas blood and normal controls were optional. All tables were imported using a delimiter-agnostic parser that supports both plain text and gzip-compressed files.

For each CReSIL record, genomic intervals were parsed from the merge_region field, normalized, and merged into non-overlapping segments. A minimum ecDNA size threshold (default: ≥100 kb) was applied to filter out low-confidence or sub-threshold circles.

#### Cross-Sample Subtraction and Control Handling

To identify tumor-specific ecDNA, we applied a bin-based subtraction strategy. Each circle’s merged intervals were converted into genomic bins of user-specified size (default: 1 kb), enabling fast comparison of presence/absence across samples. Tumor circles were removed if any of their genomic bins overlapped those from available controls. Presence in controls (present_in_B, present_in_N) was recorded for all circles, regardless of whether the corresponding control sample was provided.

When blood and/or normal stroma were supplied, the pipeline also generated Blood-only and Normal-only structural callsets by subtracting genomic bins found in other available samples (tumor and the alternate control). These structural sets were produced irrespective of whether copy-number data were available for controls.

#### Tumor quality control metrics

Tumor-level quality control (QC) metrics were computed directly from CReSIL outputs. For each tumor circle, reads-per-kilobase (reads_per_kb) was calculated as numreads / (merge_len / 1000). Summary statistics including the total number of eccDNAs, total supporting reads, mean circle length, total bases, and average coverage were generated and written to {prefix}.tumor_qc_summary.txt. The top N tumor circles ranked by reads_per_kb (default: N=10) were reported in {prefix}.tumor_topReadsPerKb.tsv.

#### Copy-number integration and ecDNA tier classification

Sample-specific copy-number information was incorporated using bedGraph files for the tumor (required) and optionally for blood and normal controls. bedGraph files were standardized to four columns (chrom, start, end, CN) and indexed per chromosome for efficient overlap queries.

For each circle, copy-number metrics were computed by intersecting merged intervals with CN segments. Weighted mean CN, weighted median CN, maximum CN, the number of CN bins overlapped, and total base pairs intersected were calculated.

ecDNA were classified into three confidence tiers according to circle size and CN metrics:

- Tier 1 (High-confidence ecDNA): size ≥500 kb and median CN ≥4.
- Tier 2 (Probable ecDNA): size ≥100 kb with median CN ≥4, or size ≥100 kb with maximum CN ≥6.
- Below CN threshold: circles that did not meet Tier 1 or Tier 2 criteria.

Tumor CN-augmented tables were written to {prefix}.withCN.tsv. Control CN-augmented and high-confidence lists were produced only when control CN bedGraphs were provided.

#### Generation of final tumor ecDNA callset

The final tumor ecDNA set consisted of all Tier 1 and Tier 2 circles. For each final circle, merged genomic intervals were exported as BED format ({prefix}.final_mergedIntervals.bed). Equivalent outputs were generated for blood and normal controls whenever high-confidence calls were available.

#### Gene, promoter overlap annotation

Gene annotations were incorporated using either a user-supplied canFam4 GTF/refFlat file or by downloading the UCSC canFam4 RefSeq GTF. Gene models were collapsed per gene to obtain minimal genomic spans.

Promoter intervals were supplied as a BED file or generated de novo from transcription start sites (TSS), using an upstream/downstream window (default: −2000 bp / +200 bp). Enhancer annotations were optionally provided as BED files.

For each final ecDNA, merged intervals were intersected with genes, promoters, and enhancers. Analogous files were produced for blood and normal ecDNA when available.

#### Tumor cell frequency estimation

Tumor cell support for each ecDNA was calculated using one of two approaches:

- Direct per-circle cell support file (--cells-t), supporting formats:
- long form: id, cell_id
- counts: id, n_cells
- lists: id, supporting_cells

Per-cell overlap inference, using individual cell CReSIL outputs and Jaccard-based presence detection. A circle was considered present in a cell if the Jaccard index exceeded a threshold (default: ≥0.20) and overlapping base pairs exceeded a minimum (default: ≥10 kb).

Circle-level cell counts were optionally normalized by total profiled cells (--total-cells-t). When requested, a pre-filled template of circle IDs with zero counts was generated.

#### Summary reporting and output structure

All analysis steps generated a comprehensive summary file ({prefix}.summary.txt) detailing controls used, the number of tumor-only circles, Tier 1 and Tier 2 counts, promoter/enhancer hits, cell-support information, and all output file paths.

### ddPCR Validation

The CNV of CTNNB1 was verified using droplet digital polymerase chain reaction (ddPCR). A reaction mix of 20 ng gDNA, 11 ul of Supermix for probes (No dUTP) (Bio-Rad Laboratories), 1.1 ul customized canine CTNNB1 CNV assay (assay ID: dCNS572026389) (Bio-Rad Laboratories), 1.1 ul customized canine RPP30 CNV assay (assay ID: dCNS431780688) (Bio-Rad Laboratories), 5 U of HeaII diluted with CutSmart buffer (New England Biolabs) was added into well of 96-well ddPCR plate and brought up to 22 ul with nuclease-free water. The ddPCR plate was then heat sealed and ran on the Automated Droplet Generator with Droplet Generation Oil for Probes (Bio-Rad Laboratories) following manufacturer’s instructions. After droplet generation, the heat-sealed ddPCR plate with droplets was run on BioRad C1000 Deep-Well Touch Thermal Cycler for PCR (10 minutes at 95 °C, 40 cycles of 94 °C for 30 seconds and 60 °C for 1 minute, then 98 °C for 10 min and cooling to 4°C). Ideal annealing temperature was determined by the prior temperature gradient run using the MSC cell line. Droplets were analyzed using the QX200 Droplet Reader and the QuantaSoft software (Bio-Rad Laboratories). The copy number of CTNNB1 gene was calculated by QuantaSoft, normalized to RPP30 reference gene.

### PCR Validation Targets

To validate predicted breakpoints and junction points in ecDNA candidates, we performed PCR amplification across putative junction sites. For each ecDNA candidate, we designed primer pairs flanking three distinct breakpoints/junction points identified through long-read sequencing. Primers were designed to amplify across the predicted junctions, with one primer binding to each side of the breakpoint, such that successful amplification would only occur if the predicted junction was present in the sample.

PCR reactions were performed using Quick-Load® Taq 2X Master Mix (New England Biolabs) according to the manufacturer’s protocol. Each 25 μL reaction contained 1 ng of genomic DNA template, 1× PCR buffer and 0.2 μM of each primer. Thermal cycling conditions consisted of initial denaturation at 95°C for 60 seconds, followed by 40-45 cycles of 95°C for 30 seconds, 59-63°C for 30 seconds (annealing temperature optimized for each primer pair), and 68°C extension time (based on expected product size, 1 minute per kb), with a final extension at 68°C for 5 minutes. PCR products were visualized on 1% agarose gels stained with SYBR Safe DNA Gel Stain.

PCR product and sequencing primers were submitted for Sanger Sequencing (QuintaraBio) to confirm the precise breakpoint junctions. Sequencing results were aligned to the reference genome using Snapgene Viewer to verify that the junction sequences matched the predicted breakpoint coordinates. A breakpoint was considered validated if PCR amplification produced a product of the expected size in basepairs and Sanger sequencing confirmed the junction sequence spanning the predicted breakpoint.

### Optical genome maps de novo assembly - orthogonal validation

Single-molecule genome maps of GP1899 were *de novo* assembled using the assembly pipeline Bionano Solve v3.5, developed by Bionano Genomics with default settings using the command below. In short, a pairwise comparison of DNA molecules (min 250 kbp) was generated to produce the initial consensus genome maps. During an extension step, molecules were aligned to genome maps, and maps were extended based on the molecules aligning past the map ends. Overlapping genome maps were then merged. Extension and merge steps were repeated five times before a final refinement of the genome maps. Clusters of molecules aligned to genome maps with unaligned ends >30 kbp in the extension step were re-assembled to identify all alleles. Resulting files were used for chromothripsis and ecDNA validations.

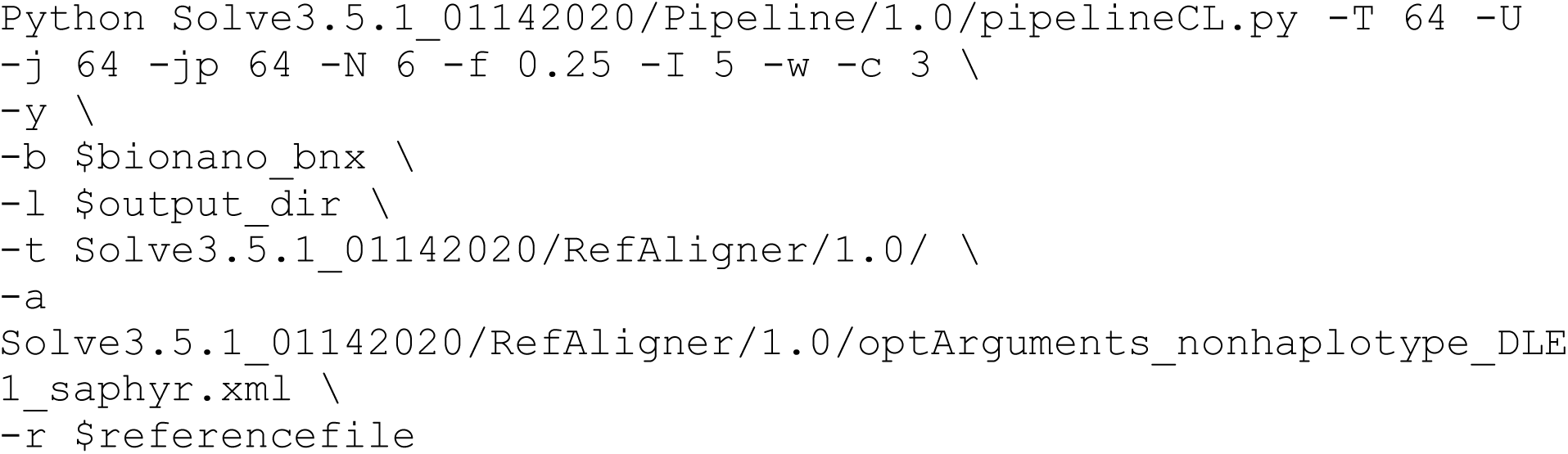

## Data, Materials, and Software Availability

All PacBio HiFi long-read sequencing bam files will be available under NCBI Project ID. Custom codes used are available under https://github.com/yilmazf/DogGenomesAnalyses.

## Acknowledgements

Funding was provided by the National Institutes of Health (NIH) grant U24HG007497 (to C.L. and F.Y.) and the Martin J. Gavin Endowment at Connecticut Children’s Medical Center (to C.L.L.). We sincerely thank Jeffrey J. Schoenebeck and Louise Van der Weyden for their thoughtful and critical review of the manuscript draft, which greatly strengthened the final version. We thank Jee Young Kwon, Leigh Maher, and Qihui Zhu for their assistance with the organization of canine samples. We gratefully acknowledge the Genome Technologies Service at The Jackson Laboratory for expert assistance with the work described herein. We also thank the Scientific Services at The Jackson Laboratory for their assistance with this work, and Research IT for providing computational infrastructure. Finally, we acknowledge the generous contributions of the dog owners who provided samples to Colorado State University, as well as the support of Colorado State University—especially Irene Mok and Dr. Susan Lana—in supplying the canine osteosarcoma materials used in this study.

## Author Contributions

C.L. and C.C.L. conceived and supervised the study. F.Y. designed and performed the computational analyses, including long-read sequence alignment, variant calling, structural variant detection, repeat-enrichment analyses, DNA methylation profiling, ecDNA identification, and data integration, and drafted the manuscript. S.A.S. performed PCR validation experiments and curated biosamples. J.D. performed digital droplet PCR validation of copy number alterations in ecDNAs. P.N., and F.H.O. contributed to data analysis. F.Y., S.A.S., F.H.O., P.N., C.C.L., and C.L. contributed to manuscript editing.

## Competing interests

C. L. is a scientific advisory board member of Nabsys and Genome Insight.

