## Supplementary Materials and Methods for "Satellite DNA fragility accompanies complex genome rearrangements and ecDNA oncogene amplification in canine osteosarcomas"

**Table of Contents**

Figures

### **Supplementary Figures**

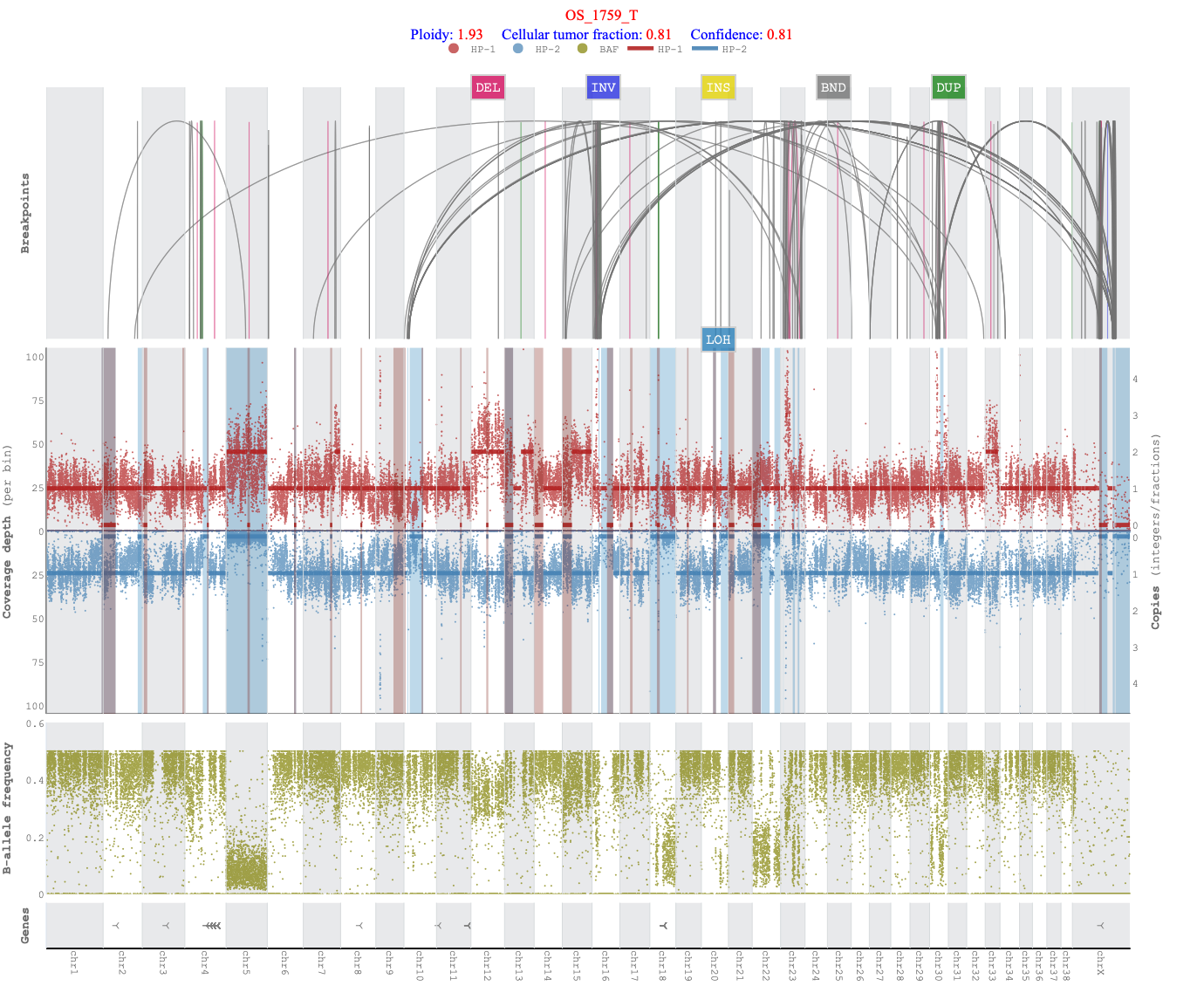
**Fig. S1.** Genome-wide copy number profile for sample Ro1759 (OS_1759_T). The figure displays three integrated panels showing chromosomal alterations across all chromosomes (1-38, X). Top panel: Structural variant arcs indicating genomic rearrangements, including deletions (DEL, pink), inversions (INV, purple), insertions (INS, yellow), translocations/breakends (BND, gray), duplications (DUP, green), and loss of heterozygosity (LOH, blue). Middle panel: Copy number profile showing coverage depth per genomic bin (red and blue dots representing different haplotypes or read pairs) with vertical bars indicating chromosome boundaries and shaded regions highlighting copy number alterations. Bottom panel: B-allele frequency plot (yellow dots) displaying allelic imbalance across the genome, with dense bands indicating regions of LOH. The sample exhibits a ploidy of 1.93 and a cellular tumor fraction of 0.81 (confidence: 0.81), indicating near-diploid tumor content with high purity. Alternating gray and white background bands demarcate individual chromosomes, with chromosome ideograms shown at the bottom.

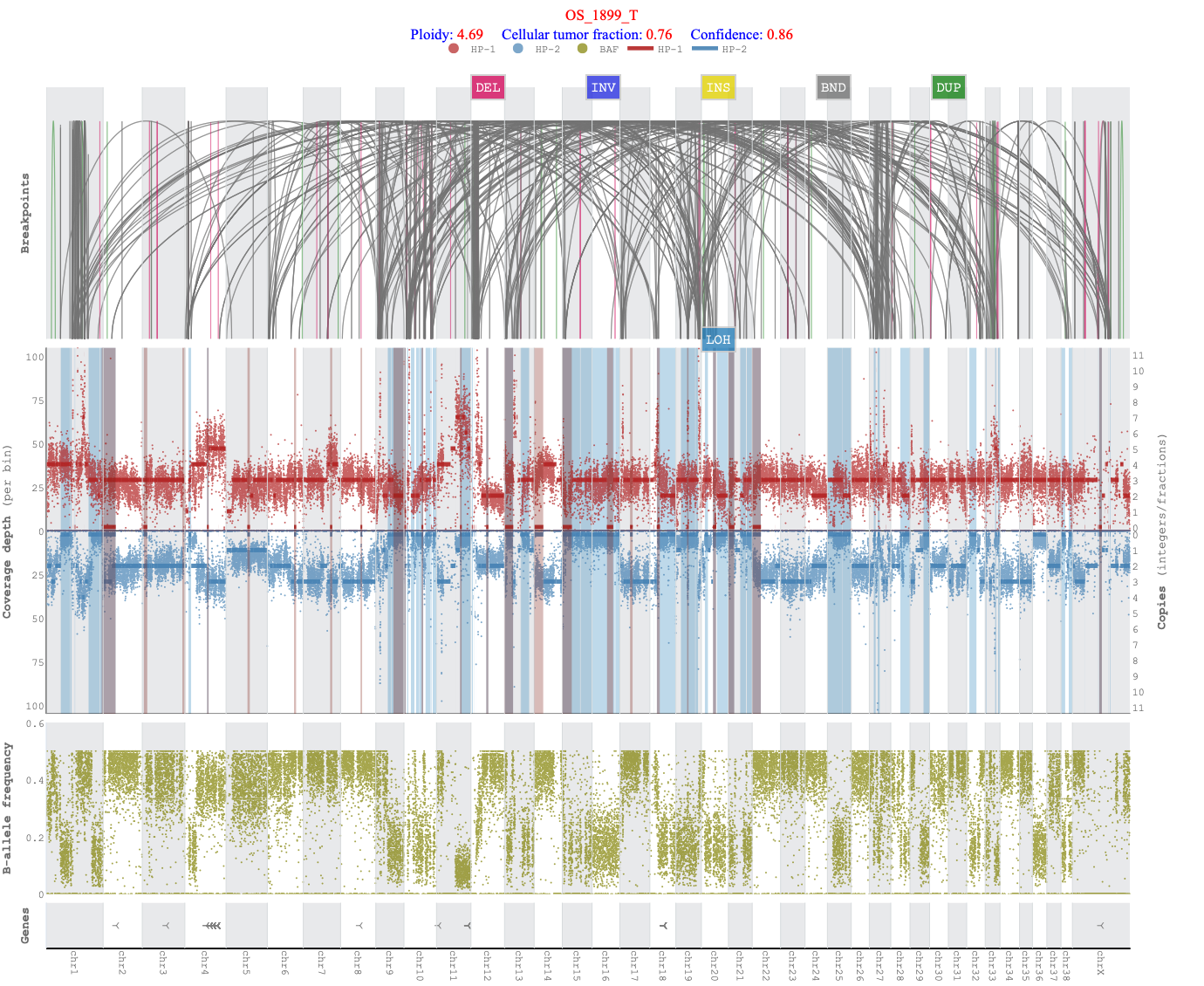
**Fig. S2.** Genome-wide copy number profile for sample GP1899 (OS_1899_T). The figure displays three integrated panels showing chromosomal alterations across all chromosomes (1-38, X). Top panel: Structural variant arcs indicating extensive genomic rearrangements, including deletions (DEL, pink), inversions (INV, purple), insertions (INS, yellow), translocations/breakends (BND, gray), duplications (DUP, green), and loss of heterozygosity (LOH, blue). The high density of interconnecting arcs reveals complex chromosomal instability. Middle panel: Copy number profile showing coverage depth per genomic bin (red and blue dots representing different haplotypes) with vertical bars indicating chromosome boundaries and shaded regions highlighting copy number alterations. The elevated coverage values (scale extending to 11 copies) reflect widespread genomic amplifications. Bottom panel: B-allele frequency plot (yellow dots) displaying allelic imbalance across the genome, with heterogeneous banding patterns indicating complex clonal architecture and regions of LOH. The sample exhibits a ploidy of 4.69 and a cellular tumor fraction of 0.76 (confidence: 0.86), indicating a highly polyploid tumor with substantial genomic complexity and high tumor purity. Alternating gray and white background bands demarcate individual chromosomes, with chromosome ideograms shown at the bottom.

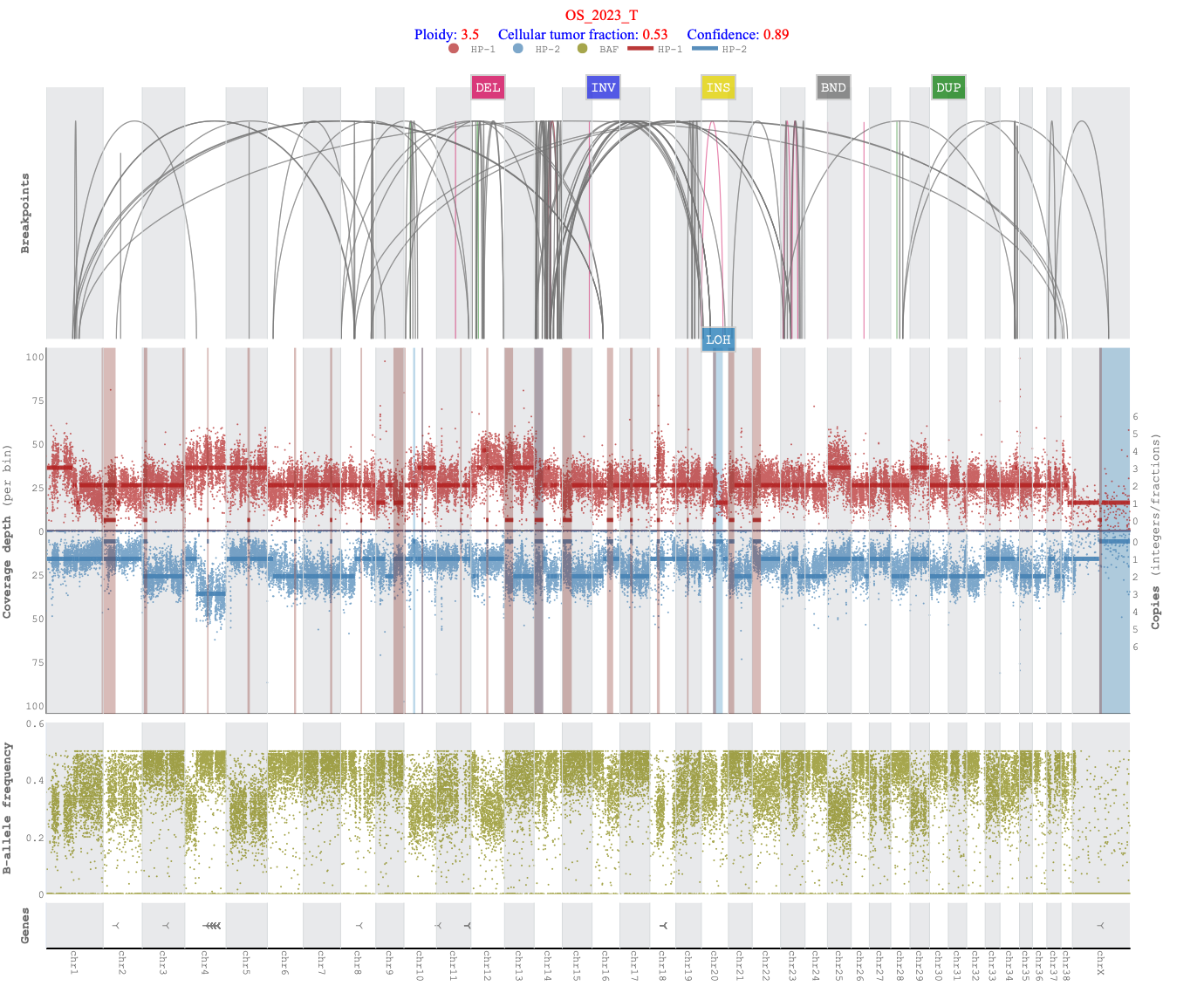
**Fig. S3.** Genome-wide copy number profile for sample GR2023 (OS_2023_T). The figure displays three integrated panels showing chromosomal alterations across all chromosomes (1-38, X). Top panel: Structural variant arcs indicating genomic rearrangements, including deletions (DEL, pink), inversions (INV, purple), insertions (INS, yellow), translocations/breakends (BND, gray), duplications (DUP, green), and loss of heterozygosity (LOH, blue). The interconnecting arcs reveal inter- and intra-chromosomal rearrangements throughout the genome. Middle panel: Copy number profile showing coverage depth per genomic bin (red and blue dots representing different haplotypes) with vertical bars indicating chromosome boundaries and shaded regions (pink and blue) highlighting copy number alterations. The profile demonstrates regional copy number gains and losses across multiple chromosomes. Bottom panel: B-allele frequency plot (yellow dots) displaying allelic imbalance across the genome, with banding patterns indicating regions of LOH and heterozygosity. The sample exhibits a ploidy of 3.5 and a cellular tumor fraction of 0.53 (confidence: 0.89), indicating a triploid/near-tetraploid tumor with moderate tumor purity and substantial normal cell contamination. Alternating gray and white background bands demarcate individual chromosomes, with chromosome ideograms shown at the bottom.

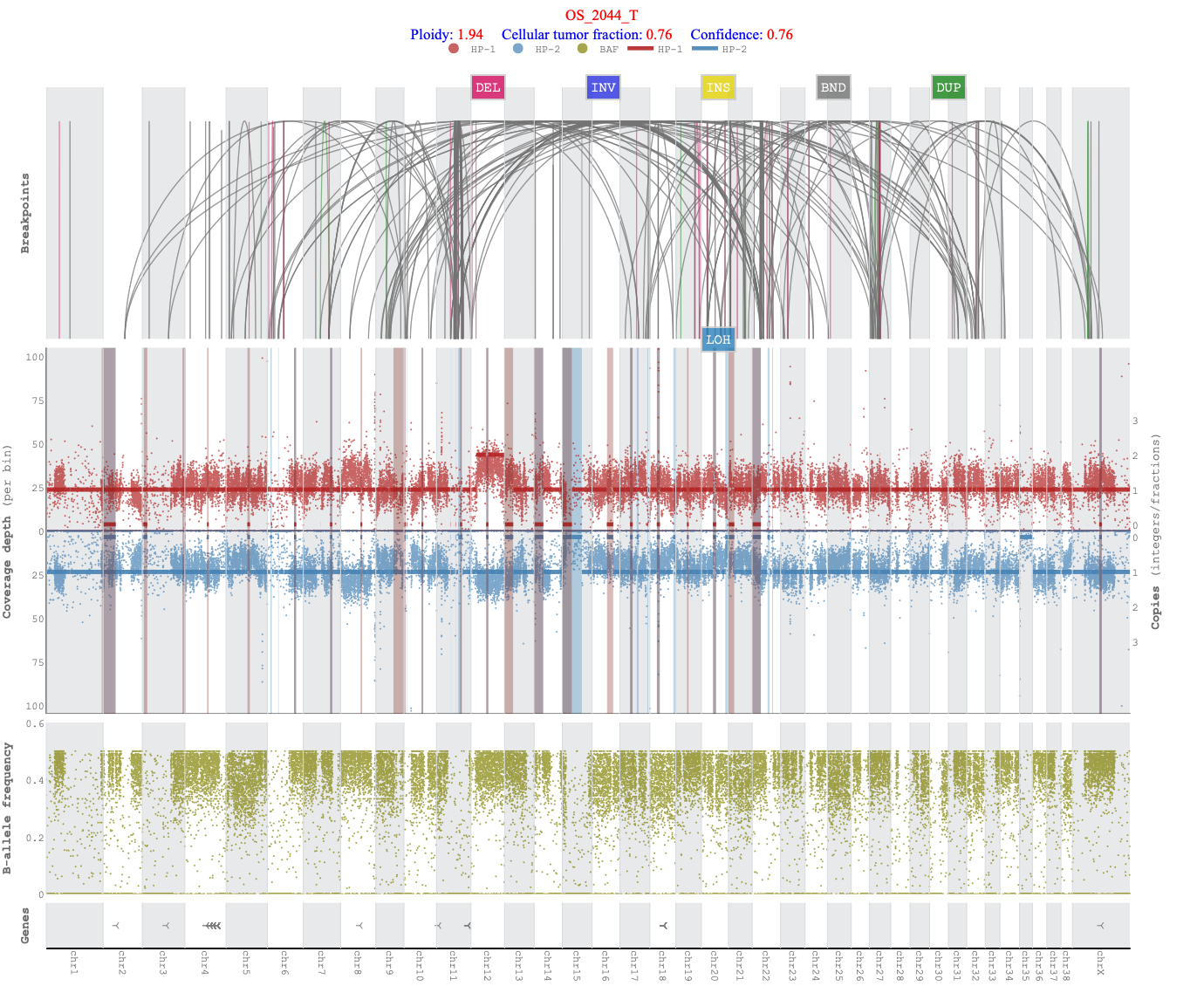
**Fig. S4.** Genome-wide copy number profile for sample GD2044 (OS_2044_T). The figure displays three integrated panels showing chromosomal alterations across all chromosomes (1-38, X). Top panel: Structural variant arcs indicating genomic rearrangements, including deletions (DEL, pink), inversions (INV, purple), insertions (INS, yellow), translocations/breakends (BND, gray), duplications (DUP, green), and loss of heterozygosity (LOH, blue). The interconnecting arcs reveal multiple inter- and intra-chromosomal rearrangements with notable complexity in the central chromosomes. Middle panel: Copy number profile showing coverage depth per genomic bin (red and blue dots representing different haplotypes) with vertical bars indicating chromosome boundaries and shaded regions (pink and blue) highlighting copy number alterations. The relatively stable baseline around diploid levels indicates moderate genomic stability with focal alterations. Bottom panel: B-allele frequency plot (yellow-olive dots) displaying allelic imbalance across the genome, with distinct banding patterns indicating widespread regions of LOH alternating with retained heterozygosity. The sample exhibits a ploidy of 1.94 and a cellular tumor fraction of 0.76 (confidence: 0.76), indicating a near-diploid tumor with high purity and minimal normal cell contamination. Alternating gray and white background bands demarcate individual chromosomes, with chromosome ideograms shown at the bottom.

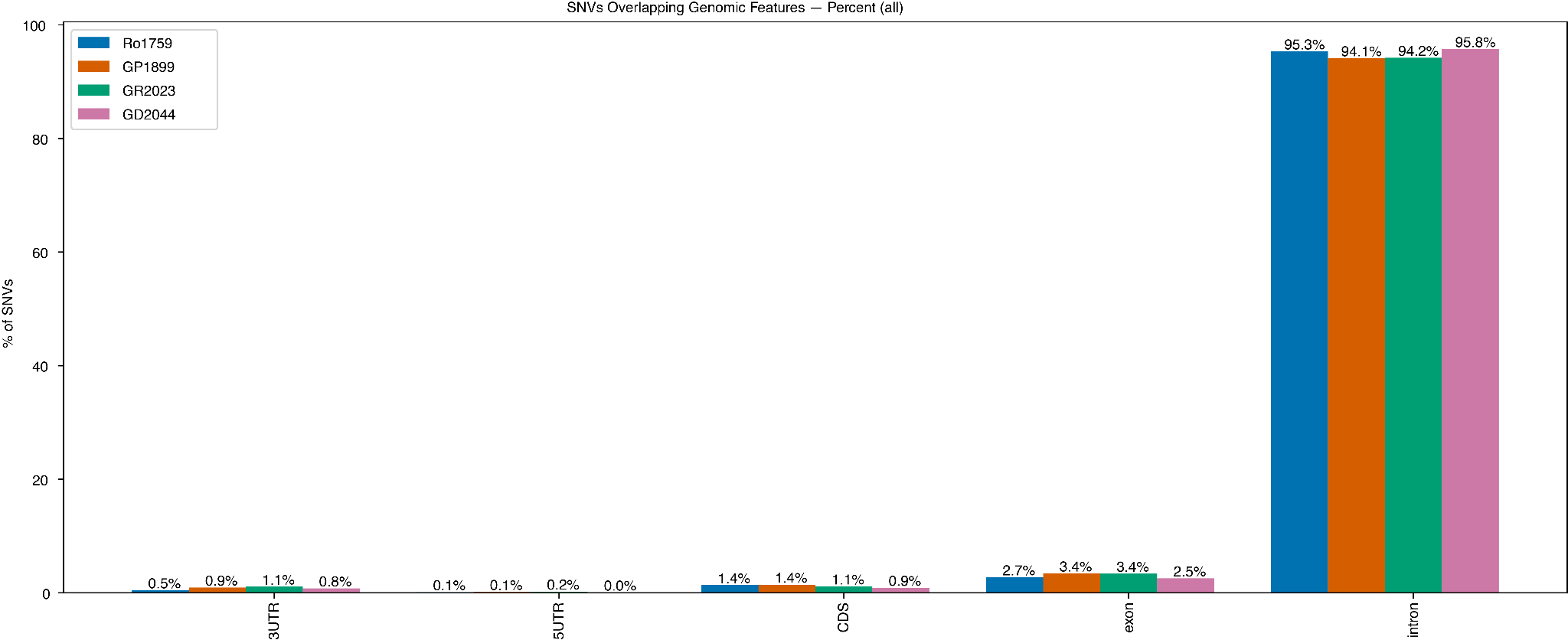
**Fig S5.** Distribution of single nucleotide variants (SNVs) across genomic features for four samples (Ro1759, GP1899, GR2023, and GD2044). The bar chart shows the percentage of SNVs overlapping with different genomic regions including 3' untranslated regions (3'UTR), 5' untranslated regions (5'UTR), coding sequences (CDS), exons, and introns. Across all samples, the majority of SNVs (94.1-95.8%) are located within intronic regions, while smaller fractions overlap with exons (2.5-3.4%), CDS (0.9-1.4%), 3'UTRs (0.5-1.1%), and 5'UTRs (0.0-0.2%). Percentage values are displayed above each bar.

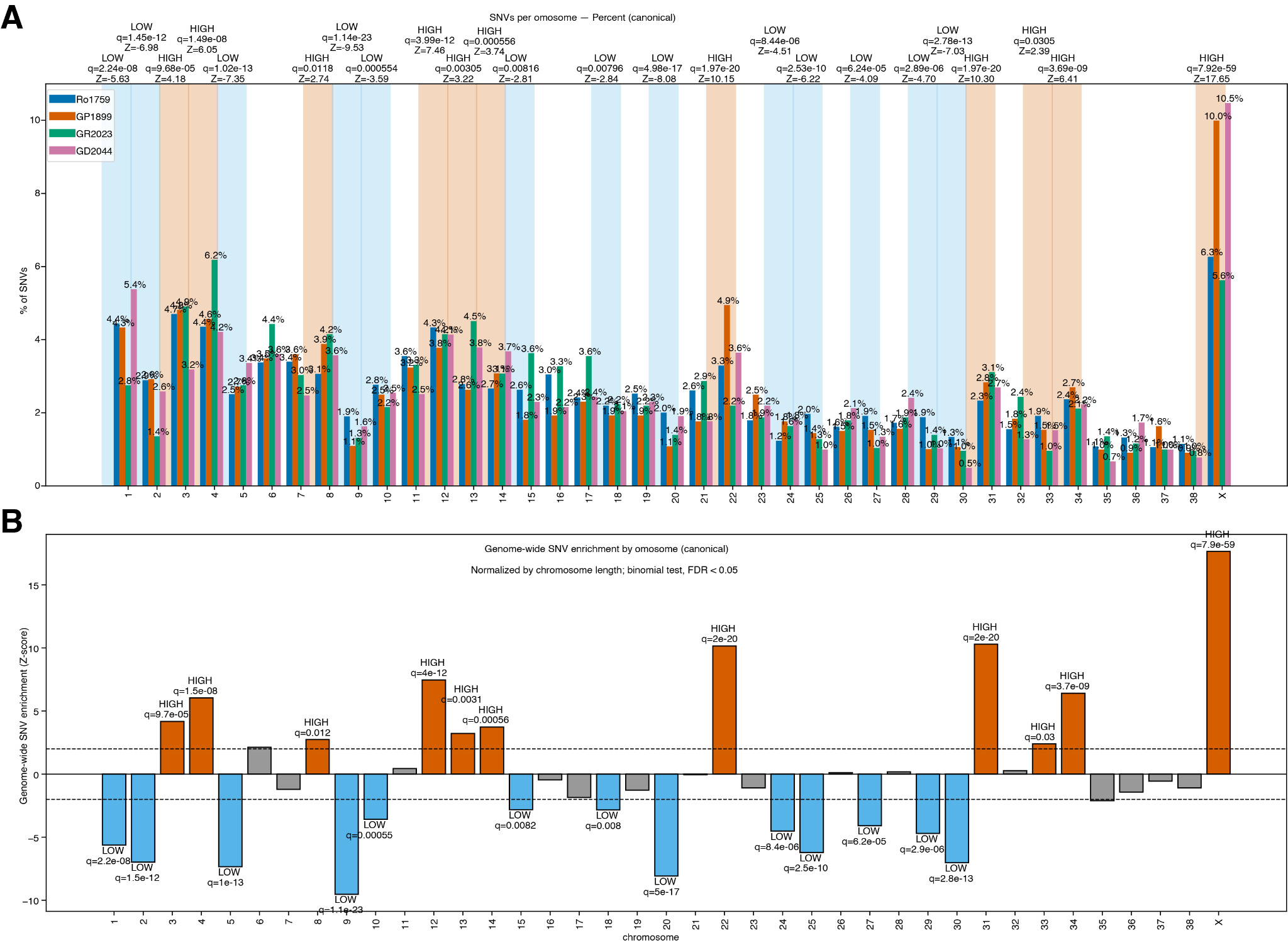

**Fig. S6.** Chromosomal distribution and enrichment of single nucleotide variants (SNVs) in cancer samples. (A) Percentage of SNVs per chromosome across four cancer samples (Ro1759, GP1899, GR2023, and GD2044). Bars represent the proportion of total SNVs mapping to each canonical chromosome (1-38, X). Background shading indicates chromosomes with significantly high (orange) or low (blue) SNV burden relative to chromosome length. Statistical significance (q-values and Z-scores) for enriched or depleted chromosomes is annotated above. The highest SNV percentage (10.5%) is observed on chromosome X in sample Ro1759. (B) Genome-wide SNV enrichment by chromosome normalized by chromosome length using a binomial test (FDR < 0.05). Positive Z-scores (orange bars) indicate chromosomes with significantly elevated SNV density (HIGH), while negative Z-scores (blue bars) indicate chromosomes with significantly reduced SNV density (LOW). Gray bars represent chromosomes without significant enrichment or depletion. Dashed horizontal lines indicate significance thresholds. Chromosome X shows the strongest enrichment (Z-score ≈ 17, q=7.9e-69), followed by chromosomes 22, 31, and 13. Chromosomes 1, 2, 5, 9, 19, 20, and 29 show significant depletion of SNVs.

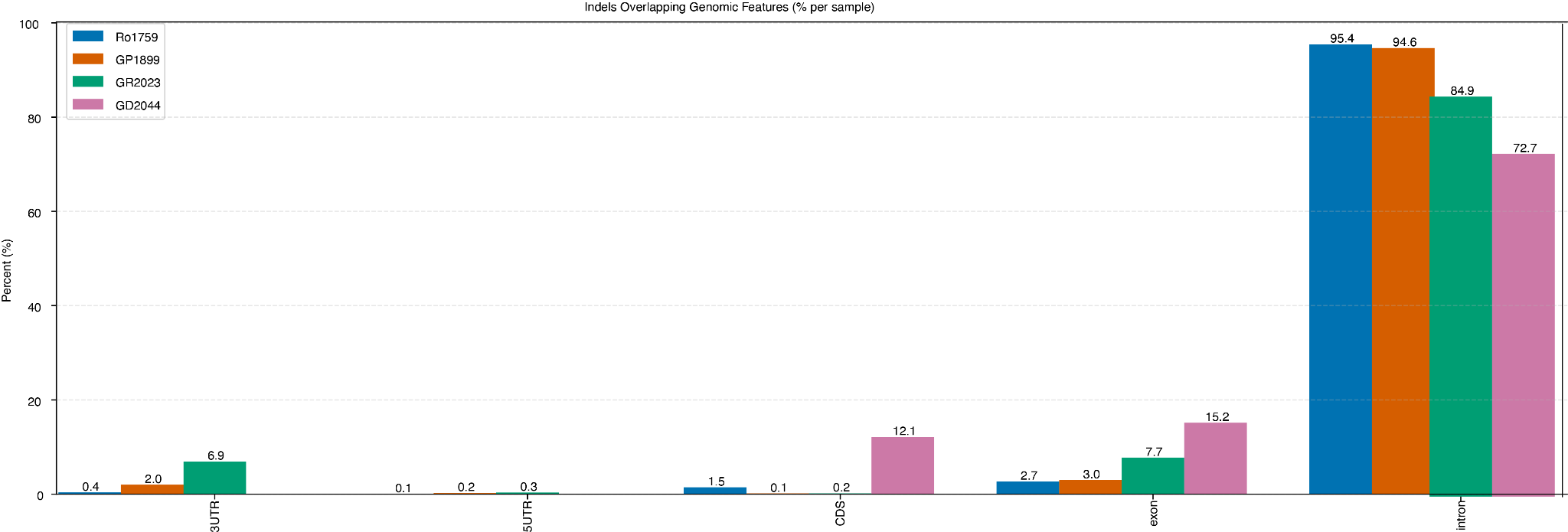

**Fig. S7.** Distribution of indels across genomic features in four samples. Bar chart showing the percentage of indels per sample that overlap with different genomic features (3' UTR, 5' UTR, coding sequence (CDS), exons, and introns) for four samples: Ro1759 (blue), GP1899 (orange), GR2023 (green), and GD2044 (pink). The majority of indels (72.7-95.4%) overlap with intronic regions across all samples, while smaller percentages are found in exons (2.7-15.2%), CDS (0.1-12.1%), and UTR regions (<7%). Sample GD2044 shows notably lower intronic overlap (72.7%) and higher exonic (15.2%) and CDS (12.1%) overlap compared to other samples.

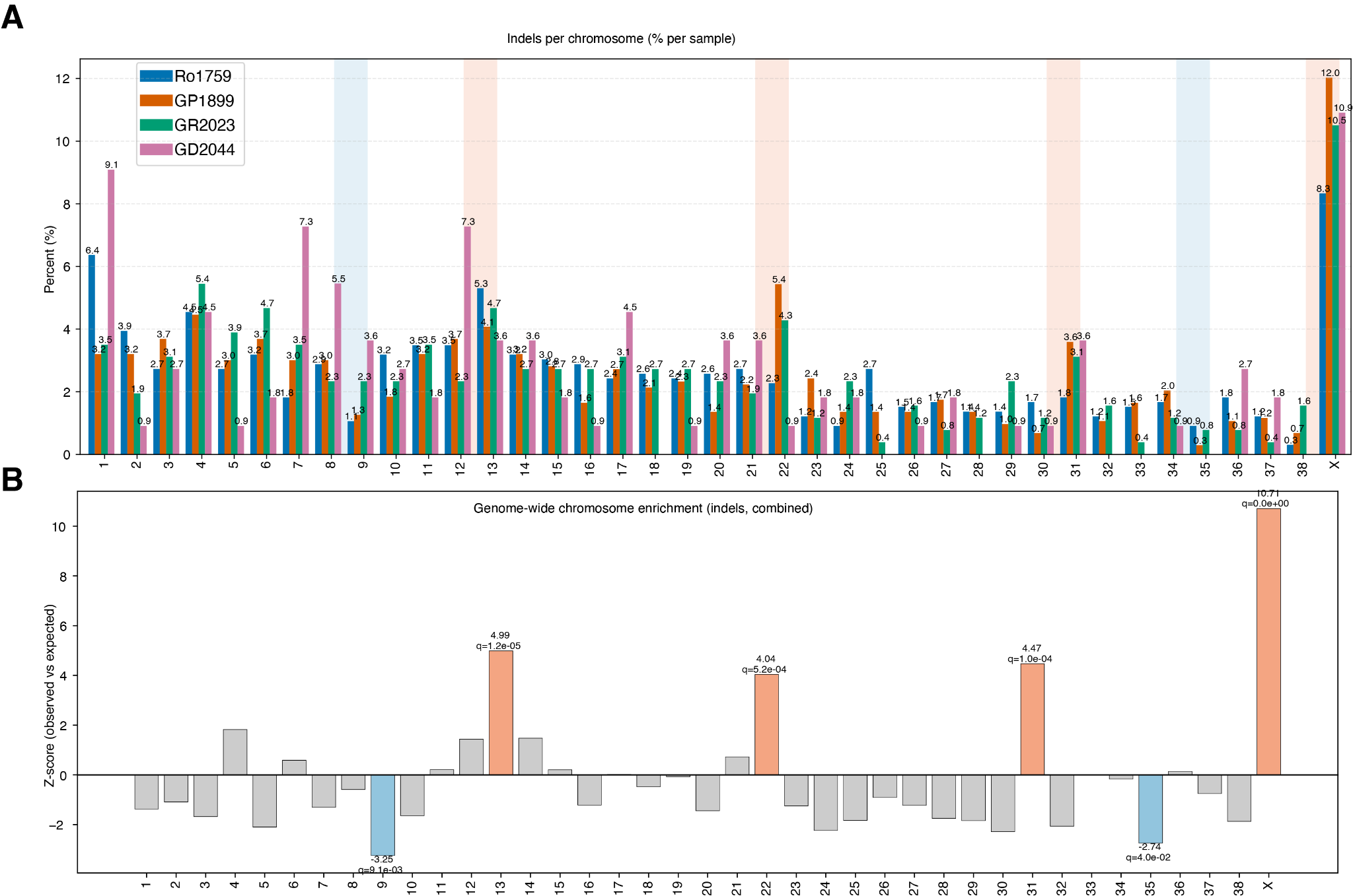

**Fig. S8.** Distribution and enrichment of indels across chromosomes. (A) Chromosome-specific indel frequencies shown as the percentage of total indels per sample for four samples: Ro1759 (blue), GP1899 (orange), GR2023 (green), and GD2044 (pink). Chromosomes 1, 7, 12, 22, and X show elevated indel frequencies across samples, with chromosome X displaying the highest percentage (8.5-12.0%). (B) Genome-wide chromosome enrichment analysis for indels across all samples combined. Z-scores represent the deviation of observed indel counts from expected values. Chromosomes with significant enrichment (orange bars; q<0.05) include chromosomes 13, 22, 31, and X, with X showing the most pronounced enrichment (Z-score=10.71, q=0e+00). Most other chromosomes show slight depletion (gray bars, negative Z-scores). Q-values indicate false discovery rate-corrected p-values.

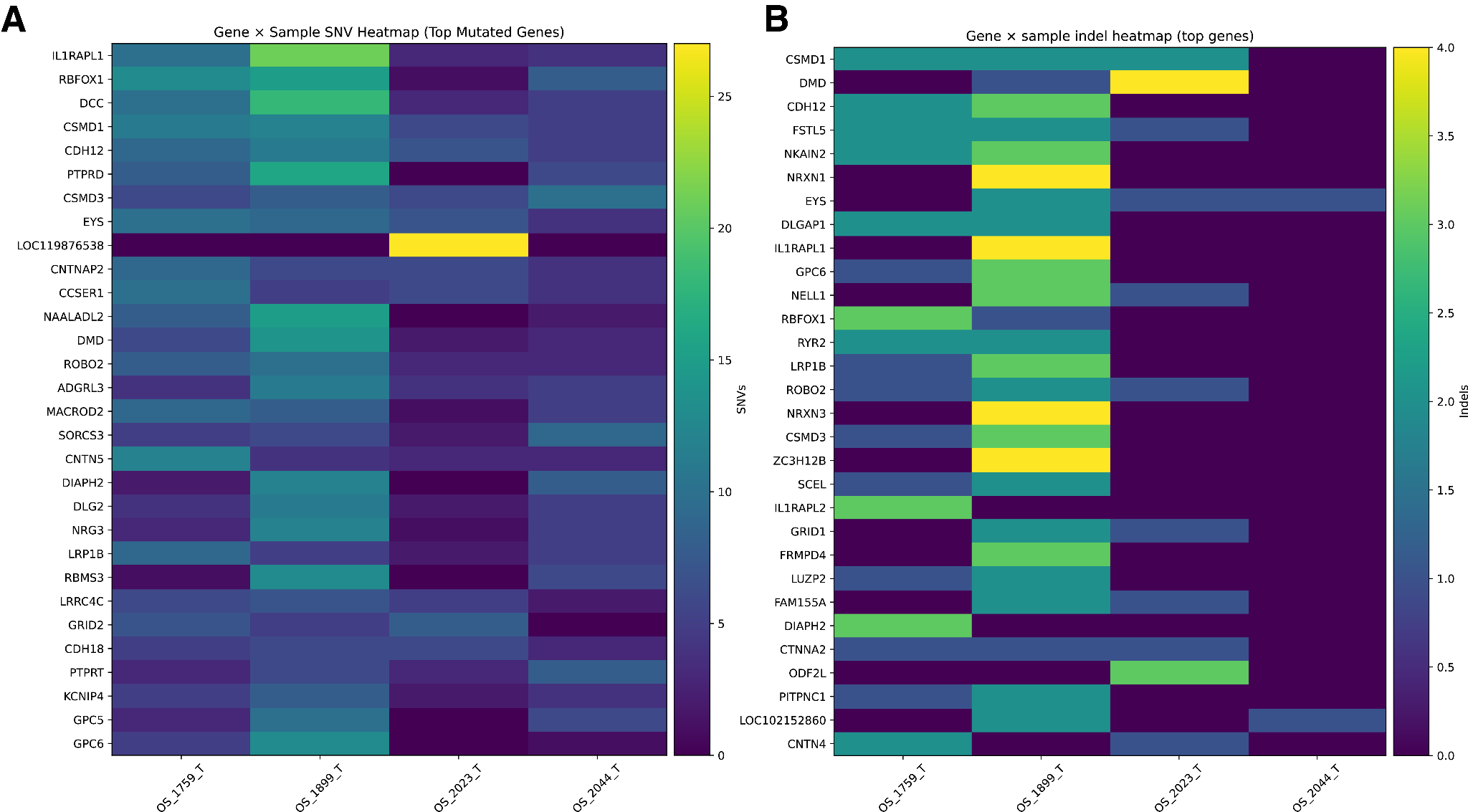

**Fig. S9.** Mutation landscape of the most frequently mutated genes across four samples. **(A)** Heatmap showing the distribution of single nucleotide variants (SNVs) in the top mutated genes. Each cell represents the number of SNVs (color scale: 0-30) in a given gene (rows) for each sample (columns). LOC119876538 shows the highest SNV count (n=28) in 2023, while genes such as IL1RAPL1, RBFOX1, and DCC exhibit moderate to high SNV burdens across multiple samples. 1899 and 2023 generally show higher SNV counts compared to other samples. **(B)** Heatmap displaying the distribution of insertions and deletions (indels) in the top mutated genes (color scale: 0-4). DMD, NRXN1, NRXN3, and ZC3H12B show the highest indel counts (n=4) in 1899. Several genes including CSMD1, FSTL5, and NKAIN2 show indels across multiple samples, though at lower frequencies than SNVs. The distinct mutation patterns between samples suggest inter-sample heterogeneity, while certain genes (e.g., CSMD1, RBFOX1, IL1RAPL1) appear in both panels, indicating they harbor multiple types of variants.

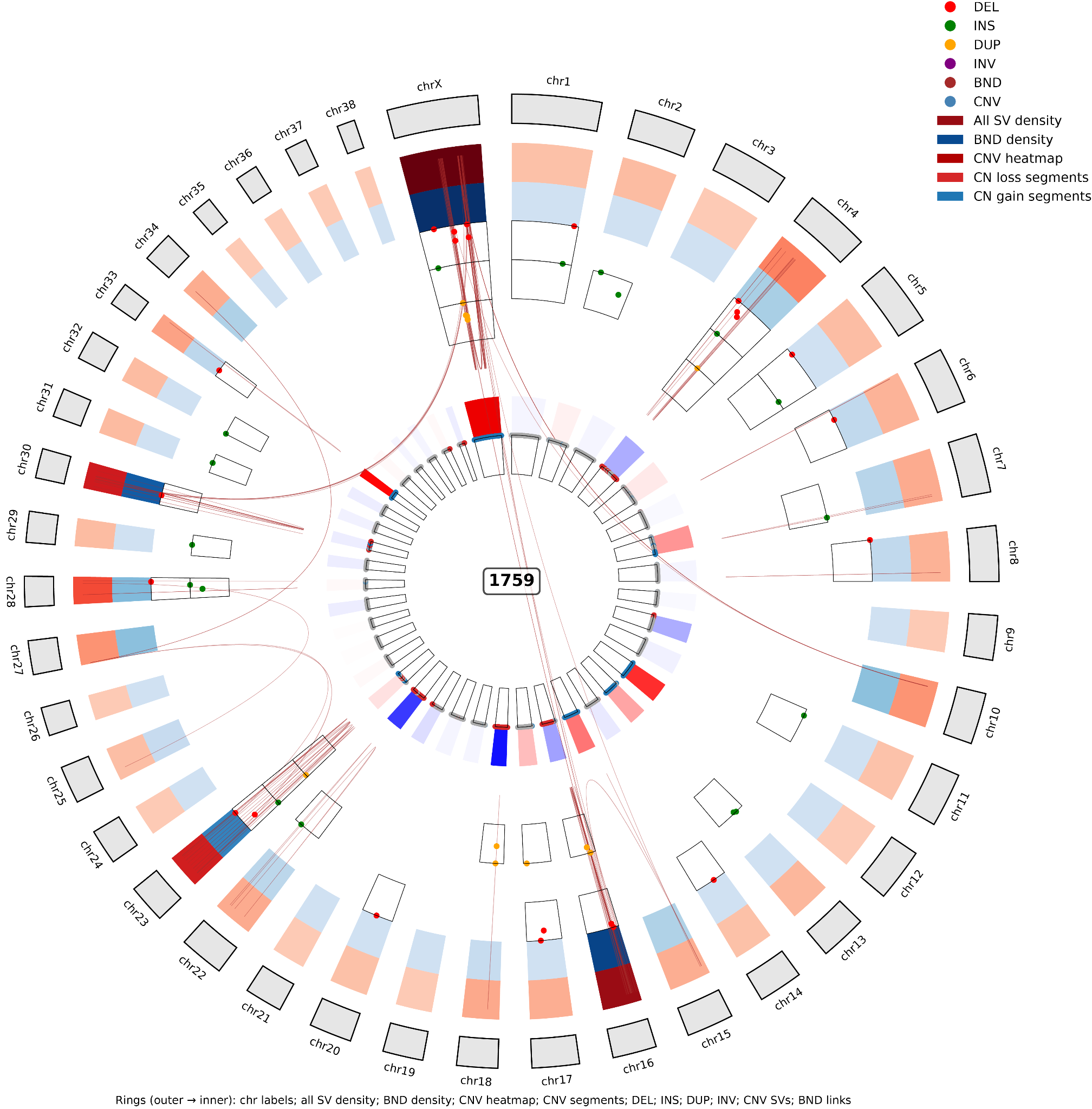
**Fig. S10.** Circos plot displaying the genomic landscape of structural variants and copy number alterations in sample Ro1759. The circular plot represents the entire genome with chromosome labels around the periphery. From outer to inner, the concentric rings display: (1) chromosome ideograms with cytogenetic band positions, (2) all structural variant (SV) density shown as a dark red heatmap, (3) breakend (BND) density displayed in dark blue, (4) copy number variation (CNV) heatmap with gains in red/orange and losses in blue, (5) specific copy number gain segments (red bars), and (6) copy number loss segments (blue bars). Individual SVs are marked on the plot with colored symbols according to type: deletions (DEL, red), insertions (INS, green), duplications (DUP, yellow/orange), inversions (INV, purple), breakends (BND, pink), and copy number variations (CNV, teal). Connecting lines in the center link genomic regions involved in structural rearrangements, with red lines indicating inter-chromosomal connections and other colored lines representing various types of genomic alterations.

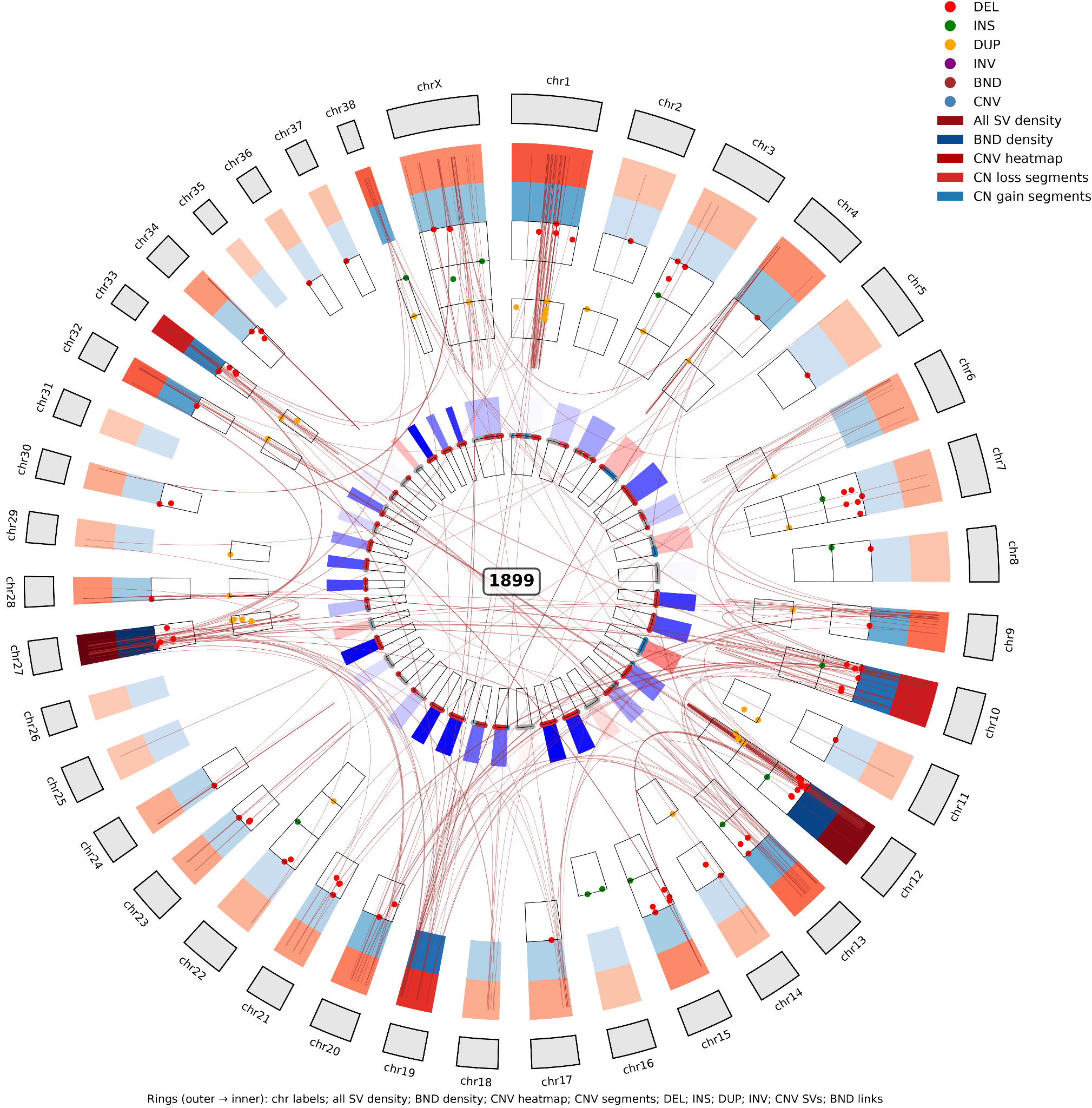

**Fig. S11.** Circos plot displaying the genomic landscape of structural variants and copy number alterations in sample GP1899. The circular plot represents the entire genome with chromosome labels around the periphery. From outer to inner, the concentric rings display: (1) chromosome ideograms with cytogenetic band positions, (2) all structural variant (SV) density shown as a dark red heatmap, (3) breakend (BND) density displayed in dark blue, (4) copy number variation (CNV) heatmap with gains in red/orange and losses in blue, (5) specific copy number gain segments (red bars), and (6) copy number loss segments (blue bars). Individual SVs are marked with colored symbols according to type: deletions (DEL, red), insertions (INS, green), duplications (DUP, yellow/orange), inversions (INV, purple), breakends (BND, pink), and copy number variations (CNV, teal). Connecting lines in the center link genomic regions involved in structural rearrangements, with red lines indicating inter-chromosomal connections and other colored lines representing various types of genomic alterations. This sample exhibits extensive genomic instability with numerous complex inter-chromosomal rearrangements, particularly involving chromosomes 2, 12, and others, along with widespread copy number alterations throughout the genome. The dense network of connecting lines indicates substantial chromosomal complexity and genomic rearrangements characteristic of chromothripsis or chromoplexy.

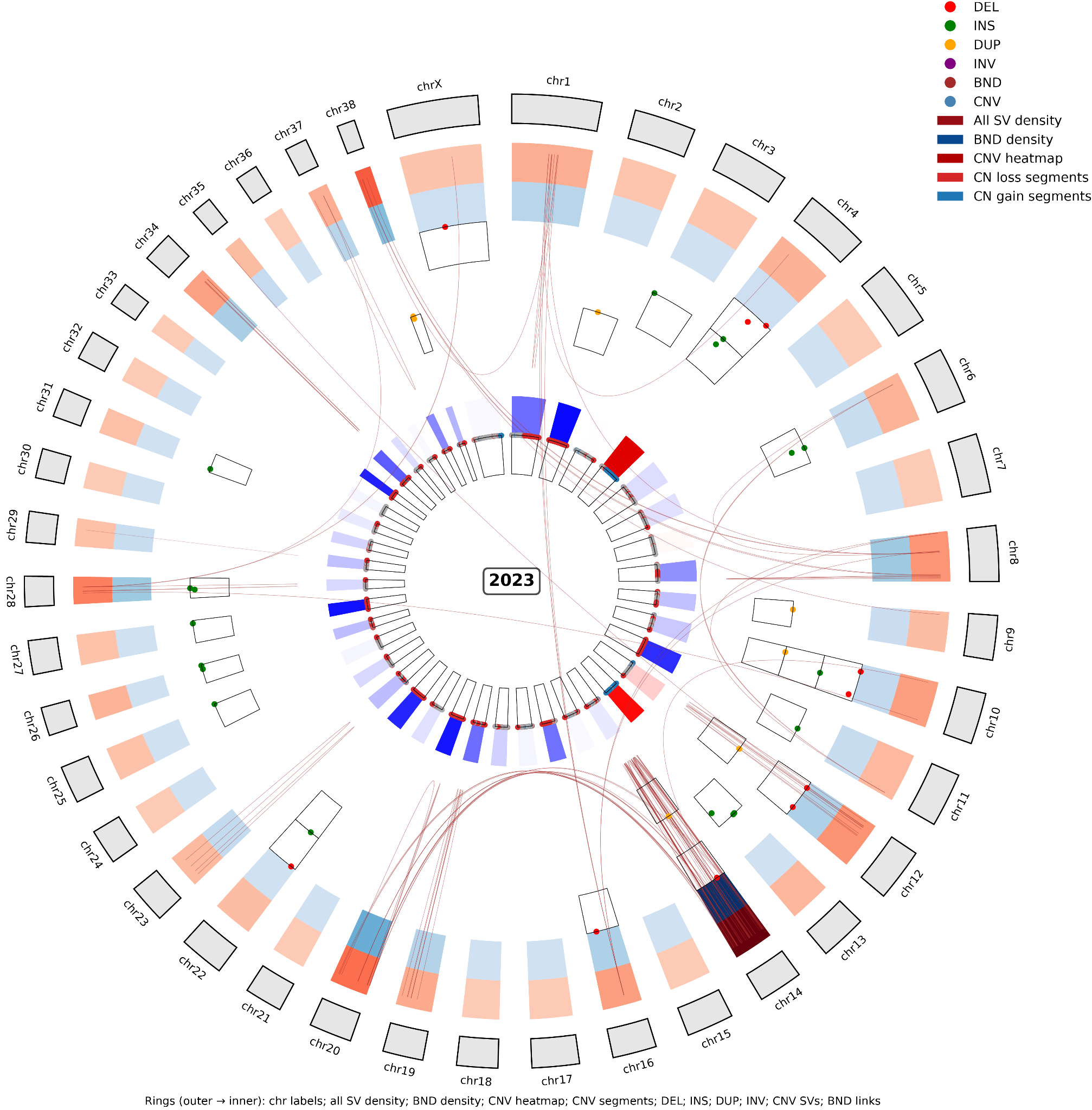

**Fig. S12.** Circos plot displaying the genomic landscape of structural variants and copy number alterations in sample GR2023. The circular plot represents the entire genome with chromosome labels around the periphery. From outer to inner, the concentric rings display: (1) chromosome ideograms with cytogenetic band positions, (2) all structural variant (SV) density shown as a dark red heatmap, (3) breakend (BND) density displayed in dark blue, (4) copy number variation (CNV) heatmap with gains in red/orange and losses in blue, (5) specific copy number gain segments (red bars), and (6) copy number loss segments (blue bars). Individual SVs are marked with colored symbols according to type: deletions (DEL, red), insertions (INS, green), duplications (DUP, yellow/orange), inversions (INV, purple), breakends (BND, pink), and copy number variations (CNV, teal). Connecting lines in the center link genomic regions involved in structural rearrangements, with red lines indicating inter-chromosomal connections and other colored lines representing various types of genomic alterations. This sample displays moderate genomic complexity with scattered SVs across multiple chromosomes, focal copy number alterations including prominent losses (blue segments) and gains (red segments), and several inter-chromosomal rearrangements, though less extensive than the highly complex chromothripsis pattern observed in GP1899.

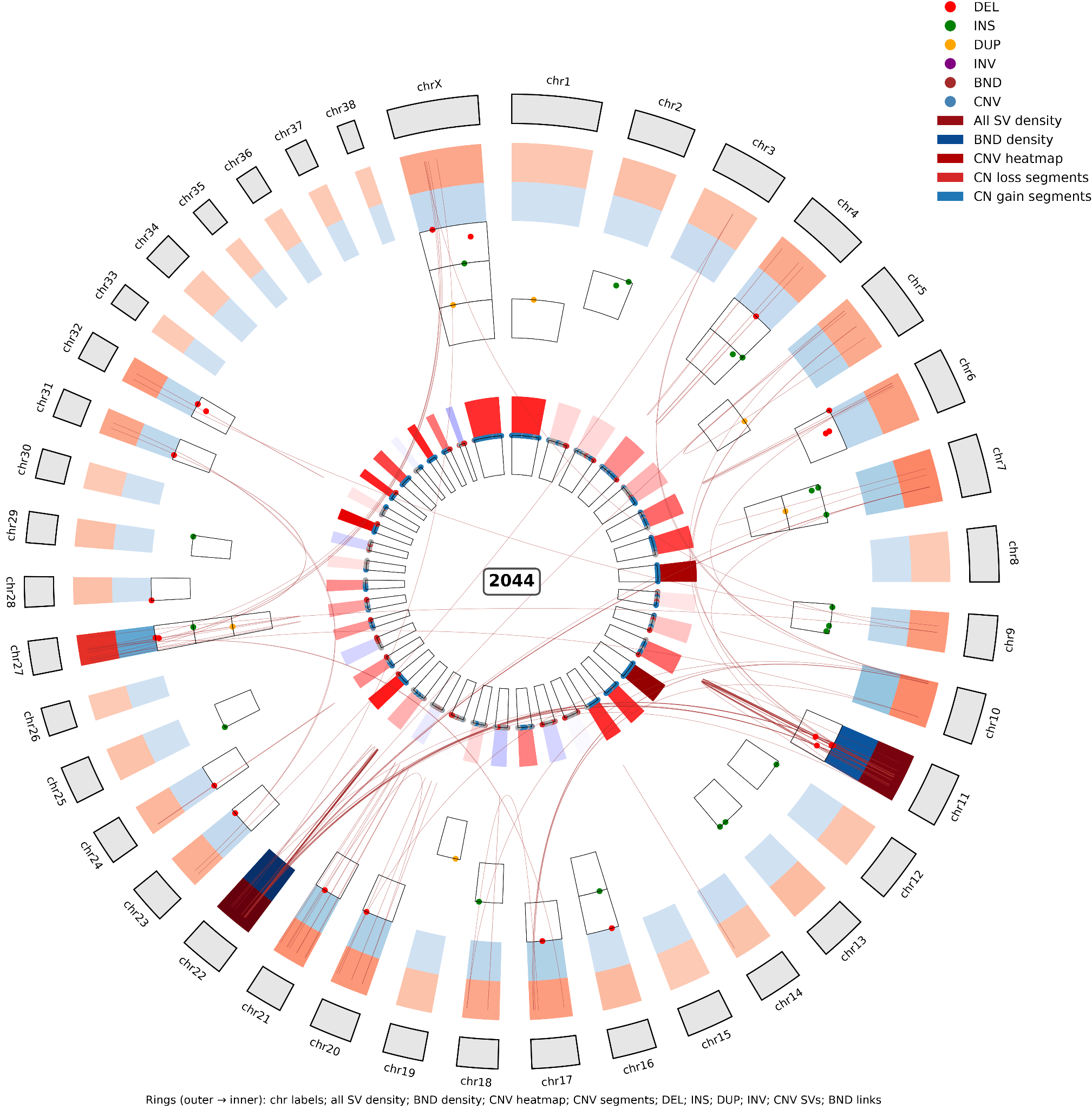

**Fig. S13.** Circos plot displaying the genomic landscape of structural variants and copy number alterations in sample GD2044. The circular plot represents the entire genome with chromosome labels around the periphery. From outer to inner, the concentric rings display: (1) chromosome ideograms with cytogenetic band positions, (2) all structural variant (SV) density shown as a dark red heatmap, (3) breakend (BND) density displayed in dark blue, (4) copy number variation (CNV) heatmap with gains in red/orange and losses in blue, (5) specific copy number gain segments (red bars), and (6) copy number loss segments (blue bars). Individual SVs are marked with colored symbols according to type: deletions (DEL, red), insertions (INS, green), duplications (DUP, yellow/orange), inversions (INV, purple), breakends (BND, pink), and copy number variations (CNV, teal). Connecting lines in the center link genomic regions involved in structural rearrangements, with red lines indicating inter-chromosomal connections and other colored lines representing various types of genomic alterations. This sample exhibits substantial genomic instability with numerous prominent copy number gains (bright red segments) distributed across multiple chromosomes, along with focal copy number losses (blue segments) and multiple inter-chromosomal rearrangements. The pattern reveals widespread chromosomal alterations characteristic of complex genomic rearrangements in osteosarcoma.

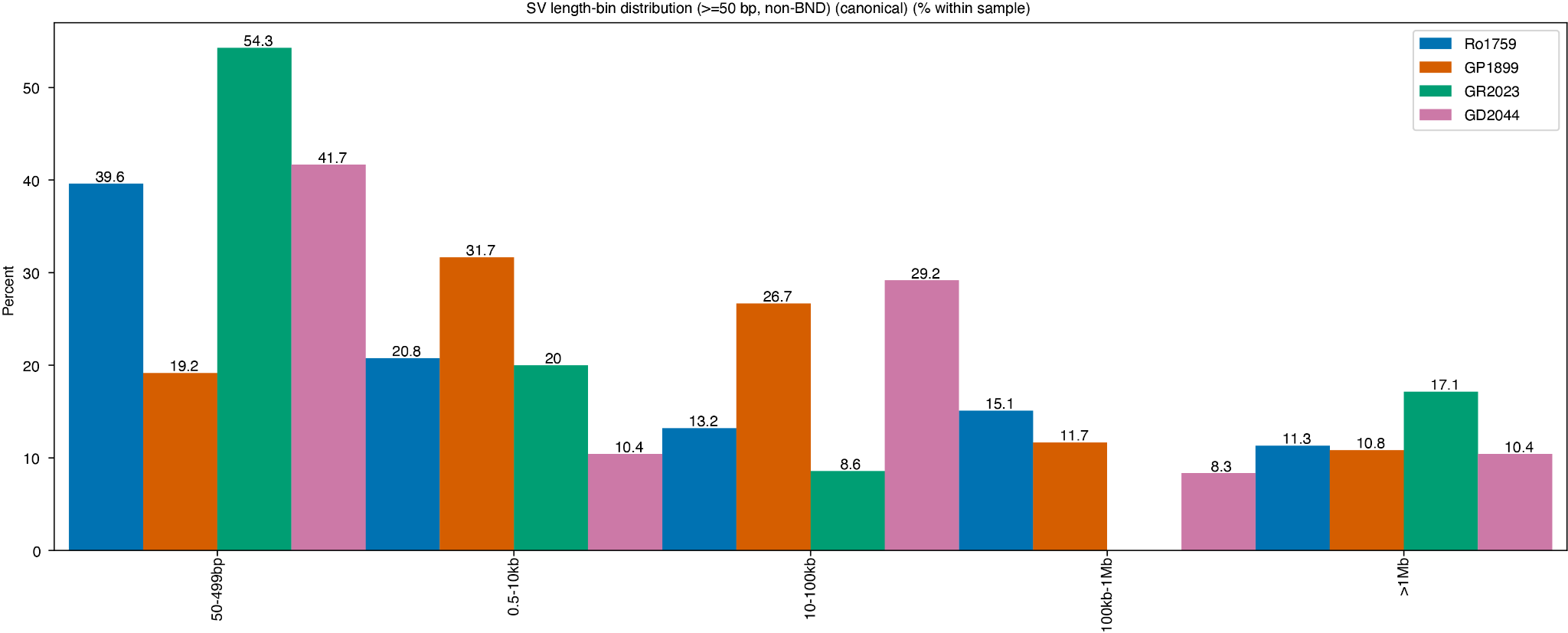

**Fig. S14.** Size distribution of structural variants across canine osteosarcoma tumors. Structural variants (SVs) ≥50 bp were binned by size and plotted as the percentage of total SVs within each sample. Only canonical SV types (deletions, duplications, insertions, inversions) are shown; breakends (BND) were excluded. Four tumor samples are displayed: Ro1759 (blue, n=226 SVs), GP1899 (orange, n=738 SVs), GR2023 (teal, n=184 SVs), and GD2044 (pink, n=238 SVs). Marked inter-tumor heterogeneity is evident in size profiles: GR2023 is dominated by small SVs (54.3% in 50-499 bp range), while GP1899 shows more even distribution across intermediate size classes. Large SVs (>1 Mb) represent a minor fraction across all samples (8.3-17.1%).

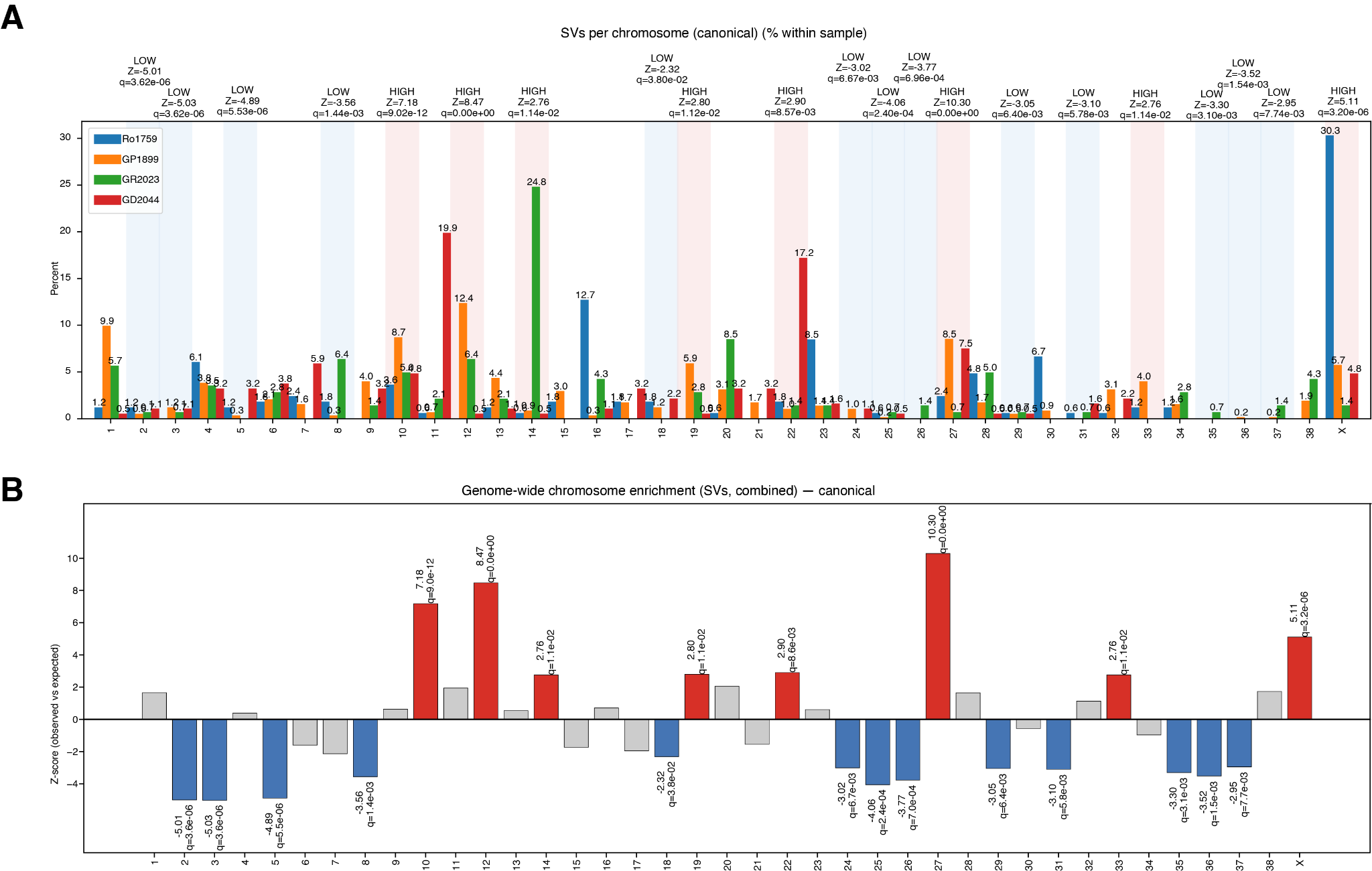

**Fig. S15.** Chromosomal distribution and enrichment of canonical structural variants. (A) Chromosome-specific frequencies of canonical SVs shown as percentage within each sample for Ro1759 (blue), GP1899 (orange), GR2023 (green), and GD2044 (red). Chromosomes are annotated with enrichment status (LOW/HIGH), Z-scores, and q-values above each cluster. Chromosome X shows the highest frequency in Ro1759 (30.3%), while chromosome 14 shows elevated frequencies in GR2023 (24.8%) and GD2044 (19.9%). Multiple chromosomes display significant enrichment or depletion across samples. (B) Genome-wide chromosome enrichment analysis for canonical SVs across all samples combined. Z-scores represent deviation from expected SV distribution. Chromosomes with significant enrichment (red bars, q<0.05) include chromosomes 11, 12, 14, 20, 22, 27, 33, and X, with chromosome 27 showing the strongest enrichment (Z-score=10.30, q=0e+00). Significantly depleted chromosomes (blue bars, q<0.05) include chromosomes 3, 5, 6, 9, 19, 25, 26, 29, 31, 35, 36, and 37. Non-significant chromosomes are shown in gray.

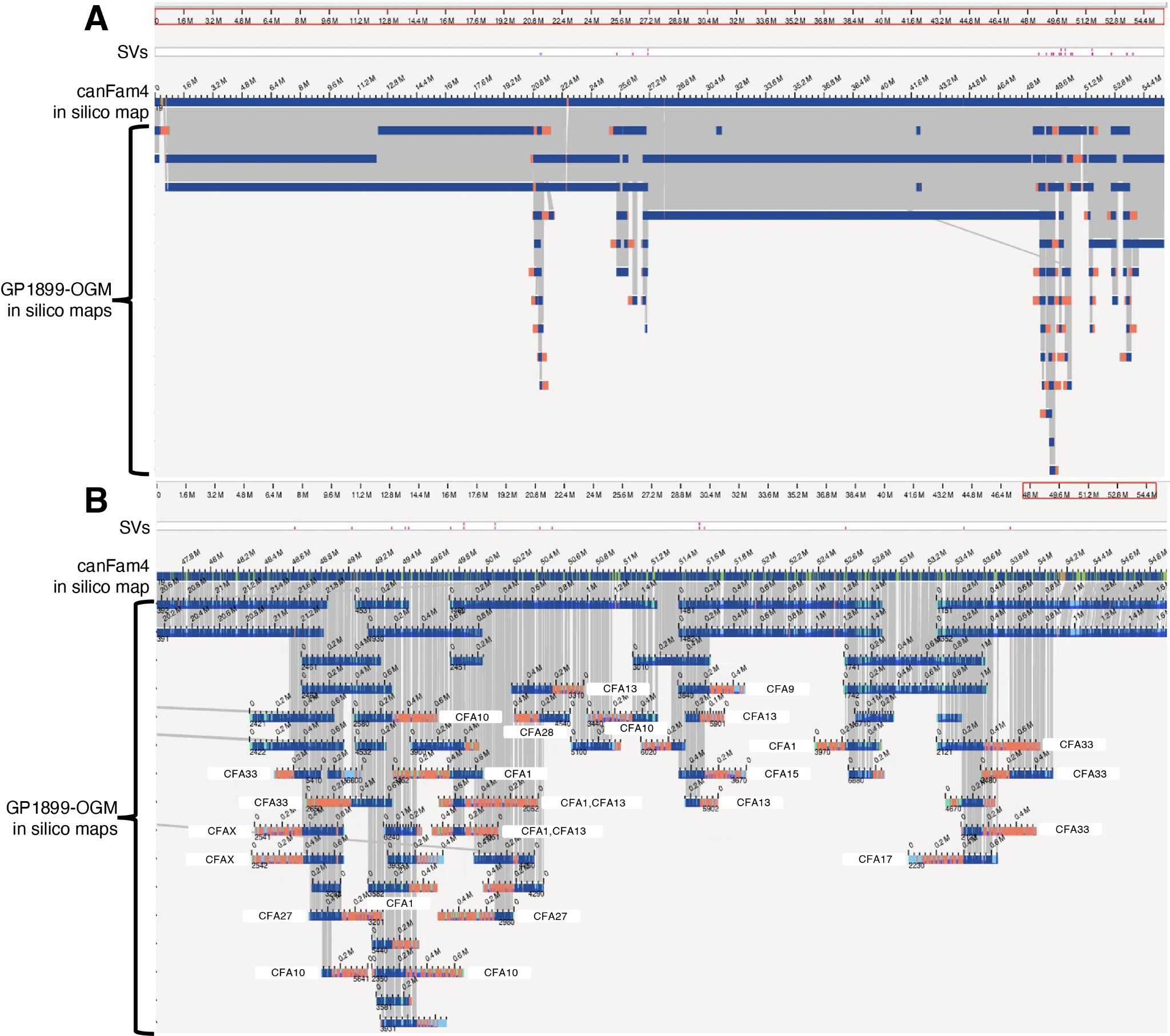

**Fig. S16.** Chromothripsis optical genome mapping (OGM) support. (A) Overview of structural variations (SVs) detected across a ~64 Mb genomic region. The top track shows the canFam4 reference genome in silico map with expected restriction enzyme site patterns (blue), while the bottom track displays the GP1899 OGM in silico maps showing discordant patterns (blue - alignment and red-misalignment marks) indicative of structural rearrangements. (B) Detailed view of a ~53 Mb subregion revealing complex chromosomal rearrangements. The GP1899-OGM data shows segments mapping to multiple different chromosomes (CFA1, CFA9, CFA10, CFA13, CFA15, CFA17, CFA27, CFA28, CFA33, and CFAX), suggesting inter-chromosomal rearrangements. Red marks indicate key breakpoint regions or areas of structural variation.

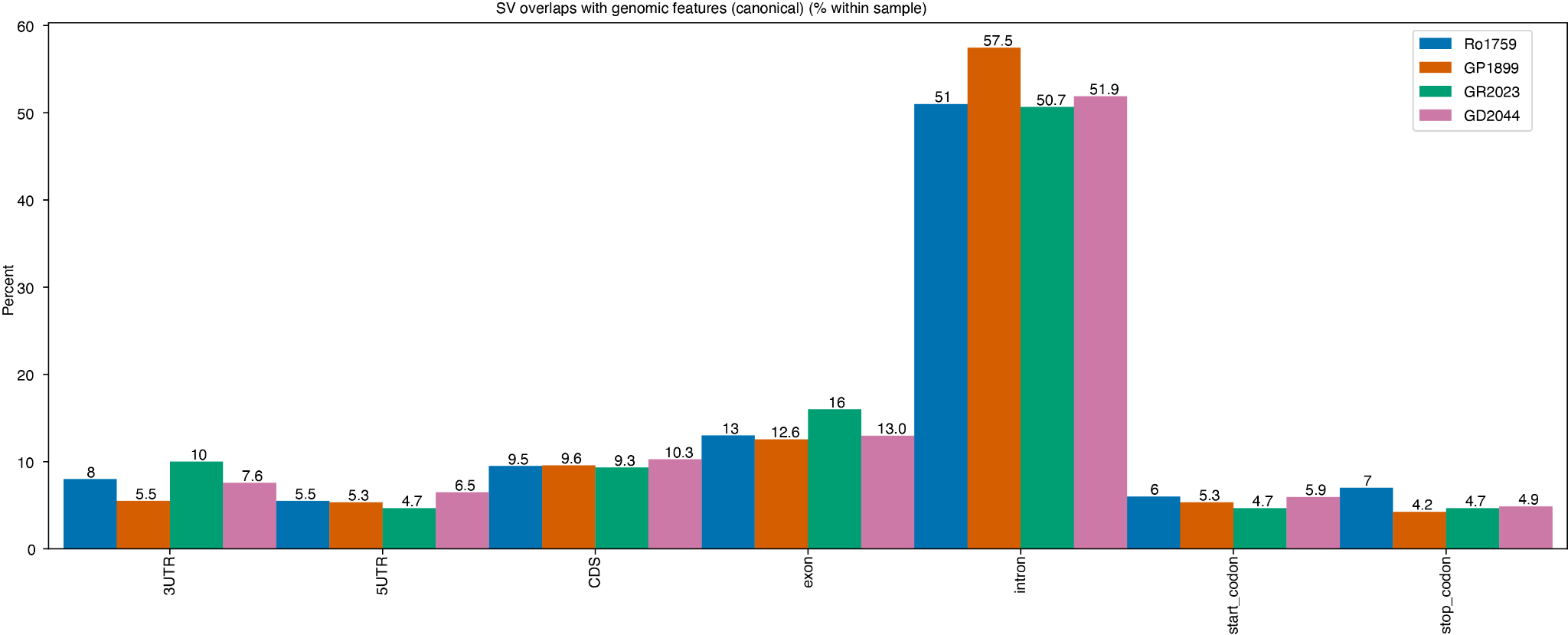

**Fig. S17.** Genomic feature overlap of structural variants in canine osteosarcoma tumors. Canonical structural variants were annotated for overlap with genomic features and plotted as the percentage of total SVs within each sample. Four tumor samples are shown: Ro1759 (blue, n=226 SVs), GP1899 (orange, n=738 SVs), GR2023 (teal, n=184 SVs), and GD2044 (pink, n=238 SVs). Intronic regions account for the majority of SV overlaps across all samples (50.7-57.5%), reflecting the prevalence of non-coding genomic space. Exonic overlaps range from 13.0-16.0%, with coding sequence (CDS) overlaps at 9.3-10.3%. UTR and codon-specific overlaps are relatively low and consistent across samples (3'UTR: 5.5-10.0%; 5'UTR: 4.7-6.5%; start/stop codons: 4.2-7.0%). The similar distribution patterns across tumors suggest that SV formation is not strongly biased by specific genomic features in canine osteosarcoma.

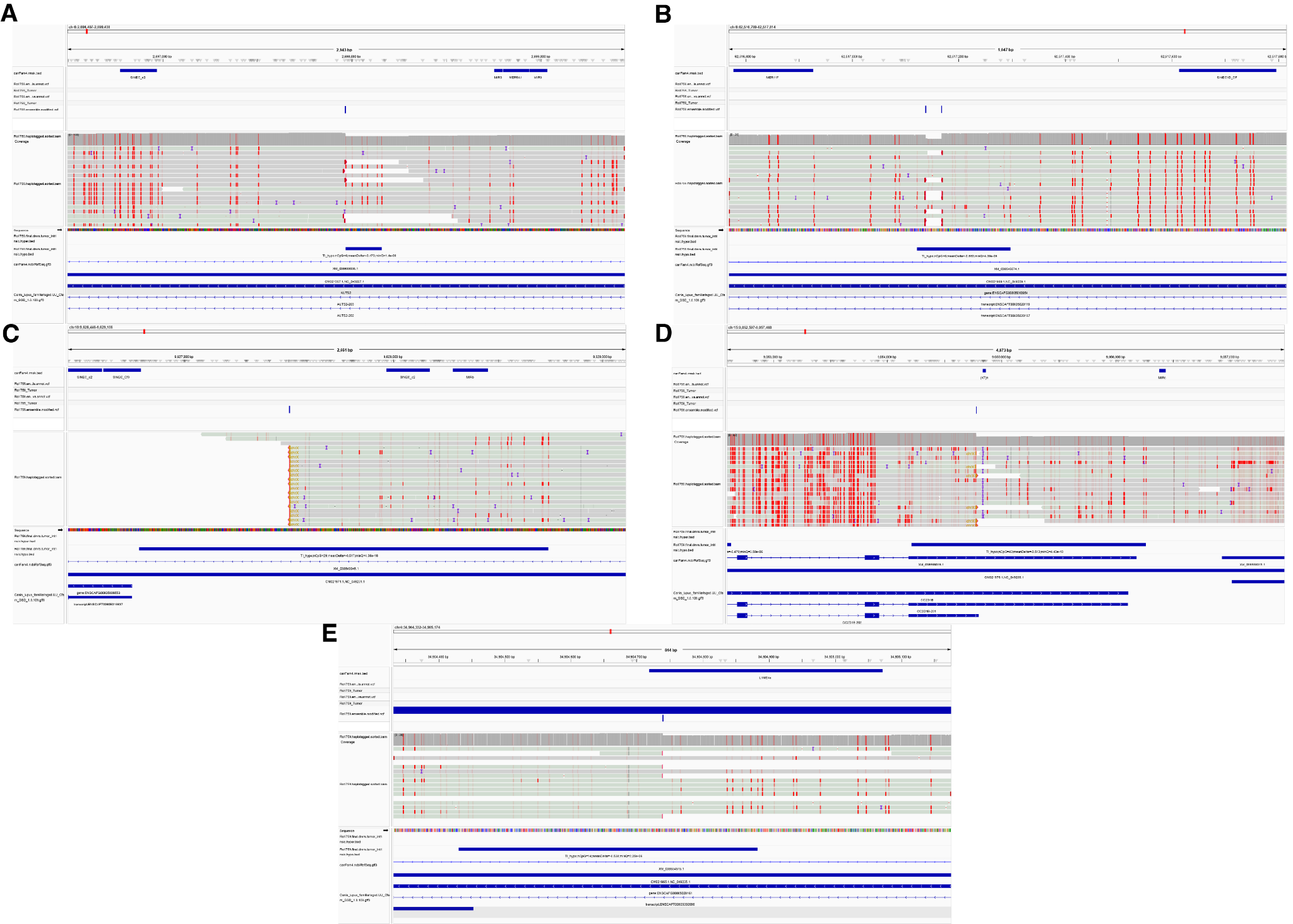

**Fig. S18.** Integrative Genome Browser (IGV) views showing sequence alignments and variant distributions across multiple genomic loci.

Panels (A-E) display genomic regions with aligned sequencing reads and identified variants. For each panel, the top track shows SV positions (dark blue bars), the middle section displays read alignments with variant positions marked as vertical colored lines (predominantly red), and the bottom tracks show additional gene annotations (dark blue bars). The gray shaded regions in the middle panels represent alignment coverage, with individual reads shown as horizontal gray bars. Genomic coordinates are indicated along the x-axis for each locus. Sample identifiers and track labels are shown on the left side of each panel.

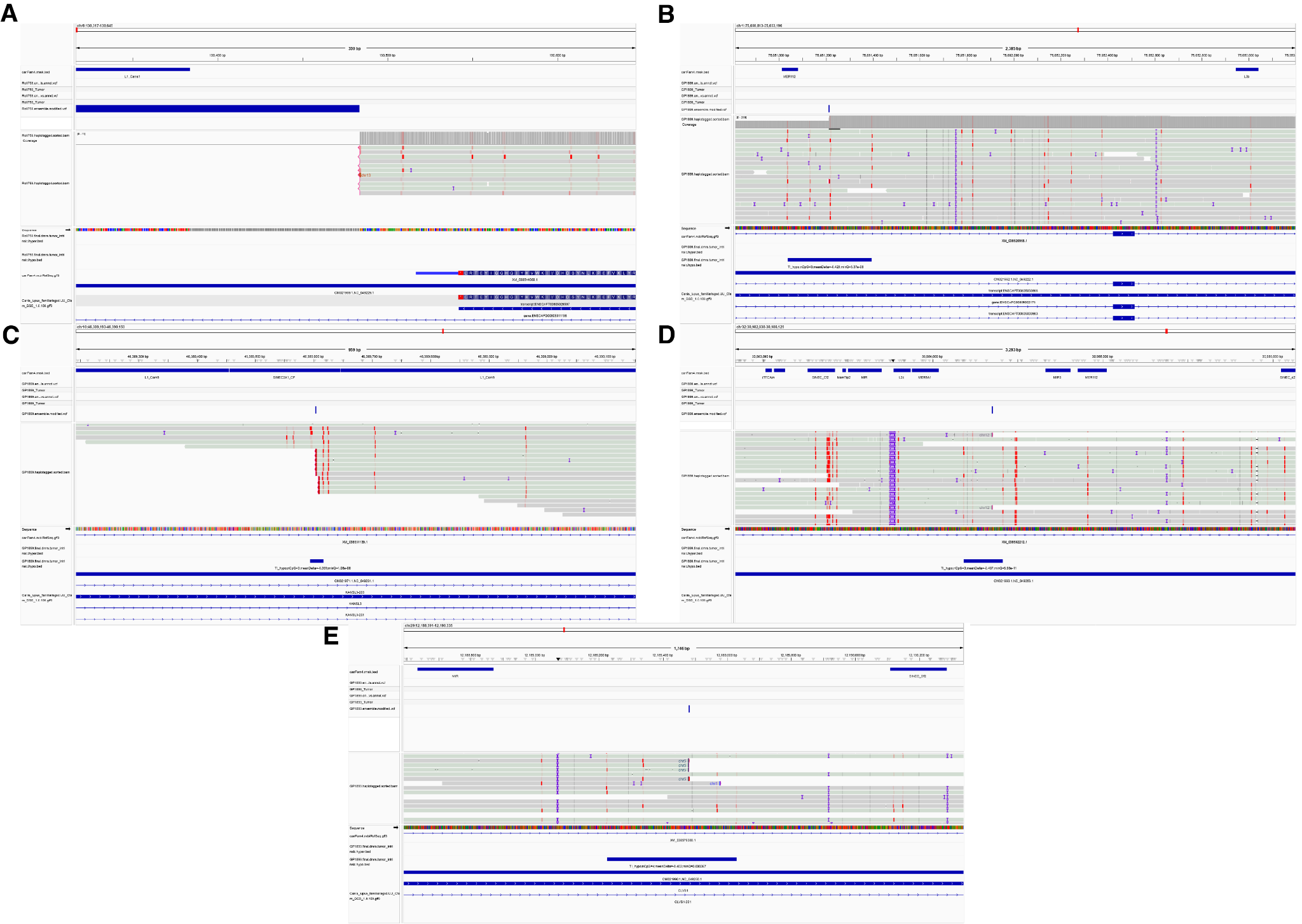

**Fig. S19.** Integrative Genome Browser (IGV) views showing sequence alignments and variant distributions across multiple genomic loci.

Panels (A-E) display genomic regions with aligned sequencing reads and identified variants. For each panel, the top track shows SV positions (dark blue bars), the middle section displays read alignments with variant positions marked as vertical colored lines (predominantly red), and the bottom tracks show additional gene annotations (dark blue bars). The gray shaded regions in the middle panels represent alignment coverage, with individual reads shown as horizontal gray bars. Genomic coordinates are indicated along the x-axis for each locus. Sample identifiers and track labels are shown on the left side of each panel.

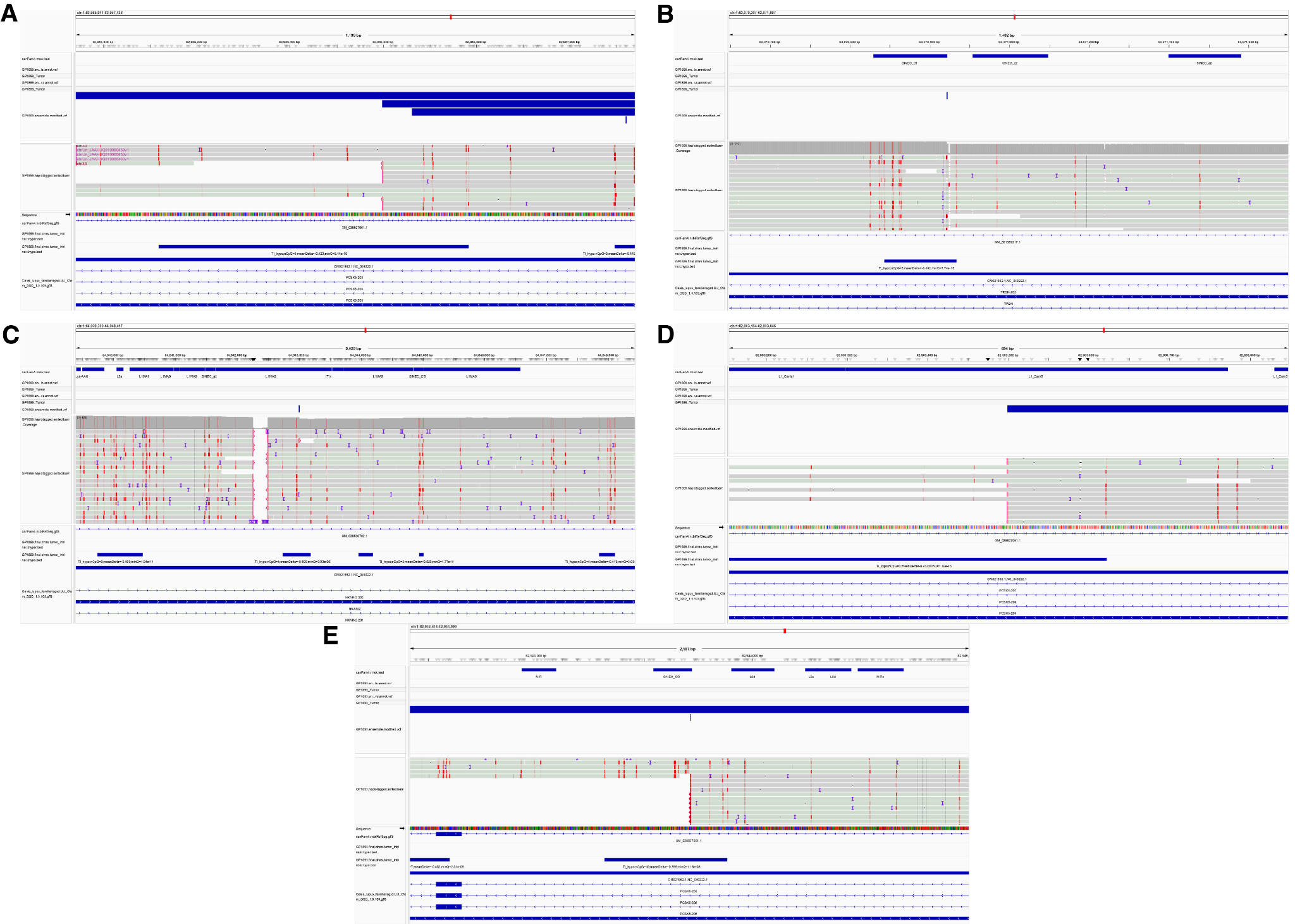

**Fig. S20.** Integrative Genome Browser (IGV) views showing sequence alignments and variant distributions across multiple genomic loci.

Panels (A-E) display genomic regions with aligned sequencing reads and identified variants. For each panel, the top track shows SV positions (dark blue bars), the middle section displays read alignments with variant positions marked as vertical colored lines (predominantly red), and the bottom tracks show additional gene annotations (dark blue bars). The gray shaded regions in the middle panels represent alignment coverage, with individual reads shown as horizontal gray bars. Genomic coordinates are indicated along the x-axis for each locus. Sample identifiers and track labels are shown on the left side of each panel.

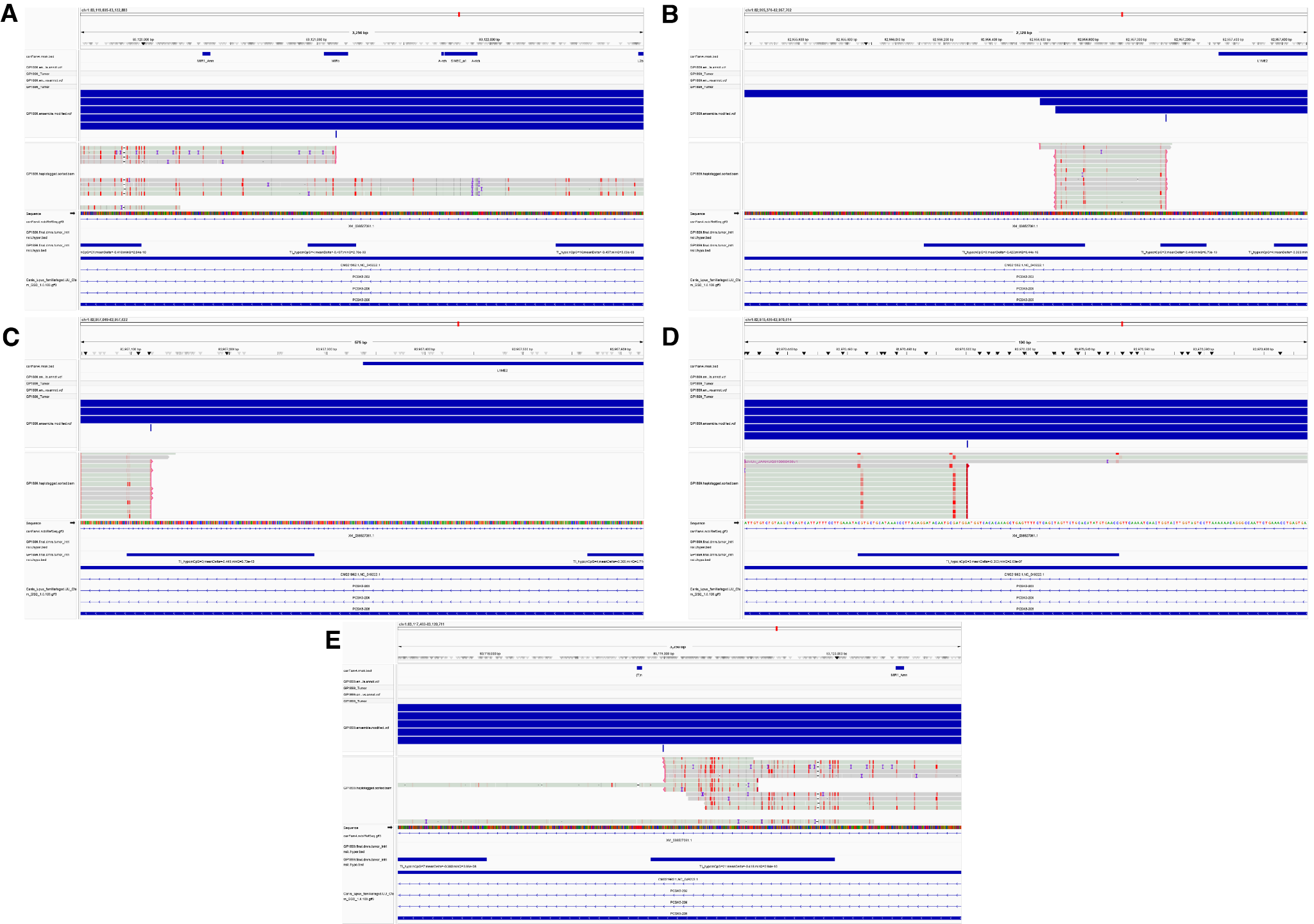

**Fig. S21.** Integrative Genome Browser (IGV) views showing sequence alignments and variant distributions across multiple genomic loci.

Panels (A-E) display genomic regions with aligned sequencing reads and identified variants. For each panel, the top track shows SV positions (dark blue bars), the middle section displays read alignments with variant positions marked as vertical colored lines (predominantly red), and the bottom tracks show additional gene annotations (dark blue bars). The gray shaded regions in the middle panels represent alignment coverage, with individual reads shown as horizontal gray bars. Genomic coordinates are indicated along the x-axis for each locus. Sample identifiers and track labels are shown on the left side of each panel.

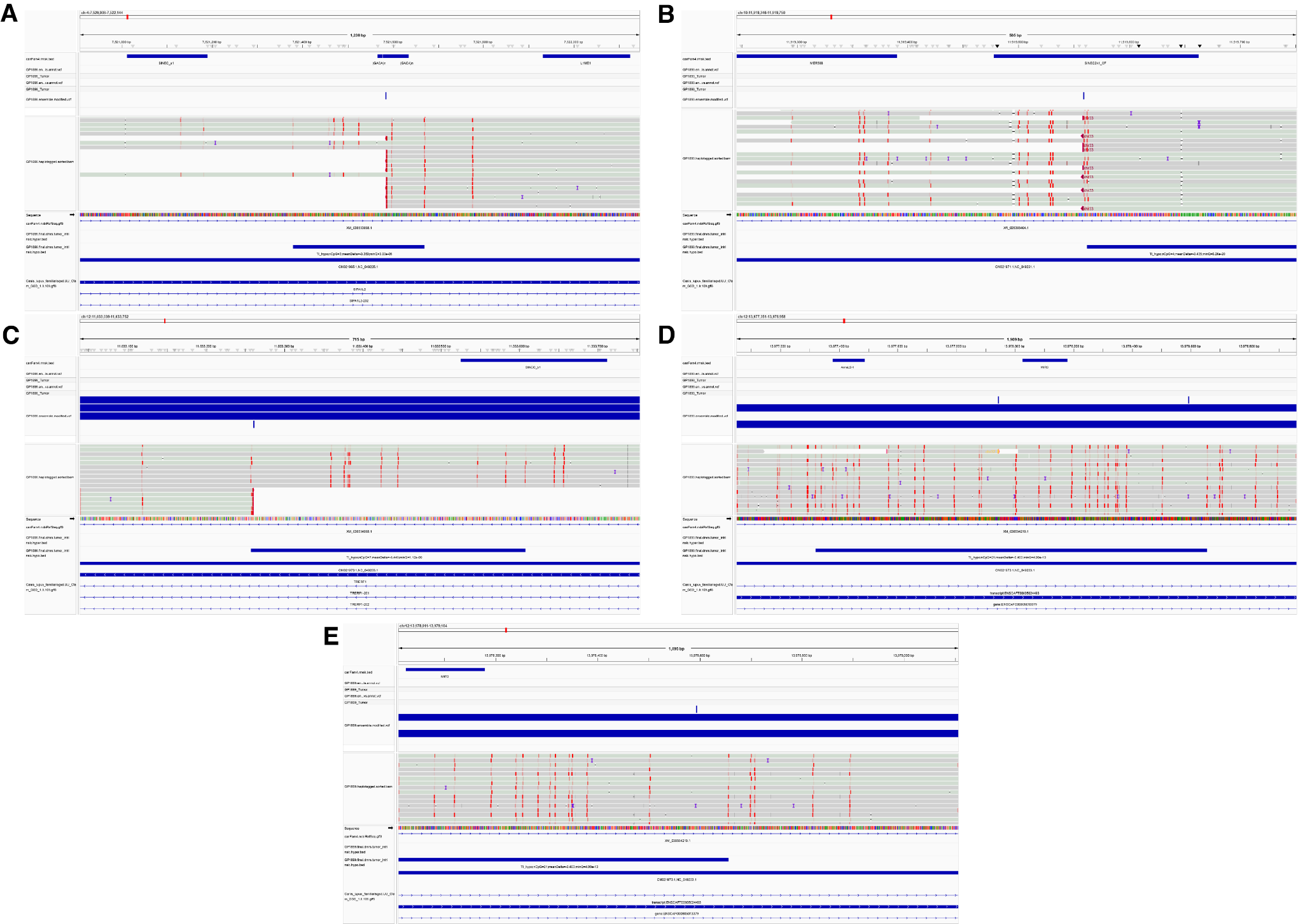

**Fig. S22.** Integrative Genome Browser (IGV) views showing sequence alignments and variant distributions across multiple genomic loci.

Panels (A-E) display genomic regions with aligned sequencing reads and identified variants. For each panel, the top track shows SV positions (dark blue bars), the middle section displays read alignments with variant positions marked as vertical colored lines (predominantly red), and the bottom tracks show additional gene annotations (dark blue bars). The gray shaded regions in the middle panels represent alignment coverage, with individual reads shown as horizontal gray bars. Genomic coordinates are indicated along the x-axis for each locus. Sample identifiers and track labels are shown on the left side of each panel.

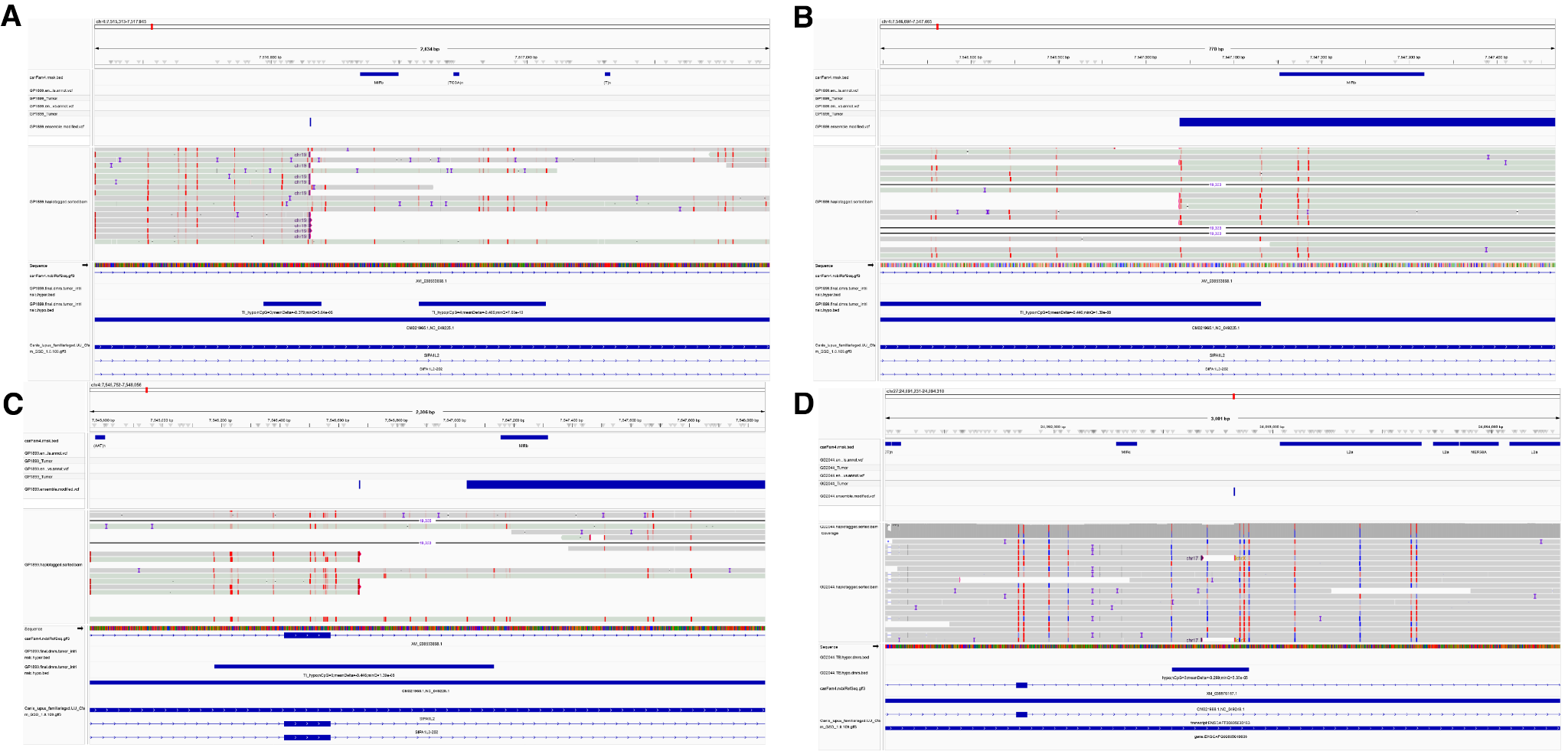

**Fig. S23.** Integrative Genome Browser (IGV) views showing sequence alignments and variant distributions across multiple genomic loci.

Panels (A-D) display genomic regions with aligned sequencing reads and identified variants. For each panel, the top track shows SV positions (dark blue bars), the middle section displays read alignments with variant positions marked as vertical colored lines (predominantly red), and the bottom tracks show additional gene annotations (dark blue bars). The gray shaded regions in the middle panels represent alignment coverage, with individual reads shown as horizontal gray bars. Genomic coordinates are indicated along the x-axis for each locus. Sample identifiers and track labels are shown on the left side of each panel.

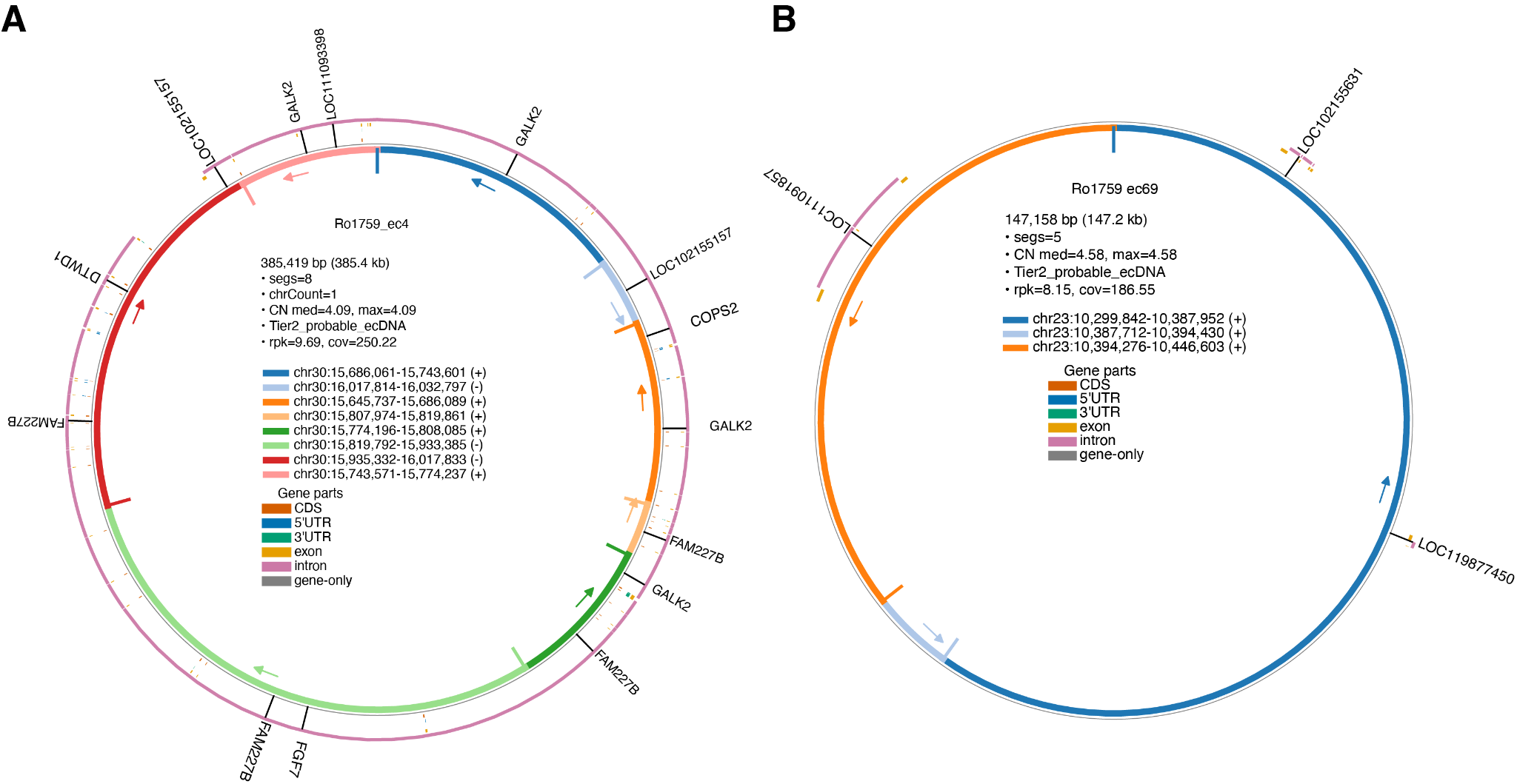

**Fig. S24.** Circular representations of extrachromosomal DNA (ecDNA) structures from canine osteosarcoma sample Ro1759. Circos plots illustrating the genomic architecture of two ecDNA elements: (A) Ro1759_ec4 (385.4 kb, 8 segments, median copy number=4.09, Tier2_probable_ecDNA) and (B) Ro1759_ec69 (147.2 kb, 5 segments, median copy number=4.58, Tier2_probable_ecDNA). Inner colored arcs represent genomic segments derived from specific chromosomal locations, with coordinates and strand orientation indicated by (+) or (-) symbols. Panel A shows eight segments exclusively from chromosome 30 spanning multiple regions, while Panel B displays five segments from chromosome 23. Segments are color-coded by gene feature annotation: CDS (orange), 5'UTR (blue), 3'UTR (teal), exon (yellow), intron (pink), and gene-only regions (gray). Arrows indicate structural orientation and connection points between segments. The outer pink ring shows the reference genome context with annotated genes labeled around the circle, including GALK2, FAM227B, RSF7 in panel A, and LOC102156531, LOC119677450 in panel B. Sequencing metrics include reads per kilobase (rpk) and coefficient of variation (cov) for each ecDNA element.

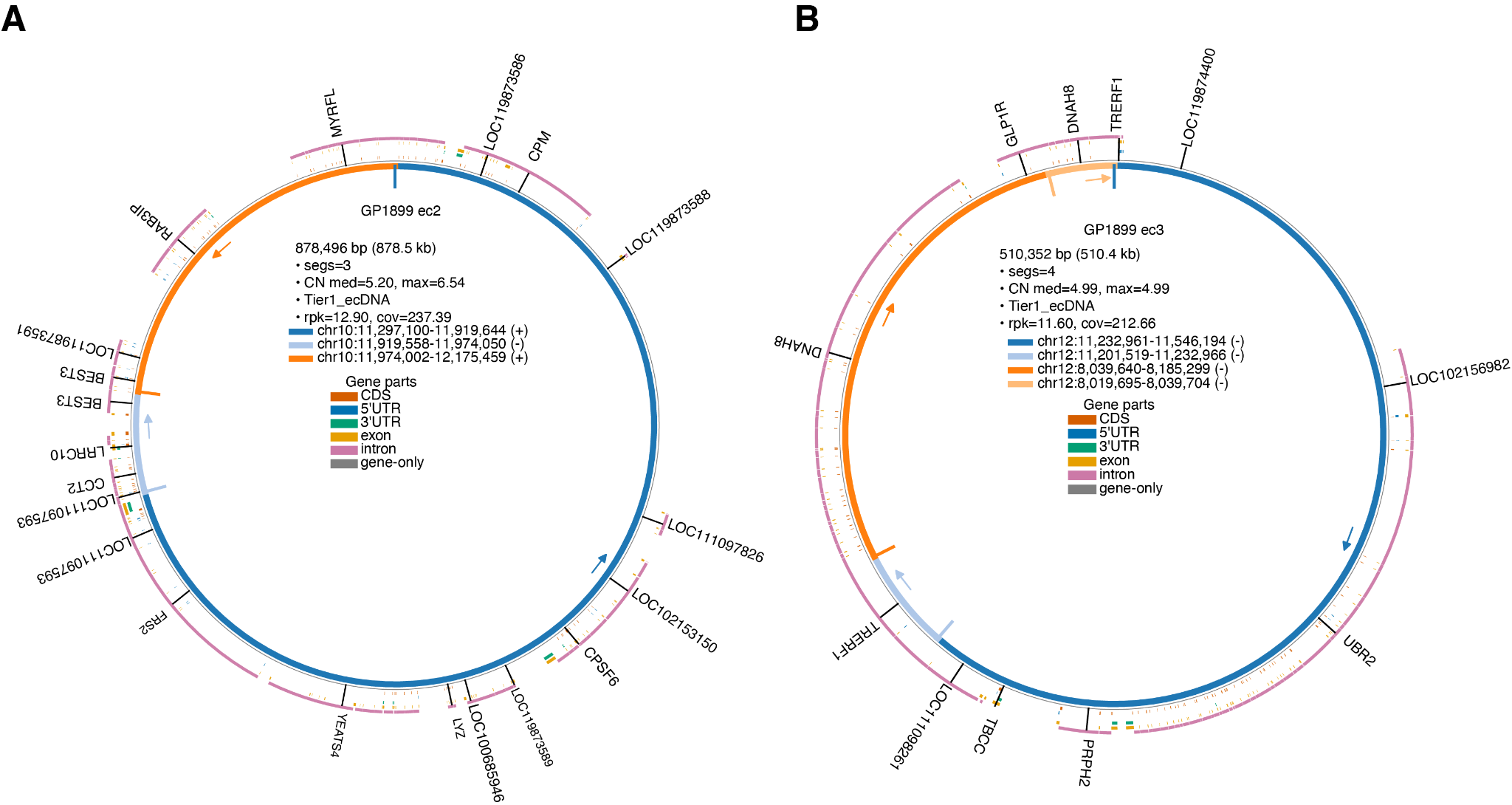

**Fig. S25.** Circular representations of extrachromosomal DNA (ecDNA) structures from canine osteosarcoma sample GP1899. Circos plots illustrating the genomic architecture of two Tier1 ecDNA elements: (A) GP1899_ec2 (878.5 kb, 3 segments, median copy number=5.20, Tier1_ecDNA) and (B) GP1899_ec3 (510.4 kb, 4 segments, median copy number=4.99, Tier1_ecDNA). Inner colored arcs represent genomic segments derived from specific chromosomal locations, with coordinates and strand orientation indicated by (+) or (-) symbols. Panel A shows three segments exclusively from chromosome 10, while Panel B displays four segments from chromosome 12. Segments are color-coded by gene feature annotation: CDS (orange), 5'UTR (blue), 3'UTR (teal), exon (yellow), intron (pink), and gene-only regions (gray). Arrows indicate structural orientation and connection points between segments. The outer pink ring shows the reference genome context with annotated genes labeled around the circles. Notable genes in panel A include NYFRL, Cryy, OBSCN, FANS4, and CCT2. Panel B includes GLRA, DNAMB, TRFRF1, and PRPHP2. Sequencing metrics include reads per kilobase (rpk) and coefficient of variation (cov) for each ecDNA element.

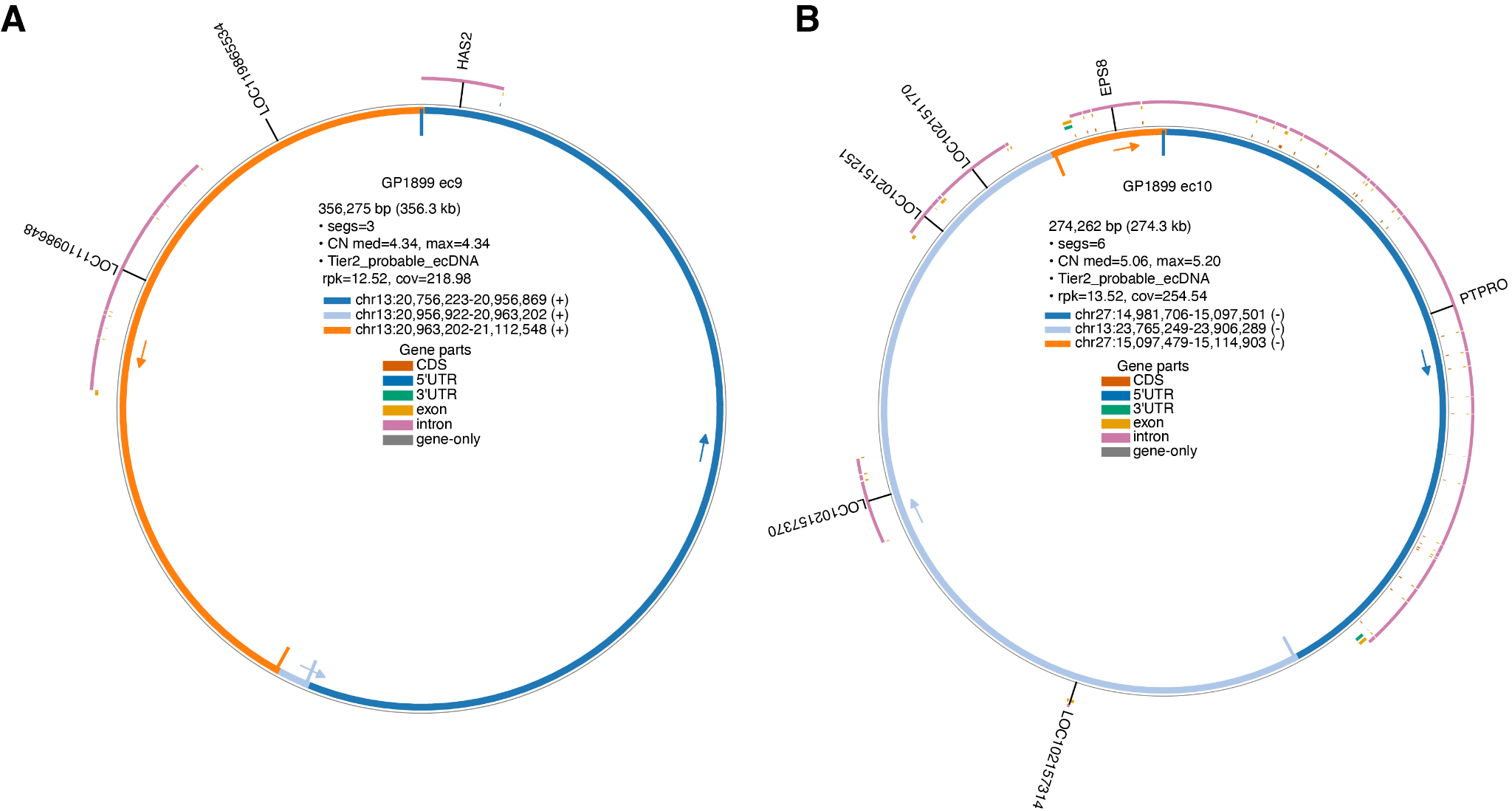

**Fig. S26.** Circular representations of extrachromosomal DNA (ecDNA) structures from canine osteosarcoma sample GP1899. Circos plots illustrating the genomic architecture of two Tier2 ecDNA elements: (A) GP1899_ec9 (356.3 kb, 3 segments, median copy number=4.34, Tier2_probable_ecDNA) and (B) GP1899_ec10 (274.3 kb, 6 segments, median copy number=5.06, Tier2_probable_ecDNA). Inner colored arcs represent genomic segments derived from specific chromosomal locations, with coordinates and strand orientation indicated by (+) or (-) symbols. Panel A shows three segments exclusively from chromosome 13, while Panel B displays six segments from chromosome 27. Segments are color-coded by gene feature annotation: CDS (orange), 5'UTR (blue), 3'UTR (teal), exon (yellow), intron (pink), and gene-only regions (gray). Arrows indicate structural orientation and connection points between segments. The outer pink ring shows the reference genome context with annotated genes labeled around the circles. Panel A includes HAS2, while Panel B features EPS8, PTPRO, and multiple LOC genes. Sequencing metrics include reads per kilobase (rpk) and coefficient of variation (cov) for each ecDNA element.

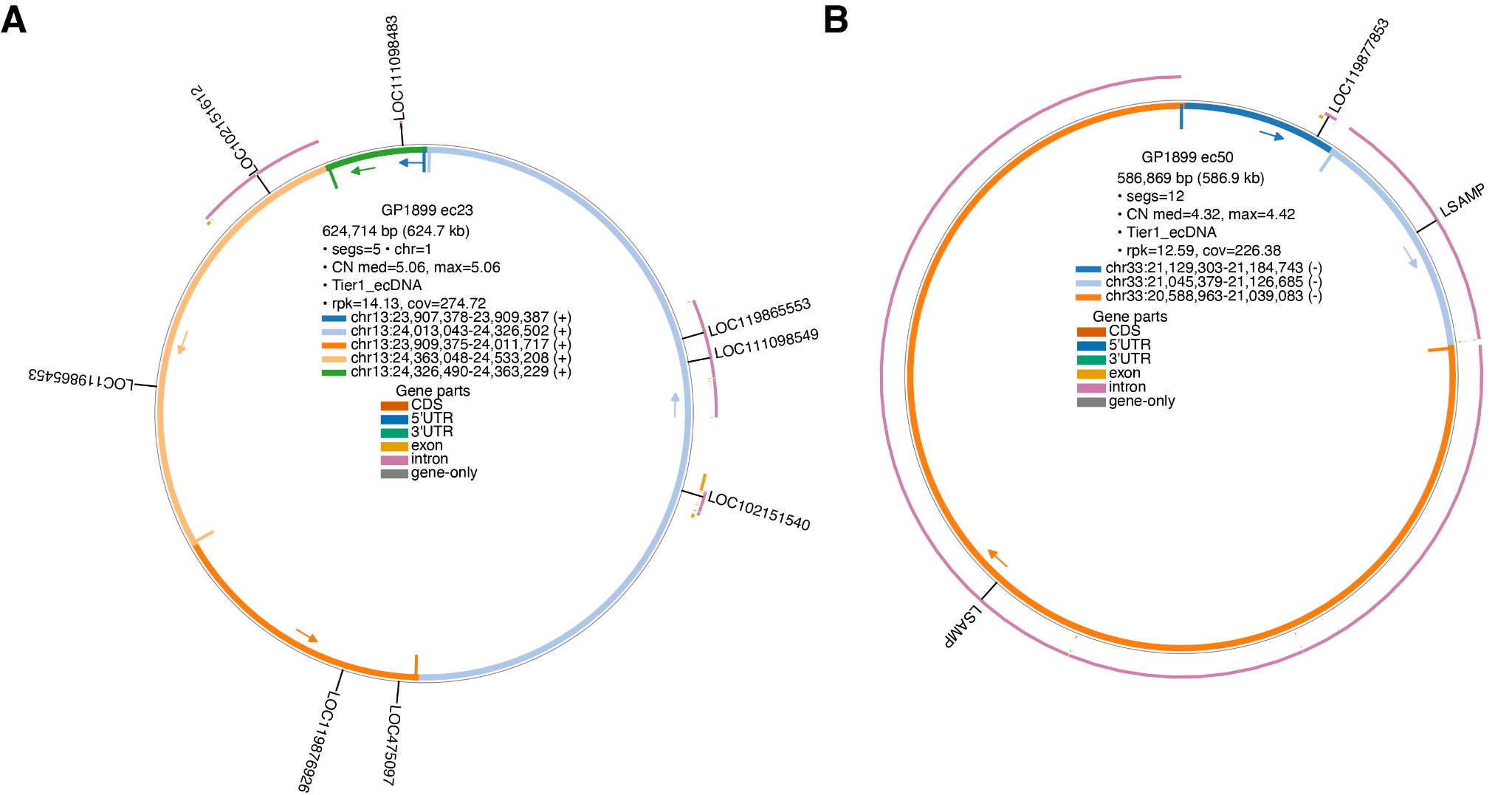

**Fig. S27.** Circular representations of extrachromosomal DNA (ecDNA) structures from canine osteosarcoma sample GP1899. Circos plots illustrating the genomic architecture of two Tier1 ecDNA elements: (A) GP1899_ec23 (624.7 kb, 5 segments, median copy number=5.06, Tier1_ecDNA) and (B) GP1899_ec50 (586.9 kb, 12 segments, median copy number=4.32, Tier1_ecDNA). Inner colored arcs represent genomic segments derived from specific chromosomal locations, with coordinates and strand orientation indicated by (+) or (-) symbols. Panel A shows five segments exclusively from chromosome 13, all on the positive strand, while Panel B displays three segments from chromosome 33, all on the negative strand. Segments are color-coded by gene feature annotation: CDS (orange), 5'UTR (blue), 3'UTR (teal), exon (yellow), intron (pink), and gene-only regions (gray). Arrows indicate structural orientation and connection points between segments. The outer pink ring shows the reference genome context with annotated genes labeled around the circles. Panel A includes multiple LOC genes (LOC111098549, LOC119865553, LOC102151540, LOC119861507, LOC111098626, LOC117609), while Panel B features LOC108671886, LSAMP, and dMYS1. Sequencing metrics include reads per kilobase (rpk) and coefficient of variation (cov) for each ecDNA element.

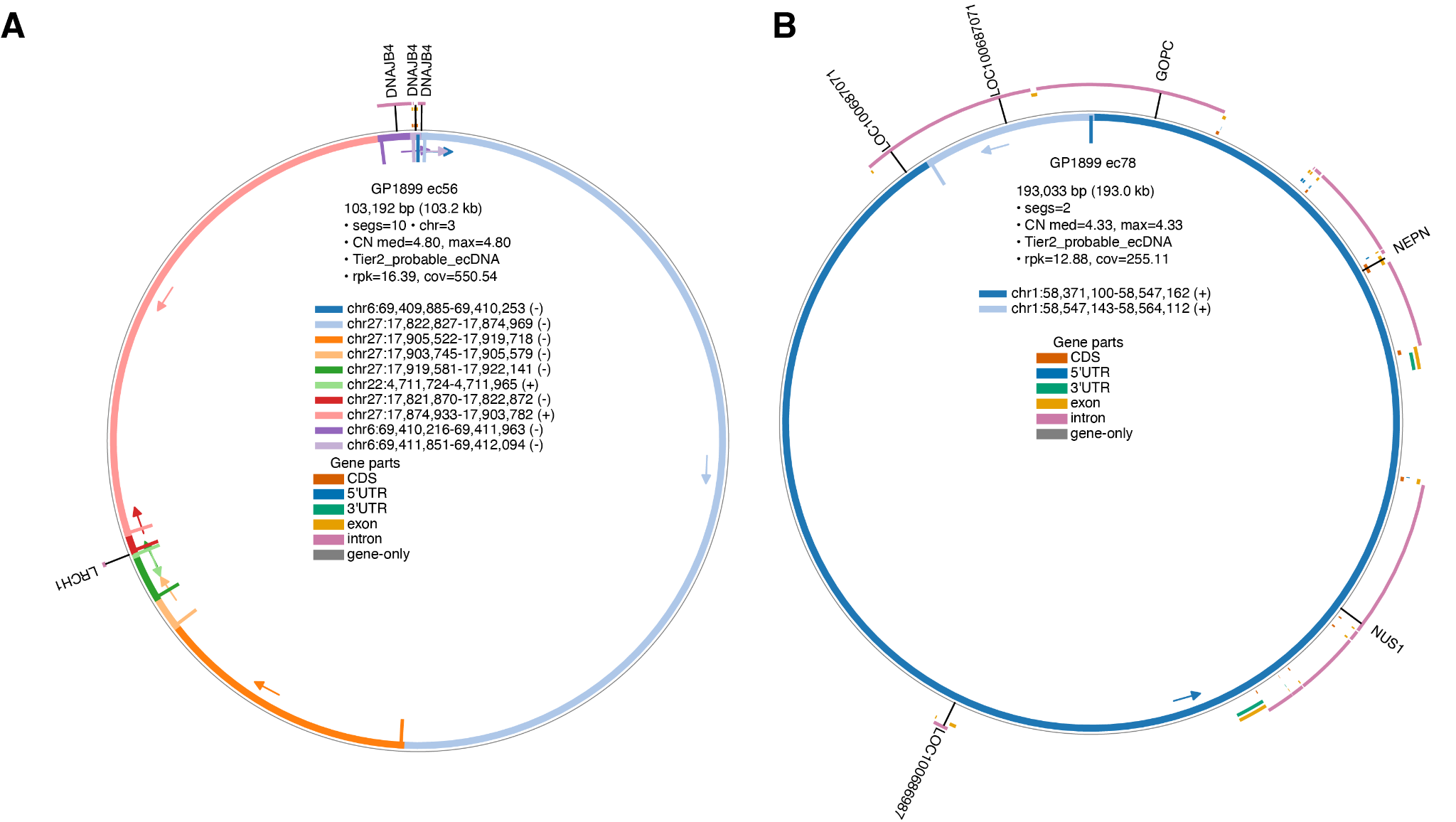

**Fig. S28.** Circular representations of extrachromosomal DNA (ecDNA) structures from canine osteosarcoma sample GP1899. Circos plots illustrating the genomic architecture of two Tier2 ecDNA elements: (A) GP1899_ec56 (103.2 kb, 10 segments, chromosome 3, median copy number=4.80, Tier2_probable_ecDNA) and (B) GP1899_ec78 (193.0 kb, 2 segments, median copy number=4.33, Tier2_probable_ecDNA). Inner colored arcs represent genomic segments derived from specific chromosomal locations, with coordinates and strand orientation indicated by (+) or (-) symbols. Panel A shows a complex structure with ten segments from multiple chromosomes (chr6, chr27, and chr22) on both positive and negative strands, while Panel B displays a simpler architecture with two segments exclusively from chromosome 1, both on the positive strand. Segments are color-coded by gene feature annotation: CDS (orange), 5'UTR (blue), 3'UTR (teal), exon (yellow), intron (pink), and gene-only regions (gray). Arrows indicate structural orientation and connection points between segments. The outer pink ring shows the reference genome context with annotated genes labeled around the circles. Panel A includes DNAJB4 (appearing three times at the top) and LHCH1, while Panel B features LOC608040, LOC100689011, GlyC, NEPN, and NUS1. Sequencing metrics include reads per kilobase (rpk) and coefficient of variation (cov) for each ecDNA element.

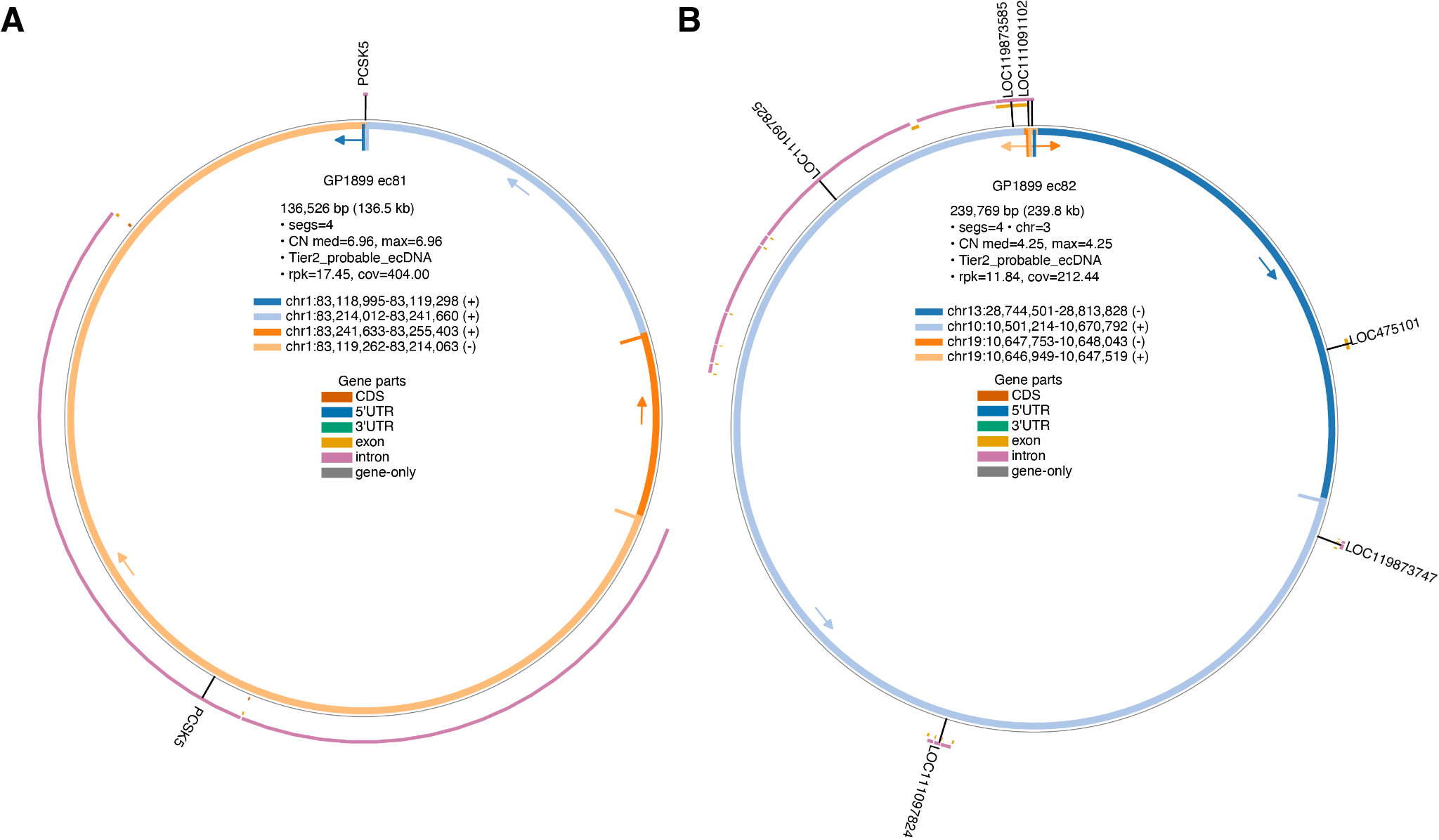

**Fig. S29.** Circular representations of extrachromosomal DNA (ecDNA) structures from canine osteosarcoma sample GP1899. Circos plots illustrating the genomic architecture of two Tier2 ecDNA elements: (A) GP1899_ec81 (136.5 kb, 4 segments, median copy number=6.96, Tier2_probable_ecDNA) and (B) GP1899_ec82 (239.8 kb, 4 segments, chromosome 3, median copy number=4.25, Tier2_probable_ecDNA). Inner colored arcs represent genomic segments derived from specific chromosomal locations, with coordinates and strand orientation indicated by (+) or (-) symbols. Panel A shows four segments exclusively from chromosome 1, with three on the positive strand and one on the negative strand, while Panel B displays four segments from multiple chromosomes (chr13, chr10, and chr19) with mixed strand orientations. Segments are color-coded by gene feature annotation: CDS (orange), 5'UTR (blue), 3'UTR (teal), exon (yellow), intron (pink), and gene-only regions (gray). Arrows indicate structural orientation and connection points between segments. The outer pink ring shows the reference genome context with annotated genes labeled around the circles. Panel A includes PCSK5 and FGBP6, while Panel B features multiple LOC genes (LOC119873585, LOC111091102, LOC110895, LOC475101, LOC119873747, LOC111097826). Sequencing metrics include reads per kilobase (rpk) and coefficient of variation (cov) for each ecDNA element.

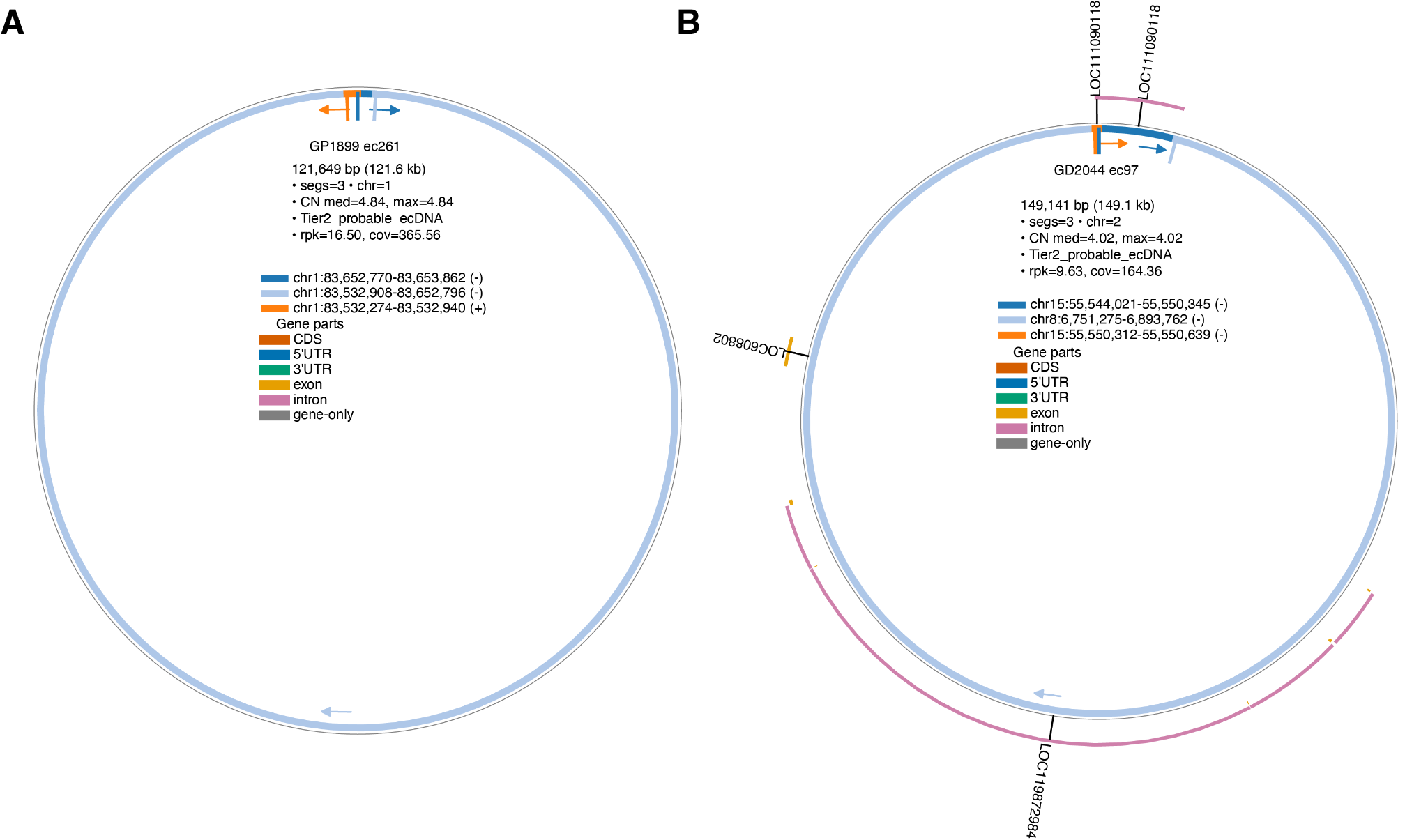

**Fig. S30.** Circular representations of extrachromosomal DNA (ecDNA) structures from canine osteosarcoma samples. Circos plots illustrating the genomic architecture of two Tier2 ecDNA elements: (A) GP1899_ec261 (121.6 kb, 3 segments, chromosome 1, median copy number=4.84, Tier2_probable_ecDNA) and (B) GD2044_ec97 (149.1 kb, 3 segments, chromosome 2, median copy number=4.02, Tier2_probable_ecDNA). Inner colored arcs represent genomic segments derived from specific chromosomal locations, with coordinates and strand orientation indicated by (+) or (-) symbols. Panel A shows a simple structure with three segments exclusively from chromosome 1, with two on the negative strand and one on the positive strand, while Panel B displays three segments from two different chromosomes (chr15 and chr8), all on the negative strand. Segments are color-coded by gene feature annotation: CDS (orange), 5'UTR (blue), 3'UTR (teal), exon (yellow), intron (pink), and gene-only regions (gray). Arrows indicate structural orientation and connection points between segments. The outer pink ring shows the reference genome context with annotated genes labeled around the circles. Panel B includes LOC111090118 (appearing twice at the top), LOC608082, and LOC119882486. Sequencing metrics include reads per kilobase (rpk) and coefficient of variation (cov) for each ecDNA element.

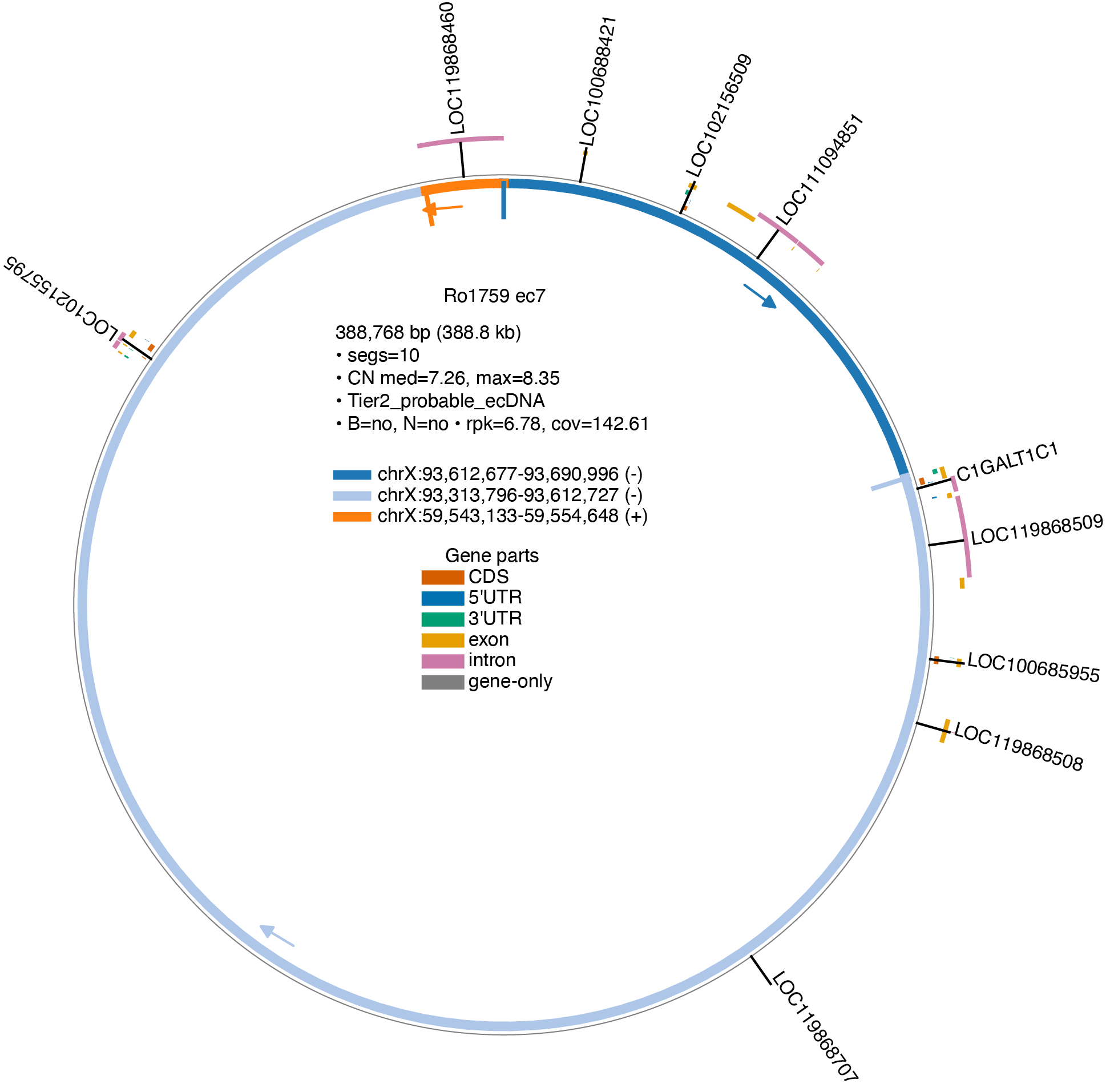

**Fig. S31.** Circular representation of an extrachromosomal DNA (ecDNA) structure from canine osteosarcoma sample Ro1759. Circos plot illustrating the genomic architecture of Ro1759_ec7 (388.8 kb, 10 segments, median copy number=7.26, Tier2_probable_ecDNA). This ecDNA was identified in tumor tissue only, with matched blood and normal tissue unavailable (B=no, N=no). Inner colored arcs represent genomic segments derived from specific chromosomal locations on chromosome X, with coordinates and strand orientation indicated by (+) or (-) symbols. The structure comprises three distinct genomic regions: two on the negative strand (shown in dark and light blue) and one on the positive strand (shown in orange). Segments are color-coded by gene feature annotation: CDS (orange), 5'UTR (blue), 3'UTR (teal), exon (yellow), intron (pink), and gene-only regions (gray). Arrows indicate structural orientation and connection points between segments. The outer pink ring shows the reference genome context with annotated genes labeled around the circle, including LOC119868460, LOC100998421, LOC102156309, LOC111094851, C1GALT1C1, LOC119868509, LOC100685955, LOC119868508, LOC119868707, and LOC115591755. Sequencing metrics include reads per kilobase (rpk=6.78) and coefficient of variation (cov=142.61).

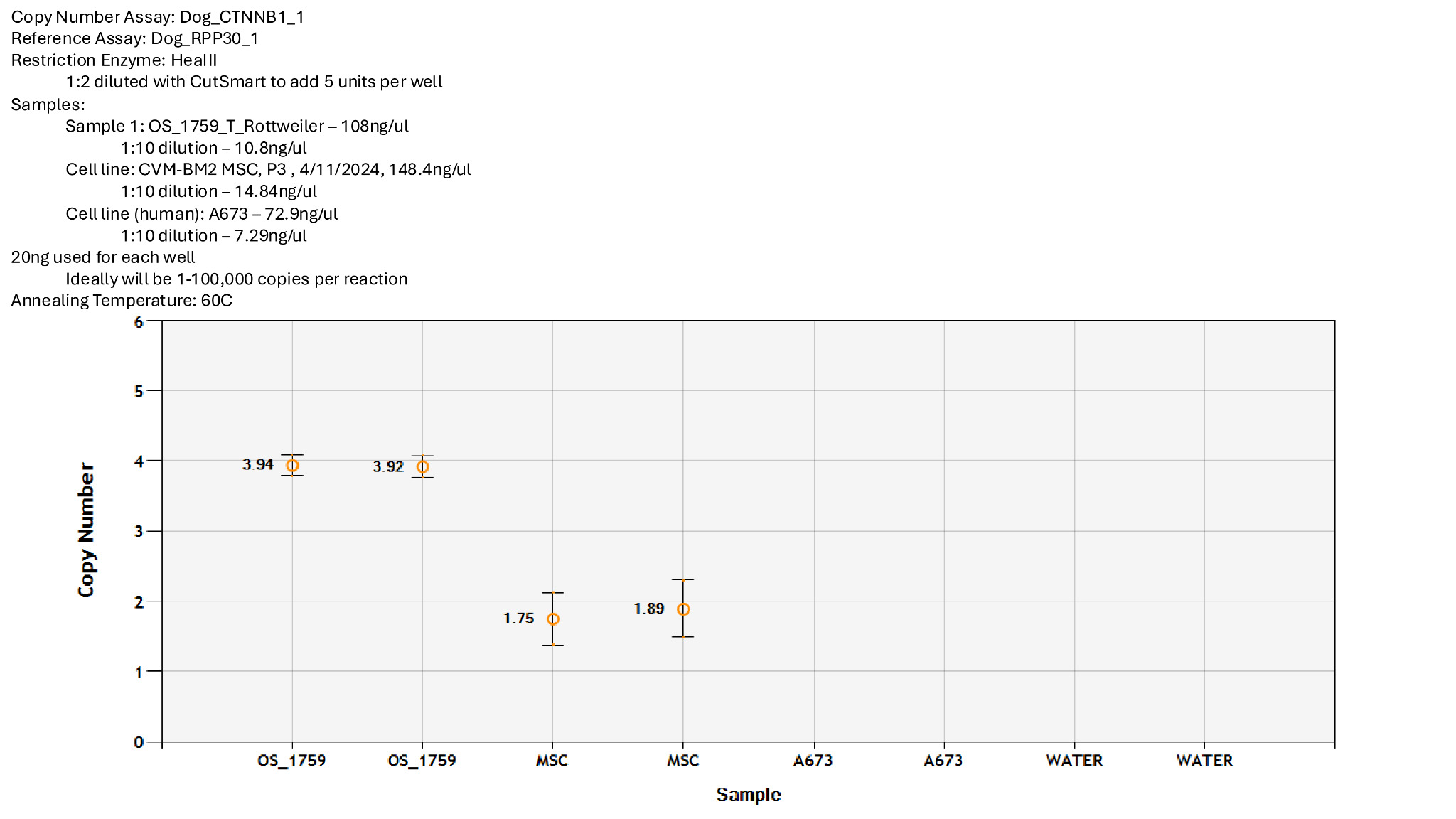

**Fig. S32.** Digital droplet PCR validation of *CTNNB1* copy number amplification in sample Ro1759. Digital droplet PCR (ddPCR) analysis confirming *CTNNB1* copy number gain in canine osteosarcoma sample Ro1759 (Rottweiler tumor). Copy number was determined using the Dog_CTNNB1_1 assay normalized to the reference gene *RPP30* (Dog_RPP30_1) following HealII restriction digestion. Ro1759 exhibits approximately 4-fold amplification of *CTNNB1* (mean copy number ~3.93), consistent with the ecDNA-mediated amplification detected by long-read sequencing. Control samples include CVM-BM2 canine mesenchymal stem cells (MSC, passage 3) and human A673 cells, both showing baseline diploid copy numbers (~1.75-1.89). Water served as a no-template negative control. Error bars represent standard deviation of technical replicates. Each well contained 20 ng of genomic DNA with an optimal copy number range of 1-100,000 copies per reaction. The temperature was 60°C.

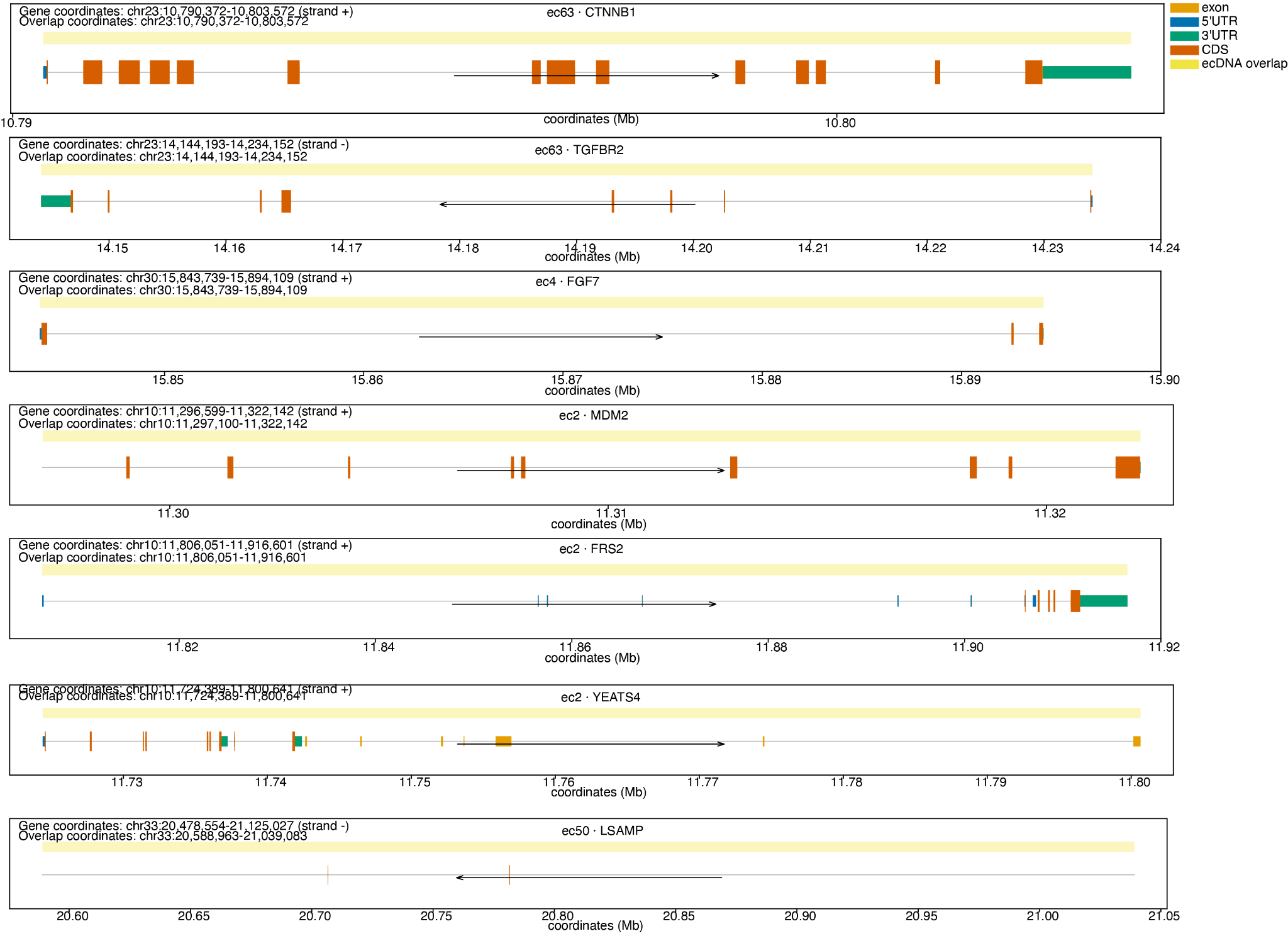

**Fig. S33.** Genomic structure and extrachromosomal DNA (ecDNA) overlap regions for cancer-associated genes. Each panel displays the genomic organization of genes amplified on distinct ecDNA elements (ec63: CTNNB1; ec4: FGF7; ec2: MDM2, FRS2, and YEATS4; ec50: LSAMP; ec63: TGFBR2). Gene coordinates and ecDNA overlap coordinates (in megabases, Mb) are indicated at the top of each panel. Exons are shown as orange boxes, with 5'UTRs in light orange, coding sequences (CDS) in dark teal, and 3'UTRs in green. Yellow shading highlights the regions of overlap between the ecDNA amplicons and the respective genes. Arrows indicate the direction of transcription (strand orientation: + or -). The genomic positions are based on human reference genome coordinates on the specified chromosomes (chr23, chr3, chr30, chr10, chr33).

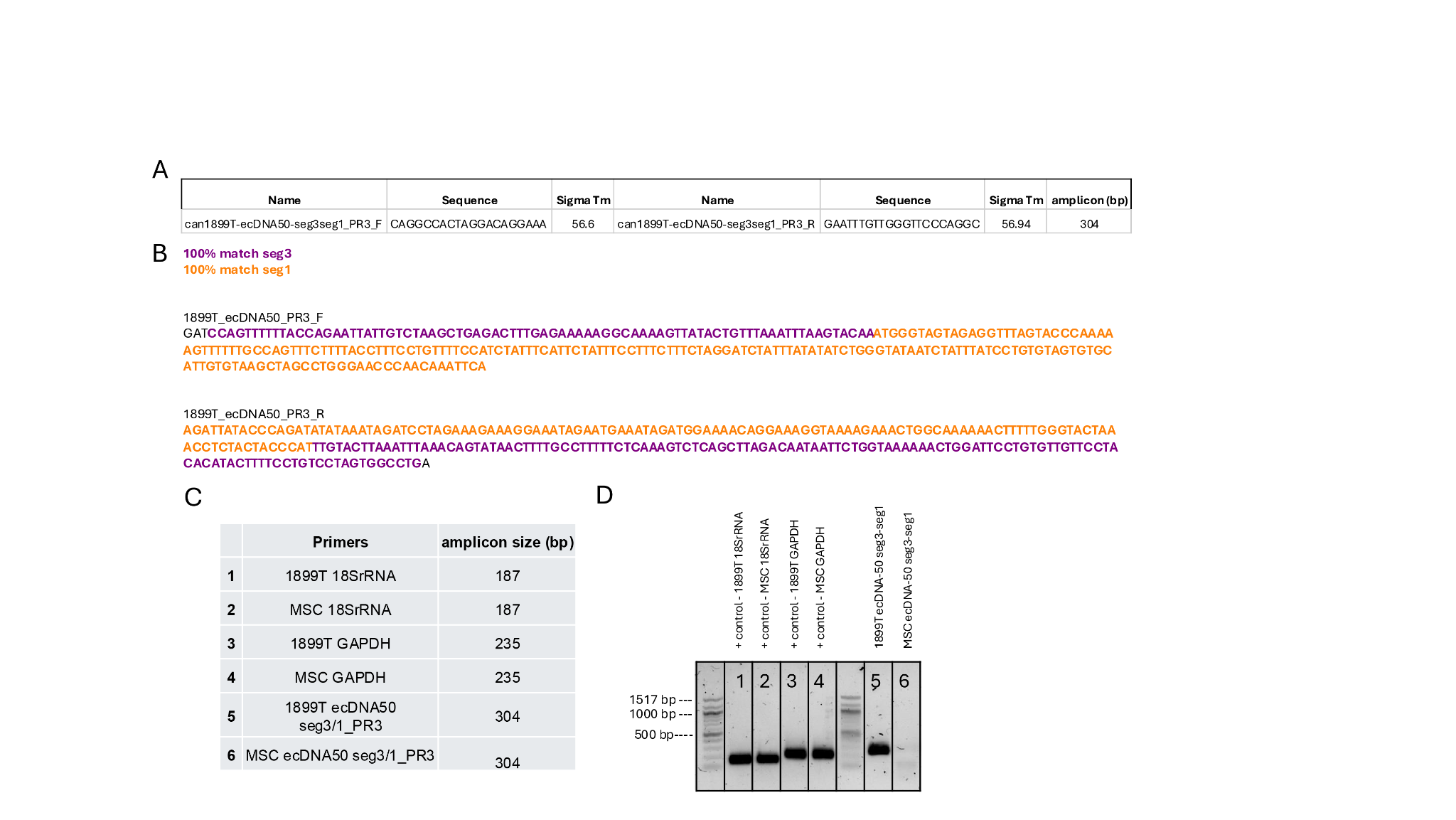

**Fig. S34.** PCR validation of the seg3-seg1 circular junction in ecDNA ec50 from sample GP1899. (A) Primer design for amplification across the seg3-seg1 junction of ec50. Forward and reverse primers (can1899T-ecDNA50-seg3seg1_PR3_F/R) were designed to span the predicted junction, yielding an expected amplicon of 304 bp. (B) Sequence alignment showing 100% match to seg3 (orange) and seg1 (red) at the circular junction breakpoint. (C) PCR primer pairs and expected amplicon sizes for validation and control genes. Primers 1-4 target endogenous reference genes (18SrRNA and GAPDH) in tumor sample 1899T and mesenchymal stem cells (MSC). Primers 5-6 target the ecDNA junction. (D) Agarose gel electrophoresis confirming specific amplification of the 304 bp junction product. Lanes 1-4: control amplicons for 18SrRNA (187 bp) and GAPDH (235 bp) in both 1899T tumor and MSC samples. Lane 5: specific amplification of the seg3-seg1 junction in 1899T ecDNA50. Lane 6: no amplification in MSC negative control, confirming the junction is tumor-specific. Molecular weight markers are shown on the left.
